# Molecular insight into how the position of an abasic site and its sequence environment influence DNA duplex stability and dynamics

**DOI:** 10.1101/2023.07.22.550182

**Authors:** Brennan Ashwood, Michael S. Jones, Yumin Lee, Joseph R. Sachleben, Andrew L. Ferguson, Andrei Tokmakoff

## Abstract

Local perturbations to DNA base-pairing stability from lesions and chemical modifications can alter the stability and dynamics of an entire oligonucleotide. End effects may cause the position of a disruption within a short duplex to influence duplex stability and structural dynamics, yet this aspect of nucleic acid modifications is often overlooked. We investigate how the position of an abasic site (AP site) impacts the stability and dynamics of short DNA duplexes. Using a combination of steady-state and time-resolved spectroscopy and molecular dynamics simulations, we unravel an interplay between AP-site position and nucleobase sequence that controls energetic and dynamic disruption to the duplex. The duplex is disrupted into two segments by an entropic barrier for base pairing on each side of the AP site. The barrier induces fraying of the short segment when an AP site is near the termini. Shifting the AP site inward promotes a transition from short-segment fraying to fully encompassing the barrier into the thermodynamics of hybridization, leading to further destabilization the duplex. Nucleobase sequence determines the length scale for this transition by tuning the barrier height and base-pair stability of the short segment, and certain sequences enable out-of-register base pairing to minimize the barrier height.

## Introduction

Design of nucleic acid technologies such as microarrays for sequencing and gene expression profiling, DNA-PAINT for imaging, dynamic DNA devices, and CRISPR-Cas systems for gene editing rely on a predictive understanding of duplex hybridization and base pairing in oligonucleotides.(1-4) Most important for these areas is a fundamental understanding of hybridization thermodynamics, kinetics, and dynamics as a function of nucleobase sequence and for non-canonical base-pairing interactions. Although molecular interpretations for sequence-dependent duplex stability are still developing,(5,6) nearest-neighbor (NN) models provide empirical yet quantitative prediction of duplex binding free energy,(7,8) and efforts have been undertaken to predict hybridization kinetics.(9-12) Numerous types of nucleobase chemical modifications and lesions are also known to influence the stability of base pairing,(13-15) and these effects are often quantified as a function of local nucleobase sequence around the modified site in NN models.(16-18) Additionally, end effects are likely to cause the position of a modification within a short oligonucleotide to influence the binding stability. Base-pair mismatches have been reported to be more destabilizing when located progressively inward from the duplex termini,(19-22) and this behavior may be general for any destabilizing modification. Such position dependence likely results from an interplay of position-dependent entropic penalties and disruption of local base-pairing dynamics, but details of the underlying molecular mechanisms at play are not yet resolved.

Formation of an abasic site (apurinic/apyrimidinic or AP site) results from the loss of a nucleobase through spontaneous or enzymatic cleavage of the glycosidic bond and is one of the most naturally abundant DNA lesions.(23,24) Relative to mismatches, which may exhibit an array of non-canonical base-pairing configurations and structural deformations depending on their sequence context,(25-27) AP sites exhibit minimal perturbation to DNA duplex structure.(28,29) Therefore, AP sites can be used to probe positional effects on duplex stability and base-pairing dynamics with minimal complexity added from non-canonical base pairing interactions. Previous work showed that duplex destabilization arising from an AP site at the center of an oligonucleotide strongly depends on the identity of the adjacent base pairs.(30-32) We recently demonstrated that the degree of destabilization from a central AP site depends on the full oligonucleotide sequence in addition to the adjacent base pairs, a consequence arising from a sequence-dependent free-energy penalty for nucleating base pairs on each side of the AP site.(33) However, it remains unclear how the position of the AP site will tune this penalty, the overall duplex destabilization, and the dynamics of base pairing in the oligonucleotide.

We investigate the impact of shifting the AP-site position on the disruption of base-pairing stability and dynamics in multiple 11-mer template oligonucleotides with variable arrangement of guanine:cytosine (G:C) and adenine:thymine (A:T) base pairs. Temperature-dependent infrared (IR) and ^1^H NMR spectroscopy demonstrate that AP sites increasingly disfavor duplex hybridization as they move inward from the terminus. A cooperative helix-coil model and molecular dynamics (MD) simulations employing the 3-site-per-nucleotide (3SPN.2) coarse-grained model(34,35) reveal that the position-dependent destabilization stems from a nucleation penalty for base pairing on each side of the AP site, which leads to a transition from multi-base-pair fraying when an AP site is near the termini to metastable half-dehybridization with central AP sites. The presence of these metastable partially-dehybridized duplex segments are directly resolved through temperature-jump IR spectroscopy (T-jump IR). Sequence-specific effects complicate the generality of these observations. For instance, ^1^H NMR spectroscopy and all-atom MD simulations indicate that certain segment sequences may circumvent this nucleation penalty by forming out-of-register base pairs. Experimental results can consistently be interpreted with 3SPN.2 MD simulations to provide detailed insight into the mechanism by which sequence and AP-site position alter the free-energy landscape for duplex hybridization.

## Materials and Methods

### Oligonucleotide preparation

Unmodified and modified DNA oligonucleotides were purchased from Integrated DNA Technologies (IDT) at desalt-grade purity. AP sites were incorporated as a tetrahydrofuran group (dSpacer). Samples were further desalted using 3 kDa centrifugal filters (Amicon) or dialyzed in ∼1.5 L ultrapure water for 24 h using Slide-A-Lyzer cassettes (2kDa cutoff, Thermo Scientific). Oligonucleotides were prepared in pH* 6.8 400 mM sodium phosphate buffer (SPB, [Na^+^] = 600 mM) for all measurements. Samples were prepared in deuterated SPB, lyophilized to dryness, and re-dissolved in D_2_O for FTIR and T-jump IR spectroscopy to avoid spectral interference with the H_2_O bending vibrational band. Oligonucleotide concentration was verified with UV absorbance using a NanoDrop UV/Vis spectrometer (Thermo Scientific).

### Temperature-dependent FTIR spectroscopy

FTIR temperature series were collected using the same method as previously reported.(36) FTIR spectra were acquired using a Bruker Tensor FTIR spectrometer at 2 cm^-1^ resolution. Samples were held between two 1 mm CaF_2_ windows with a pathlength set by a 50 µm spacer. All measurements were performed with a 1:1 ratio of complementary strands and a total oligonucleotide concentration of 2 mM. To ensure oligonucleotides start in their minimum energy conformation, the solution of complementary strands was placed in a water bath at 90 °C for 3 min and cooled to room temperature under ambient conditions prior to each measurement. The temperature was ramped with a ∼2.6 °C step size and equilibrated for 3 min at each step. A discrete wavelet transform using the Mallat algorithm and symlet family was applied to the 1490 – 1750 cm^-1^ region of the FTIR spectra to separate and subtract the D_2_O background absorption from the data.(37,38)

### Two-dimensional IR spectroscopy

Two-dimensional infrared spectroscopy (2D IR) temperature series were acquired using a previously described setup employing a pump-probe beam geometry.(39) Spectra were collected using parallel pulse polarization (ZZZZ) at a fixed waiting time (*t*_2_) of 150 fs. The pump pulse pair delay (*t*_1_) was scanned from -160 to 1900 fs in 16 fs steps. To monitor the full range of ring and carbonyl vibrations, the detection frequency axis was measured with approximately 6.2 cm^-1^ resolution. Samples were prepared identically as for FTIR measurements. Sample temperature was controlled using a recirculating chiller (Ministat 125, Huber). Temperature series were performed with a ∼2.8 °C step size and equilibrated for 4 min at each step.

### Temperature-jump IR spectroscopy

We have previously described the temperature-jump (T-jump) IR spectrometer used in this work.(40,41) Measurements were performed using the same sample conditions and sample cell as for FTIR measurements. The T-jump magnitude (∆*T* = *T*_*f*_ − *T*_*i*_) was set to ∼15 °C for all measurements as determined from the change in mid-IR solvent transmission following heating by the near-IR T-jump pulse. The T-jump induced change in mid-IR vibrations of DNA are probed using heterodyned dispersed vibrational echo spectroscopy (HDVE)(42) with parallel (ZZZZ) pulse polarization and fixed at a waiting time (*t*_2_) of 150 fs. The real part of the t-HDVE spectrum is reported and contains similar information to an IR pump-probe spectrum.

### ^1^H NMR spectroscopy

^1^H NMR measurements were performed on a Bruker AVANCE III 600 MHz spectrometer equipped with a Bruker TXI probe. Nuclear Overhauser effect spectroscopy (NOESY) experiments were performed with a mixing time (*t*_*mix*_) of 200 ms unless noted otherwise. Measurements were performed in both 95% H_2_O/5% D_2_O and fully deuterated pH* 6.8 400 mM SPB solutions. 2048 and 1900 complex points were acquired in detection (*t*_2_) and evolution (*t*_1_) delays, respectively, over spectral widths of 24 ppm for 95% H_2_O samples and 12 ppm for fully deuterated solutions. Solutions contained ∼1 mM 3-(Trimethylsilyl)propionic-2,2,3,3-d_4_ acid (TMSP, Sigma-Aldrich) as a frequency reference.

Total correlation spectroscopy (TOCSY) temperature series measurements were measured with DIPSI II isotropic mixing and *t*_*mix*_ = 20 ms. The sample was equilibrated for 5 min and auto gradient and lock shimmed with TopShim at each temperature prior to acquiring spectra. 2048 and 512 complex points were acquired in *t*_2_ and *t*_1_, respectively, over spectral widths of 12 ppm.

### Coarse-grained molecular dynamics simulations

Molecular dynamics (MD) simulations were performed using the coarse-grained 3-Site-Per-Nucleotide (3SPN.2) model.(34) AP sites are incorporated by removing a nucleobase site as previously reported.(33) All systems were simulated using the LAMMPS package(43) compiled with the 3SPN.2 model plugin.(34) Simulations were performed in the NVT ensemble in a periodic box of side length 8.5 nm, equivalent to an effective 5.3 mM oligomer concentration. Classical mechanical equations of motion were integrated via Langevin dynamics in the scheme developed by Bussi and Parrinello(44) with a 20 fs integration time step. Solvent was treated implicitly with an experimentally informed friction coefficient of 9.94 × 10^-11^ m^2^/s.(34,45) A 600 mM implicit salt concentration was used and electrostatic interactions were treated using the Debye-Hückel with a 5 nm cutoff radius.(46) All simulations were initialized from unmodified (WT) B-DNA duplex configurations generated by the 3SPN.2 software.

### Enhanced sampling of hybridization free-energy landscape

We sampled the hybridization free-energy landscape at various temperatures using well-tempered metadynamics (WTMetaD) via the PLUMED plugin,(47,48) as described previously.(33) For each system, we selected a collective variable (CV) that described the average distance between the beads representing native Watson-Crick-Franklin (WCF) base pairs (*r*_*bp*_). The CV was calculated by averaging over all available in-register pair distances – 11 for WT or 10 for AP sequences. Statistics were accumulated over the WTMetaD runs within the quasi-static regime where the applied bias was converged(49) and was analytically reweighted to report unbiased thermodynamic averages and free-energy landscapes in various order parameters beyond those in which sampling was conducted.

### Markov state model temperature-jump simulations

Markov state models (MSMs)(50,51) were constructed from 25 independent and unbiased 10 µs 3SPN.2 MD simulations of each WT and AP system. MD simulations were performed near the sequence-dependent melting temperature calculated from the WTMetaD simulations (*T*_*m,MD*_). Initial velocities were assigned from a Maxwell-Boltzmann distribution at *T*_*m,MD*_. Each simulation was conducted for 12 μs and frames saved to disc every 100 ps, resulting in 1-2 hybridization and dehybridization events per trajectory. The first 2 μs of each run was discarded to produce 25 × 10 μs = 250 μs (2.5 million frames) of simulation data for each sequence.

An additional 25 μs MD simulation was performed for each system 15 °C below *T*_*m,MD*_, and the data was projected into the microstates generated at *T*_*m,MD*_ and used as the initial distribution for relaxation simulation with the MSM built at *T*_*m,MD*_. Each microstate was scored in terms of the average number of intact A:T and G:C base pairs. Base pairs were assigned using a radial cutoff of 0.85 nm for G:C and 0.90 nm for A:T, and each nucleobase was restricted to one base pair at a time. Further details of MSM T-jump simulations were reported previously.(33)

### All-atom molecular dynamics simulations

CGCcap-AP2 and CCend-AP2 DNA duplex topologies were constructed from an intact template of each sequence using AMBERTools22.(52) Only the first 5 base pairs of each sequence were used for all-atom simulation (CGCAT for CGCcap-AP2 and CCTAT CCend-AP2) to minimize unnecessary computational cost for the MD simulations. As described below, the last three residues of each 5-mer were constrained to their native state to reduce fraying and simulate the presence of a longer stabilizing duplex. The AP2 base atoms were removed and replaced with a hydrogen atom, and a small excess charge was redistributed to the 1′ hydroxyl. The parmbsc1 force field was used with TIP3 water.(53,54) A cubic simulation box with periodic boundaries and 4.58 nm side length was used to maintain a 1 nm spacing between the DNA and box boundaries. NaCl ions were added to the box to maintain a 600 mM ionic concentration and to balance the negative charge from the 8 phosphate backbone groups.

Energy minimization was performed with the steepest descent algorithm to ensure a maximum force below 1000 kJ mol^-1^ nm^-1^. Consecutive equilibration simulations were performed in the NVT and NPT ensembles for 100 ps each. Production runs were performed for 1 µs in the NVT ensemble. Simulations were propagated with a 2 fs time step using a leap-frog integrator,(55) and frames were saved every 200 ps. The simulation temperature was maintained at 300 K using a velocity-rescaling thermostat.(56) The LINCS algorithm was used to constrain hydrogen bonds,(57) and a Particle Mesh Ewald was used to calculate long-range electrostatic interactions.(58) The E root-mean-square deviation (eRMSD)(59) of the last three base pairs was calculated using PLUMED v2.8.(47,60) eRMSD compares structures using only the relative distance and orientation of nucleobases, whereas the full structure RMSD often fails to differentiate between nucleic acid conformations with distinct base-pairing configurations. To ensure the last three base pairs retained their native configuration, the eRMSD was restrained by setting an upper wall at 0.6. For each sequence, 20 independent simulations were performed from the same initial state.

Trajectory analysis was performed using the MDTraj and PyEMMA Python libraries.(61,62) Average hydrogen bond distances were calculated between the three hydrogen bonding atoms on the 5′ terminal C1 (N4H2, N3, O2) and corresponding atoms (O6, H1, N2H1) on the 3′ terminal G1. For the CCend-AP2 sequence, an out-of-register hydrogen bond distance was calculated between the same set of C1 atoms and the corresponding set on the internal G2. This second pairing represents a C:C mismatch for CGCcap-AP2, therefore an analogous distance was calculated using the average distance of the available hydrogen bonding atoms on each residue.

## Results & Discussion

### Position-dependent destabilization of DNA duplex by an abasic site

#### Duplex destabilization depends on AP-site position

An AP site destabilizes the duplex, and the magnitude of destabilization depends on the position of the AP site. We compare the thermodynamic impact of incorporating an AP site at the second (AP2), fourth (AP4), and central sixth (AP6) position relative to the unmodified form (WT) of a homogenous sequence 5ʹ-TTTTTTTTTTT-3ʹ + complement (T11). Duplex melting curves extracted from a two-state thermodynamic analysis of FTIR temperature series show that the melting transition midpoint (*T*_*m*_) decreases by 5 to 20 °C as the AP site is shifted from the second to sixth position (Figs. 1a & S1-S3). The change in the hybridization free energy between WT and AP sequences 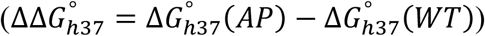 increases linearly in 6 kJ/mol increments from AP2 to AP4 to AP6 (Fig. 1b). The change in hybridization van’t Hoff enthalpy 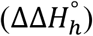 and entropy 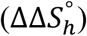 also increase by ∼45 kJ/mol and ∼120 J/molK over this range, respectively. After correcting to an equivalent temperature of 25 °C, isothermal titration calorimetry (ITC, Section S1.3) gives similar 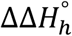 values to FTIR for T11-AP2 and T11-AP4 but a smaller value for T11-AP6.

**Figure 1.**
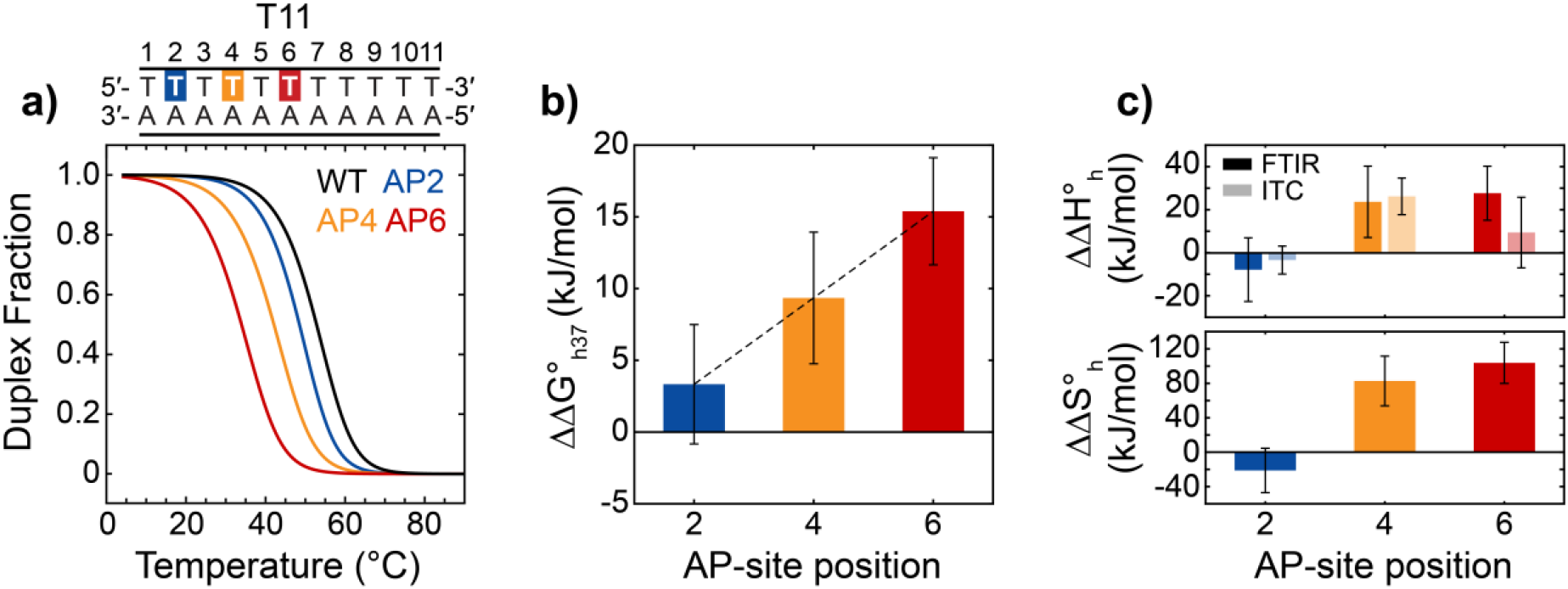
DNA duplex destabilization depends on AP site position. **(a)** Duplex melting curves for a homogeneous A:T sequence that is unmodified (WT, black) or contains an AP site at the 2 (AP2), 4 (AP4), or 6 (AP6) position. Melting curves were extracted from a two-state analysis of FTIR temperature series (Section S1.2). **(b)** Change in hybridization free energy at 37 °C for each AP sequence with respect to the WT sequence, 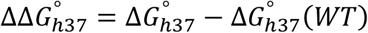. The dashed line corresponds to a linear fit with a slope of 3 kJ/mol per base-pair index. **(c)** Change in hybridization enthalpy 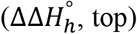 and hybridization entropy 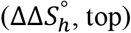 with respect to the WT sequence. Light 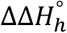 bars correspond to values measured with ITC in non-deuterated solution (Section S1.3). FTIR and ITC 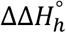 values were shifted to 25 °C using the change in heat capacity (∆*C*_*p*_) measured for T11-WT with ITC. FTIR and ITC error bars correspond to 95% confidence intervals from two-state fits. An AP site is least destabilizing near the duplex termini and becomes increasingly destabilizing when moved inward.

#### Position-dependent destabilization arises from segment nucleation barrier

The position-dependent destabilization from an AP site may be understood through a statistical treatment of duplex hybridization. Helix-coil (HC) models provide one of the simplest statistical descriptions of cooperative DNA duplex melting thermodynamics.(63,64) Base-pairing thermodynamics are described by two equilibrium constants: *s*, which is the Boltzmann factor of the free energy for adding an individual base pair next to an intact base pair 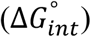 and *σ*, which is the Boltzmann factor of the free energy for nucleating a stretch of base pairs 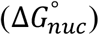.

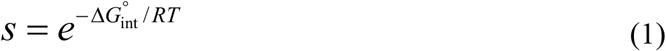

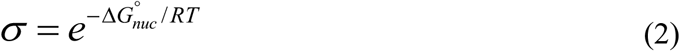

Since this is a mean-field model, we apply the average value of *s* across all possible contacts within the duplex, ⟨*s*⟩, as determined from nearest-neighbor (NN) enthalpy 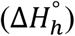 and 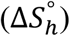 parameters.(7) The large penalty to nucleation makes *σ* << 1, which is the source of cooperativity in the model. We assume 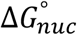 is purely entropic such that *σ* is independent of temperature.

It is generally unfavorable to have multiple disconnected stretches of base pairs in short duplexes, so we describe the WT system with an internal partition function (*Z*_*int,D*_) limited to a single stretch of base pairs.(63-65)

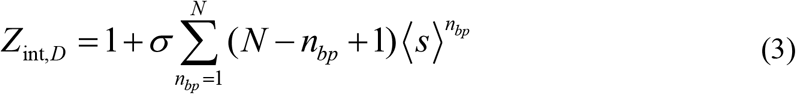

*N* is the maximum number of possible base pairs, and *n*_*bp*_ is the number of intact base pairs for a given microstate. As detailed in Section S2, we can extend the HC model to treat an AP site as a defect that splits the duplex into two stretches of *N*_1_ and *N*_2_ base pairs (Fig. 2a). The internal partition function for the duplex can then be written as a product of two partition functions similar to eq. 3 for the two stretches, *Z*_*int,D*_ = *Z*_*int,D*1_*Z*_*int,D*2_. Each stretch has its own nucleation penalty (*σ*_1_, *σ*_2_) and ⟨*s*⟩ value, which corresponds to the average base-pair equilibrium constant across the respective stretch. Our usage of a second nucleation penalty is similar to incorporating a ‘defect’ free-energy penalty as in previous statistical models of duplexes containing a mismatch.(19)

**Figure 2.**
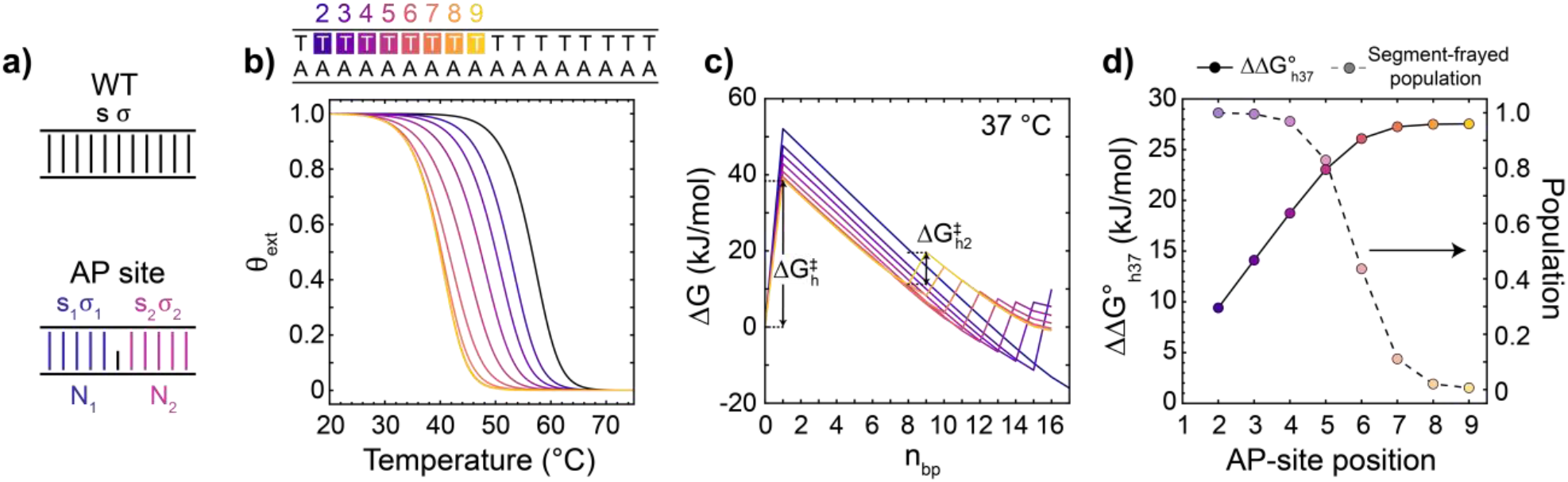
Helix-coil modeling of position-dependent duplex destabilization from an AP site. **(a)** Schematic of two-stretch helix-coil (HC) model used to model the impact of AP-site position on duplex stability. The WT duplex contains a single nucleation penalty (*σ*) and base-pair stability constant (*s*) applied uniformly to each base-pair site.(63,64) An AP site splits the duplex into two base-pair stretches with their own *s* and s values. **(b)** Fraction of duplexes containing one or more base pairs (*θ*_*ext*_) for an unmodified T17:A17 sequence (black) and those containing an AP site at the 2 – 9 positions (purple to yellow). *σ*_1_ = *σ*_2_ = 10^-4^ for all sequences. **(c)** Free-energy profiles at 37 °C for each sequence, illustrating barriers for strand association 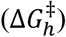 and nucleation of the second-segment 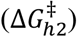. **(d)** 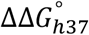 (dark circles and solid line) and the corresponding fraction of duplexes with the short base-pair segment completely frayed (light circles and dashed line). Shifting the AP site inward from the terminus leads to an increase in the number of frayed base pairs and greater destabilization of the duplex. The degree of destabilization levels off once the binding stability of the short segment is large enough to overcome the nucleation penalty.

To calculate melting curves from the HC model, we determine the fraction of oligonucleotide strands that contain one or more base pairs (*θ*_*ext*_) as a function of temperature. *θ*_*ext*_ depends on both external and internal degrees of freedom of the system. We evaluate the external partition functions of the single-strand and duplex states as the number of possible ways of arranging single-strand and duplex molecules as a self-avoiding walk on a 3D cubic lattice,(66) allowing us to compute *θ*_*ext*_ in terms of *Z*_*int,D*_.

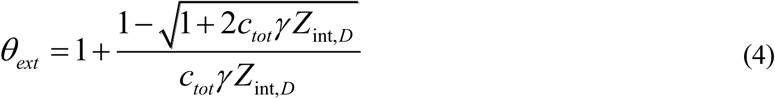

where *γ* = 6*c*°N_*A*_*V*_*ss*_

*c*° is the standard-state concentration of 1 M, *N*_*A*_ is Avogadro’s constant, *V*_*ss*_ is the volume of a single-strand molecule, and *c*_*tot*_ is the total concentration of oligonucleotides. We neglect the temperature-dependence of *V*_*ss*_ such that the temperature-dependence of *θ*_*ext*_ comes only from *Z*_*int,D*_.

Figure 2b shows calculated *θ*_*ext*_ melting curves for an extended homogeneous sequence, T17:A17. Moving an AP site from the second to sixth base pair site leads to a gradual reduction in the duplex melting temperature, consistent with the experimental observation for T11. The shift in melting temperature becomes minor and eventually levels off as the AP site moves closer to the duplex center. This trend in the *θ*_*ext*_ curves may be understood from examining the free-energy profile (FEP) as a function on the number of intact base pairs (*n*_*bp*_) calculated from the HC model (Fig. 2c). The WT sequence exhibits a single free-energy barrier to hybridization 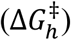 between the single-strand (*n*_*bp*_ = 0) and duplex states that arises from the reduction in translation and configurational (set by *σ*) entropy upon binding of single strands, and formation of the remaining base pairs is cooperative and downhill in energy. In the HC model, the AP site disrupts this cooperativity and introduces an additional entropic barrier for forming base pairs on each side of the AP site 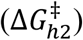, and we recently experimentally and computationally verified the presence of 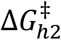in duplexes with a central AP site.(33) The AP-site position controls the position of 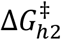 along the hybridization FEP. When an AP site is at the second base-pair position, the adjacent terminal base pair does not have enough binding stability to overcome 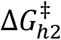 and remains highly frayed at 37 °C. As the AP site shifts from the second to sixth base-pair position, the shorter duplex segment increases in length and base-pairing stability but still remains highly frayed (Fig. 2c). The number of frayed base pairs increases with segment length, leading to the position-dependent decrease in duplex stability observed from the *θ*_*ext*_ curves and 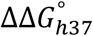. Once the short segment reaches a length of seven base pairs, the binding stability overcomes 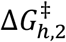 and duplex destabilization levels off. 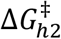 is fully encompassed into the duplex state at these AP-site positions, creating a local free-energy minimum along *n*_*bp*_ that corresponds to configurations with one segment dehybridized. The levelling off is also facilitated by a reduction in the magnitude of 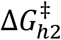 from 14 to 9 kJ/mol as the AP site moves inward from the termini.

The HC calculations suggest that the position-dependent destabilization from an AP site depends only on the magnitude of the nucleation barrier and the binding stability of the short segment. An additional minor effect is that the number of base-pair arrangements possible over the whole duplex decreases as the AP site moves closer to the duplex, resulting in a reduction of base-pairing combinational entropy and a free-energy penalty of 1 – 2 kJ/mol (Fig. S6). Given the small magnitude of this effect, duplex destabilization is essentially only position-dependent when an AP site is close to the termini. This range is roughly ∼1 – 9 base pairs from our model calculations but depends on the molecular properties of 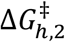 and the short duplex segment.

#### Interplay of nucleobase sequence and AP-site position in duplex destabilization

Beyond the factors that influence a homogeneous duplex, destabilization by an AP site will depend on multiple aspects of the nucleobase sequence. Previous reports showed that 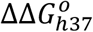 is highly sensitive to the identity of the nucleobase being removed as well as the bases adjacent to the AP site.(31,32) Further, nucleobase sequence will influence the binding stability of the short duplex segment as well as the magnitude of 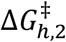 . We use three additional 11-mers with distinct base-pairing properties to investigate sequence-dependent destabilization from an AP site (Fig. 3a). “CGCcap” contains three G:C base pairs on one end to create asymmetry in the duplex base-pairing stability, “CCends” places a pair of G:C base pairs at the termini to minimize base-pair fraying, and “GCGcore” has central G:C base pairs to promote fraying of the A:T termini.(36,67)

**Figure 3.**
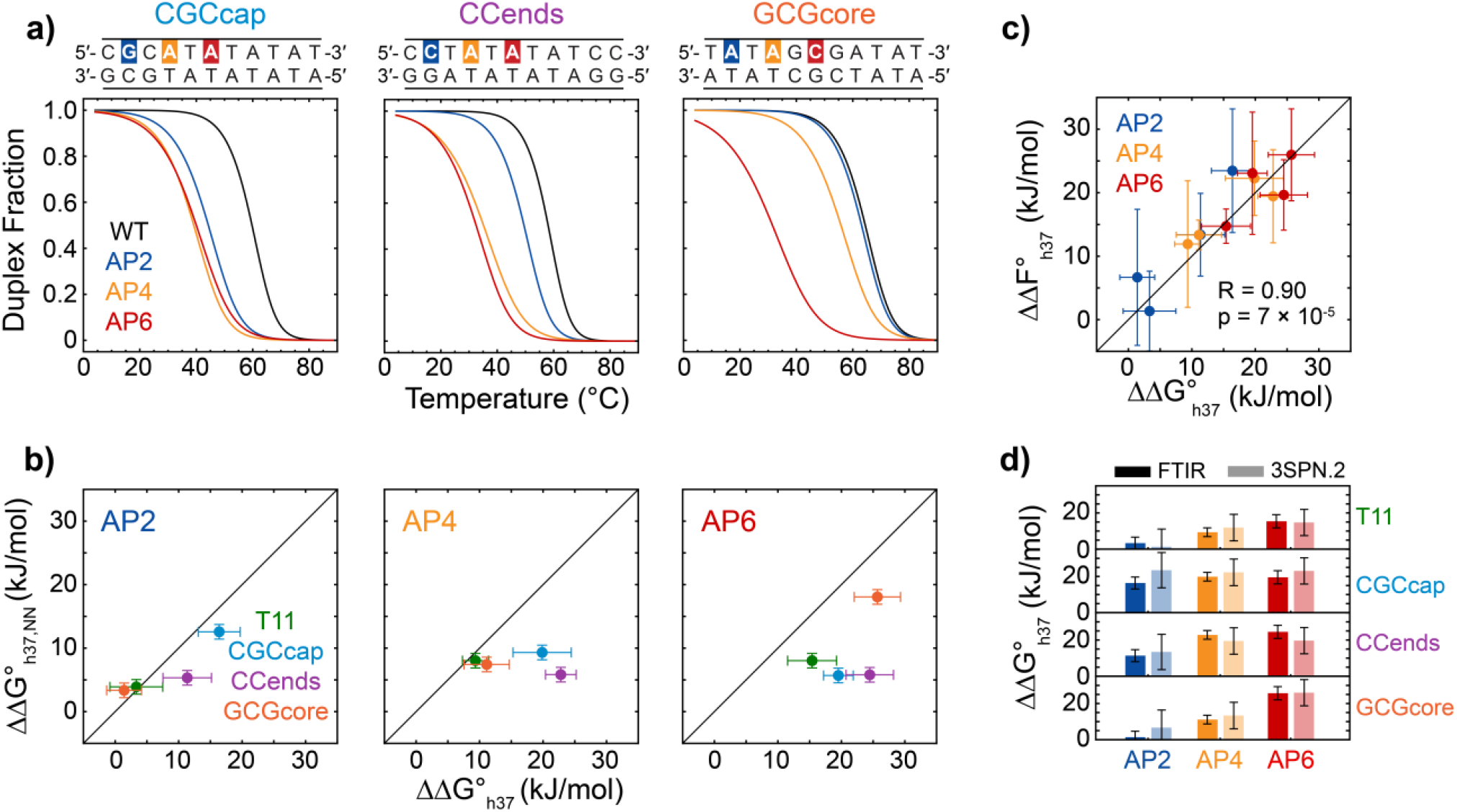
Sequence-dependent duplex destabilization from an AP site. **(a)** Duplex melting curves for CGCcap, CCends, and GCGcore sequences determined from a two-state analysis of FTIR temperature series. **(b)** Scatter plots of 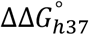 from FTIR vs. nearest-neighbor (NN) model calculations of duplex free-energy change from an AP site 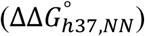 for all sequences. Plots are separated by AP-site position. The comparison indicates that the NN model systematically underestimates 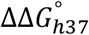 and poorly predicts sequence-dependent destabilization from an AP site. **(c)** Scatter plot of 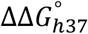 from FTIR vs. 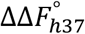 from 3SPN.2-determined melting curves for all sequences with an AP site. The Pearson correlation coefficient (R) and p value is listed. **(d)** Comparison of 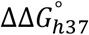 (dark bars) and 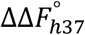 (light bars) for each sequence. Error bars for 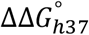 and 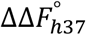 are propagated from two-state fits, and NN error bars correspond to the reported standard deviation from comparison with experimental data.(7)

An AP site is generally more destabilizing when shifted inward from the termini, but Fig. 3a illustrates how the relative placement of G:C and A:T base pairs significantly tunes this position-dependent trend. At identical position, incorporating an AP site at a G:C base pair is more destabilizing than at an A:T base pair. However, the removal of an interior A:T base pair can be equivalent or even more destabilizing than removing a near-terminal G:C base pair. The strength of local base-pairing and stacking interactions around the AP site decreases from AP2 to AP4 to AP6 modifications for CGCcap, which effectively cancels out the position-dependent effects and leads to a similar destabilization for all AP-site positions. Further, incorporating an AP site at the central A:T site of CCends leads to twice the magnitude of 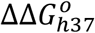 relative to removal of the guanine at the second site.

The AP-site position and its local sequence context alone do not fully capture the sequence-dependent trends in 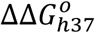. This point is illustrated in Fig. 3b by comparing 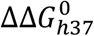 values from FTIR and those calculated with Santa Lucia’s NN model 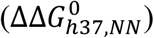.(7) 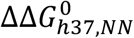 values for AP4 and AP6 sequences are calculated by removing the two NN parameters associated with the AP site. For AP2 sequences, the 3ʹ NN parameter is removed and the 5ʹ parameter is halved to account for having just a single base-pair and stacking interaction at the terminal base. The NN model systematically underestimates the magnitude of 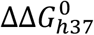 due to the lack of a free-energy penalty for base pairing on each side of the AP site. Further, NN effects do not predict the sequence-dependence of 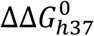at fixed AP-site position. For example, 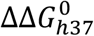 spans 14 kJ/mol for AP4 sequences yet 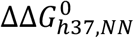 values for these sequences fall within a ∼3 kJ/mol window. In another example, CCends-AP6 and GCGcore-AP6 exhibit 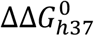 values within 1 kJ/mol and their 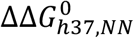 predictions are separated by 12 kJ/mol. These observations suggest the presence of additional sequence-dependent contributions to the free-energy penalty from an AP site.

#### Sequence-dependent nucleation barrier and short-segment stability

We next aim to evaluate how the properties of the segment-nucleation barrier and binding stability of the short segment depend on nucleobase sequence. The HC model qualitatively captures the effect of sequence on duplex destabilization (Figs. S6 – S7), but it neglects numerous molecular factors that are necessary for an accurate description of the underlying base-pairing free-energy landscape. Therefore, we next characterized the underlying hybridization free-energy landscape as a function of AP-site position using coarse-grained MD simulations. In particular, the threshold segment length for stable hybridization, its sequence-dependence, and the molecular behavior of few base-pair segments were examined.

#### Nucleation barrier and short-segment stability from coarse-grained MD simulations

We conducted MD simulations with the 3SPN.2 coarse-grained model employing well-tempered metadynamics (WTMetaD) to sample the hybridization free-energy landscape at 7 – 8 temperatures from *T*_*m,MD*_ − 20 °C to *T*_*m,MD*_ + 20 °C, where *T*_*m,MD*_ is the melting temperature determined with 3SPN.2. Incorporation of an AP site in 3SPN.2 has negligible impact on the B-DNA duplex structure, and the free nucleobase remains predominantly intrahelical (Fig. S8),(33) consistent with previous structural characterization of duplexes containing AP sites.(28) Two-state analysis of 3SPN.2-determined melting curves produces hybridization Helmholtz free energies (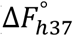 and 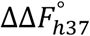) that are well-correlated (R = 0.90) with experimental values (Figs. 3cd & S9), indicating that 3SPN.2 reasonably predicts the sequence-dependent free-energy penalty from an AP site. In contrast, 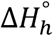 and 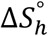 are poorly captured by 3SPN.2, particularly for AP sequences (Figs. S9).

Figure 4 shows FEPs computed from the probability distribution along *n*_*bp*_ for CGCcap, CCends, and GCGcore sequences that qualitatively agree with those from the HC model (Figs. 2c & S7). WT sequences show a single free-energy barrier to hybridization 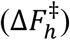 peaked at *n*_*bp*_ = 2 or 3 and hybridization is energetically downhill for *n*_*bp*_ > 3. AP sequences must overcome a second barrier 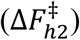 to form a fully intact duplex. Just as for 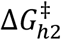 from the HC model, 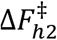 arises from the energy penalty for nucleating the second base-pair segment and creates a local free-energy minimum that corresponds to duplex configurations containing intact base pairs on only one side of the AP site (Figs. S10 and S11). 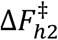 lies between *n*_*bp*_ = 9 and *n*_*bp*_ = 10 in AP2 sequences and is not observable along the *n*_*bp*_ coordinate, yet a barrier for forming the terminal base pair is observed as a function of average base-pair separation (*r*_*bp*_, Figs. S12-S15). Both 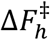 and 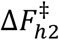 arise from entropic penalties for nucleation and are partially balanced by favorable enthalpy changes for base pairing (Figs. 4b & S16). 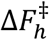 is predominantly a reflection of the reduction in translational and orientational entropy upon bimolecular association, which are absent from 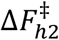 . Instead, 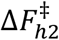 is dominated by a reduction in conformational entropy of the unhybridized segment and is therefore 2-to-5-fold smaller than 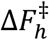 (Fig. S16). 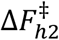 is also sensitive to the sequence of nucleating base pairs adjacent to the AP site (Fig. 4d). 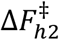 is smallest for GCGcore-AP6, particularly at low temperature, and involves formation of a G:C base pair adjacent to the AP site whereas 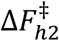 requires formation of one or two A:T base pairs on the second segment in the other sequences, leading to a larger barrier height.

**Figure 4.**
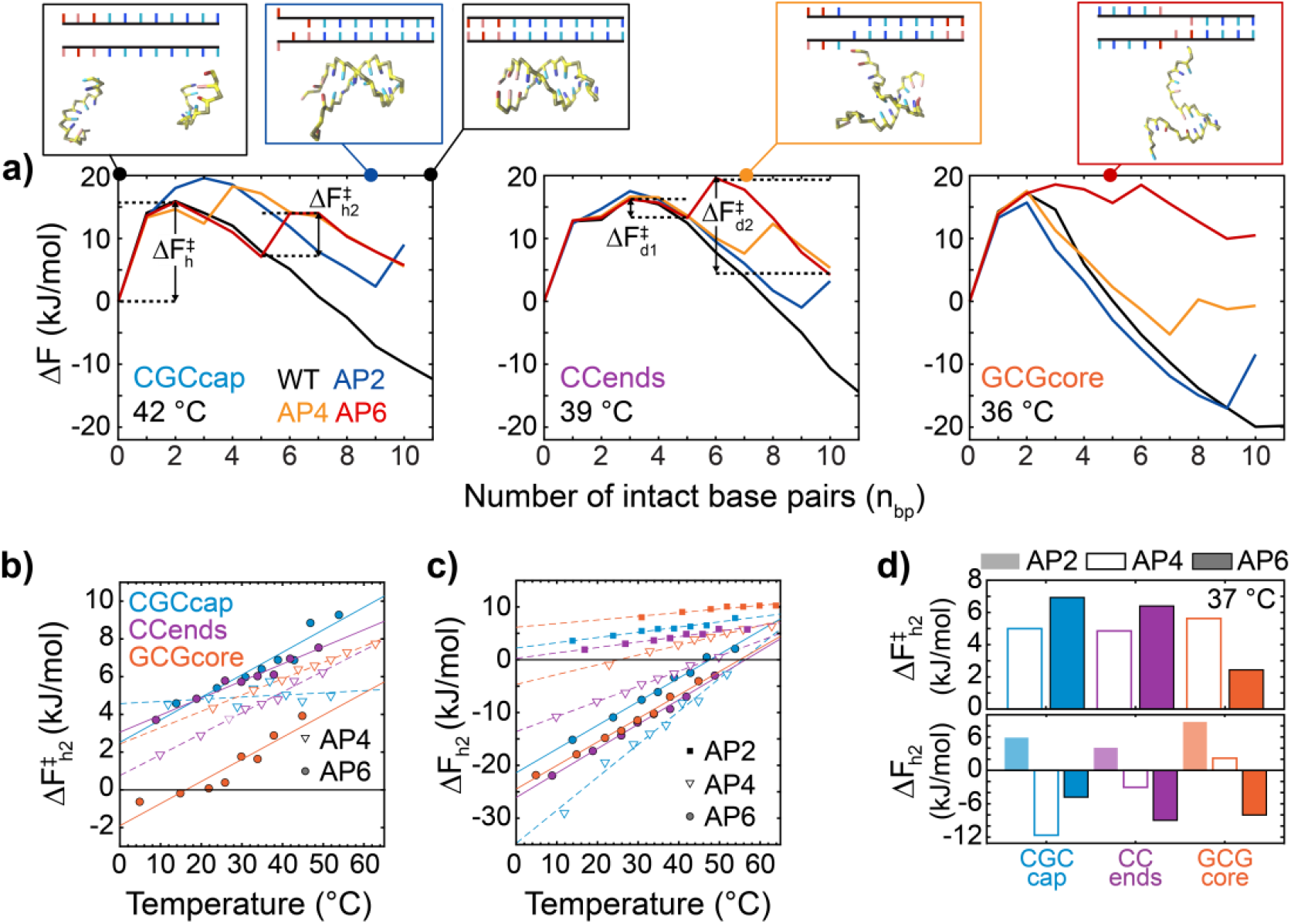
Sequence-dependent free-energy profiles for duplex hybridization. **(a)** Free-energy profiles as a function of the number of intact base pairs, Δ*F*(*n*_*bp*_) = −*RT* In *p*(*n*_*bp*_)/*p*(0), for each sequence from 3SPN.2 MD simulations employing WTMetaD. Base pairs were assigned using a 0.7 nm radial separation cutoff. Simulations were carried out at temperatures 10-20 °C below the 3SPN.2-determind *T*_*m*_ (*T*_*m,MD*_). AP2, AP4, and AP6 sequences show an additional hybridization barrier for *n*_*bp*_ **>** 2 for base pairing on both sides of the AP site 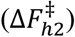. **(b)** 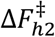 for AP4 (triangles) and AP6 (circles) sequences as a function of temperature. **(c)** Free-energy difference between states with weak-segment frayed and intact, Δ*F*_*h*2_ =−*RT* In *p*(*n* > *n*_*sd*_)/*p*(*n*_*sd*_), for all sequences where *n*_*sd*_ is the number of base-pairs in the segment-dehybridized state. Dark-solid, dashed, and light-solid lines correspond to linear fits for AP6, AP4, and AP2 sequences, respectively. Extracted internal-energy and entropy changes are provided in Fig. S16. **(d)** (top) 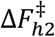 and (bottom) Δ*F*_*h*2_ at 37 °C for each sequence. 3SPN.2 MD simulations suggest that two to five base pairs, depending on the G:C content, provide enough stability for a segment to hybridize.

The AP-site position and nucleobase sequence determine the position and magnitude of the 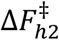 barrier along *n*_*bp*_ and the base-pairing stability of the second segment. The combination of these factors determine the free energy for hybridization of the second segment (Δ*F*_*h*2_). The terminal base pair adjacent to the AP-site in AP2 sequences has an insufficiently stabilizing potential energy for base pairing to overcome the unfavorable entropy, leading to a positive free-energy change for forming the last base pair (Δ*F*_*h*2_ > 0, Fig. 4c). The threshold length of the weak segment to overcome the nucleation penalty and reach Δ*F*_*h*2_ < 0 depends on the sequence of the segment. If the weak segment contains multiple G:C base pairs like in CCends-AP4, then three base pairs are sufficient to form a stable segment. In contrast, the pure A:T segment of GCGcore-AP4 is largely frayed at 37 °C and four to six base pairs are needed to form a stable segment as evidenced by CGCcap-AP4 and CGCcap-AP6. This sequence-dependent length scale for forming a stably bound segment contributes to the mismatch between 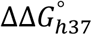 and 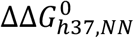 observed in Fig. 3b. For instance, the spread in 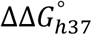 for AP4 sequences may primarily occur because the A:T-rich short segments of T11-AP4 and GCGcore-AP4 are more weakly bound than the short segments of CGCcap-AP4 and CCends-AP4.

#### Observation of sequence-dependent nucleation barrier with T-jump IR spectroscopy

To test for the presence of metastable partially hybridized configurations and a second barrier to hybridization, we turned to T-jump IR spectroscopy, which can be used to directly measure the kinetics of short-segment dehybridization.(33) In these experiments, the sample is equilibrated at a temperature below *T*_*m*_ and optically heated by ∼15 °C (∆*T* = *T*_*f*_ − *T*_*i*_) within 7 ns. The ensuing changes in base pairing of the duplex are monitored with heterodyned dispersed vibrational echo spectroscopy (HDVE)(42) from nanosecond-to-millisecond time delays. HDVE spectroscopy reports on changes in ring and carbonyl vibrational bands similarly to an IR pump-probe spectrum, and we report the spectra as the difference in signal at a given time delay after the T-jump relative to the maximum of the initial-temperature spectrum, Δ*S*(*t*) = [*S*(*t*) − *S*(*T*_*i*_)]/ max[*S*(*T*_*i*_)]. We use the change in excited-state absorptions of the guanine ring mode at 1550 cm^-1^ and of the adenine ring mode at 1605 cm^-1^ as reporters of G:C and A:T base pairing, respectively, as in previous studies.(33,36,67)

T-jump measurements reveal distinct kinetic behavior between WT and AP sequences (Figs. 5a and S17-S21). WT sequences show two dehybridization kinetic components on ∼100 ns (*τ*_1_) and ∼100 µs (*τ*_3_) timescales followed by thermal relaxation back to *T*_*i*_ at ∼1 ms. The T-jump IR data are well described by global lifetime fitting with exponentially damped spectral components, and the same number of components and similar kinetics are obtained from inverse-Laplace-transform rate distribution spectra (Figs. S22-S23). The *τ*_1_ and *τ*_3_ processes were previously assigned to T-jump induced terminal fraying of the duplex and full-strand dehybridization, respectively,(33) consistent with reports for similar sequences.(67) The temperature-dependent trends in timescale and amplitude of *τ*_3_ support its assignment to complete strand dissociation (Figs. S25-S26). Many of the AP4 and AP6 sequences show an additional kinetic component labeled *τ*_2_ between 500 ns to 10 µs with sequence-dependent amplitude and timescale. The *τ*_2_ response corresponds to dehybridization of a complete stretch of base pairs on one side of the AP site (segment dehybridization) and reveals the presence of a free-energy barrier from the fully intact duplex to segment-dehybridized configurations 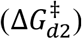. The observation of 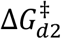 verifies the presence of 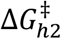 in the reverse direction, as illustrated from the FEPs in Fig. 4a and previously reported for the AP6 sequences.(33)

**Figure 5.**
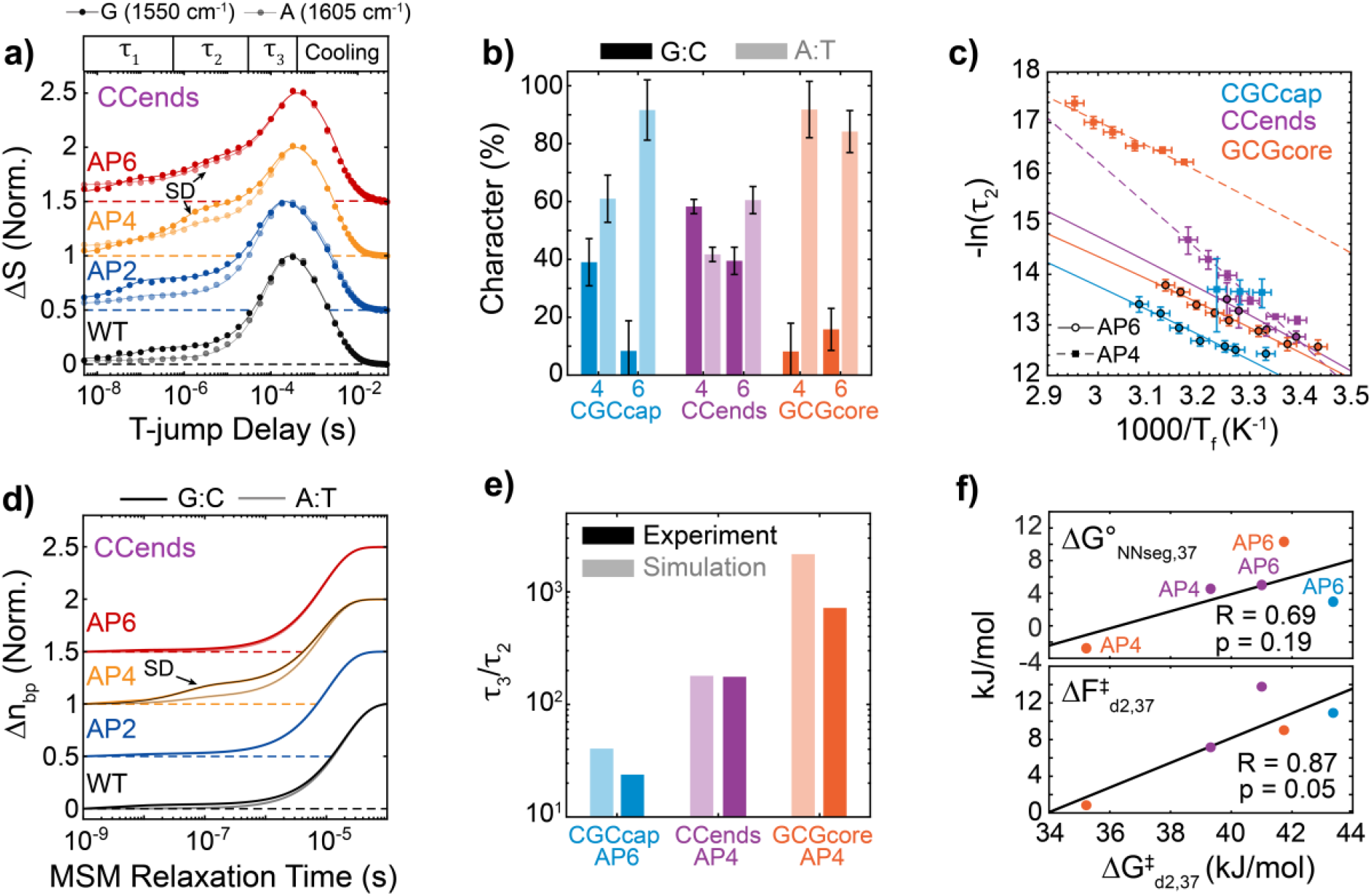
Comparison of segment-dehybridization kinetics observed in experiment and simulation. **(a)** T-jump IR (t-HDVE) time traces plotted for CCends sequences as the normalized signal change relative to the maximum of the initial-temperature spectrum, Δ*S*(*t*) = [*S*(*t*) − *S*(*T*_*i*_)]/max[*S*(*T*_*i*_)]. Δ*S* time traces probed at 1550 cm^-1^ (dark) report on changes in G:C base pairing and those at 1605 cm^-1^ (light) report on changes in A:T base pairing. T-jumps are performed from approximately *T*_*m*_–15°C to *T*_*m*_ for each sequence. Traces are shifted vertically with respect to one another, and dashed lines indicate respective baselines. Solid lines correspond to three- or four-component fits from global lifetime analysis (Section S5.2). A microsecond timescale segment-dehybridization response (SD) is observed for CCends-AP4 and CCends-AP6. **(b)** Percentage of G:C (dark, 1550 cm^-1^) and A:T (light, 1605 cm^-1^) base-pair loss character observed in segment-dehybridization t-HDVE responses. Percentages are derived from amplitudes determined from global lifetime fitting (eqs. S38ab). Error bars indicate 95% confidence intervals propagated from global fits. **(c)** Temperature-dependent observed rate constants for segment-dehybridization (1/*τ*_2_) determined from T-jump IR. Data are fit to a Kramers-like equation in the high-friction limit (eq. S36). The rate trend for CGCcap-AP4 is not fit due to insufficient data. Vertical error bars correspond to 95% confidence intervals from global lifetime fitting and horizontal error bars are the standard deviation in T-jump magnitude. **(d)** Markov state model (MSM) T-jump simulations of the normalized change in intact G:C (dark) and A:T (light) base pairs, Δ*n*_*bp*_(*t*) = [*n*_*bp*_(*t*) − *n*_*bp*_(*T*_*i*_)]/[*n*_*bp*_(*T*_*f*_) − *n*_*bp*_(*T*_*i*_)], from *T*_*m,MD*_–15°C to *T*_*m,MD*_. Traces are shifted vertically with respect to one another, and dashed lines indicate respective baselines. Black/gray solid lines for CCends-AP4 correspond to fits to a sum of two exponential components. **(e)** Ratio of full-strand dissociation time constant (*τ*_3_) to *τ*_2_ from T-jump IR (dark) and MSM T-jump simulations (light) indicating qualitative agreement between simulation and experiment. **(f)** Scatter plots of the segment-dehybridization free-energy barrier determined from T-jump IR 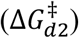 vs. (top) the dehybridization free energy for the respective segment calculated from the NN model (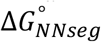, Table S3) and (bottom) 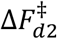 determined from 3SPN.2 MD simulations at 37 °C. Linear fits (solid black line) are shown for each plot with the respective Pearson correlation coefficient (R) and p value. 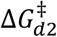 shows better correlation with 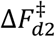 than 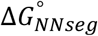 .

The spectral properties and kinetics of the *τ*_2_ process support its assignment to segment-dehybridization and provide insight into the properties of 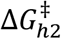 . The fraction of A:T and G:C base-pair loss character during *τ*_2_ determined from global lifetime fitting (Figs. 5b and S27) match the expected signal change for dehybridizing either half of CCends-AP6 and GCGcore-AP6, the 5′-TATAT-3′ segment in CGCcap-AP6, and the 5′-CCT-3′ segment in CCends-AP4. Additionally, the segment dehybridization rate 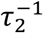 increases exponentially with temperature as expected for an active process, illustrated by the Arrhenius plot in Fig. 5c, which we analyze using a Kramers-like equation (eq. S36). The *τ*_2_ response of CGCcap-AP4 is more difficult to interpret. It has low amplitude and mixed G:C and A:T character even though each segment is purely composed of G:C or A:T base pairs. Due to the similar binding stability of 5′-CGC-3′ and 5′-TATATAT-3′ segments, it is possible that *τ*_2_ contains similar amplitude from both segment-hybridization pathways. Lastly, we observe that GCGcore-AP4 does not have a *τ*_2_ response; however, unlike all other sequences, 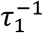 increases exponentially with temperature in an active manner. We therefore assign *τ*_1_ in GCGcore-AP4 to dehybridization of the 5′-TAT-3′ segment and note that it may overlap in time with terminal fraying dynamics from the other end of the duplex.

We find large variation in segment-dehybridization kinetics across sequence and segment length from T-jump IR measurements. To more directly connect these measurements to simulation, we built kinetic models of T-jump relaxation using Markov state models (MSMs) constructed from 250 µs of unbiased 3SPN.2 simulations near *T*_*m,MD*_ for each sequence (Figs. S29-S32) as previously reported for WT and AP6 sequences.(33) The change in the fraction of intact A:T and G:C base pairs (∆*n*_*bp*_) is computed as a function of relaxation time at *T*_*m,MD*_ after initiating the MSM kinetics with a population distribution obtained at *T*_*m,MD*_–15°C to replicate the initial experimental T-jump ensemble. Relaxation traces in Figs. 5d and S17 show two to three kinetic components that correspond to terminal fraying (1 – 30 ns, *τ*_1_), segment-dehybridization (50 – 500 ns, *τ*_2_), and complete strand dissociation (1 – 50 µs, *τ*_3_). The MSM relaxation timescales are accelerated by ∼1 order of magnitude relative to T-jump IR measurements, as previously reported for the 3SPN.2 model,(68) and therefore we use the ratio between *τ*_3_ and *τ*_2_ (or *τ*_1_ for GCGcore-AP4) to compare simulation and experimental timescales. Experimental and simulated *τ*_3_/*τ*_2_ values show qualitative agreement and an increase from CGCcap-AP6 to CCends-AP4 to GCGcore-AP4, suggesting that 3SPN.2 accurately captures sequence-dependent variation in 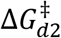 (Fig. 5e). We also find correlation (R = 0.87) between the 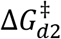 and 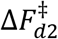 from the FEPs in Fig. 4a at 37 °C (Fig. 5f). The correlation between 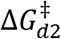 and the calculated NN dehybridization free energy of the respective unbinding segment 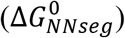 is substantially poorer (R = 0.69), suggesting that both the dehybridization energy of the segment 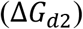 and its nucleation barrier 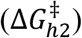 significantly contribute to the sequence-dependence of 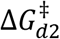.

MSM relaxation profiles generally exhibit a lower segment-dehybridization response amplitude than in experiment, and negligible response is observed for CGCcap-AP4, CCends-AP6, and GCGcore-AP6 even though each sequence possesses a metastable segment-dehybridized state (Figs. 4a, 5d, and S32). For each of these three sequences, we observe 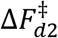 to be larger than the free-energy barrier for dehybridization of the final base-pair segment 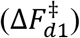 at *T*_*m,MD*_ (Fig. S16). When 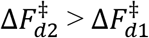, the waiting time to surpass 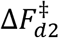 is the rate-limiting step for dehybridization, and we do not expect to observe a segment-dehybridization response in the T-jump kinetics. In contrast, we observe 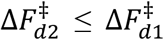 for sequences where there is a large amplitude segment-dehybridizaiton response (CCends-AP4, CGCcap-AP6, GCGcore-AP4). Segment-dehybridization is only observed experimentally for CGCcap-AP4 and CCends-AP6 at the few lowest measured temperatures, which may be explained by 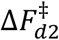 increasing relative to 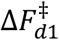 and surpassing it at higher temperatures (Fig. S16). The overall lower amplitude of segment-dehybridization response in simulations therefore suggests that the ratio of 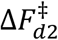 to 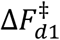 is generally overestimated, and as 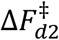 is fairly correlated with 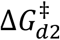, we suggest that sequence-dependent error in this ratio primarily arises from 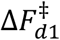 . Error in 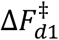 may stem from inaccurate modeling of base-pair and stacking interactions around the AP site for which 3SPN.2 is not parametrized. Further, 3SPN.2 simulations tend to poorly predict sequence-dependent variation in the dehybridization barrier among canonical oligonucleotides.(68)

## Contributions from non-canonical base pairing

### Fraying and out-of-register base pairing at near-terminal AP sites

The T-jump results directly report on 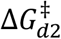 in AP4 and AP6 sequences, but it is still not clear which sequences exhibit fraying of the short segment at equilibrium. For this purpose, we monitor the temperature-dependence of G:C and A:T features in the IR spectrum to check for the presence of segment-dehybridization at temperatures well below *T*_*m*_. The temperature-dependent intensity of the guanine and adenine ring modes are used to report on the fraction of intact G:C and A:T base pairs, respectively. Relative to FTIR, 2D IR temperature series are found to provide greater contrast between spectral changes arising from base-pair disruption and those giving rise to melting curve baselines (Figs. S35 – S37). Except for GCGcore-WT, which shows signatures of A:T terminal fraying, the A:T and G:C temperature profiles of WT sequences are essentially identical and suggest that duplex melting is nearly all-or-nothing (Fig. S37). CGCcap-AP2 shows a clear deviation from all-or-nothing behavior where G:C base-pair character is lost at temperatures 20-30 °C below A:T melting, likely corresponding to disruption of the terminal G:C base pair as predicted from 3SPN.2 MD simulations (Fig. 6a). CCend-AP2, a modified version of CCends with only a single CC terminus, exhibits concerted changes of G:C and A:T signatures and no indication of melting below 1 °C (Figs. S33 – S34), suggesting that the terminal G:C base pair is stabilized relative to CGCcap-AP2.

**Figure 6.**
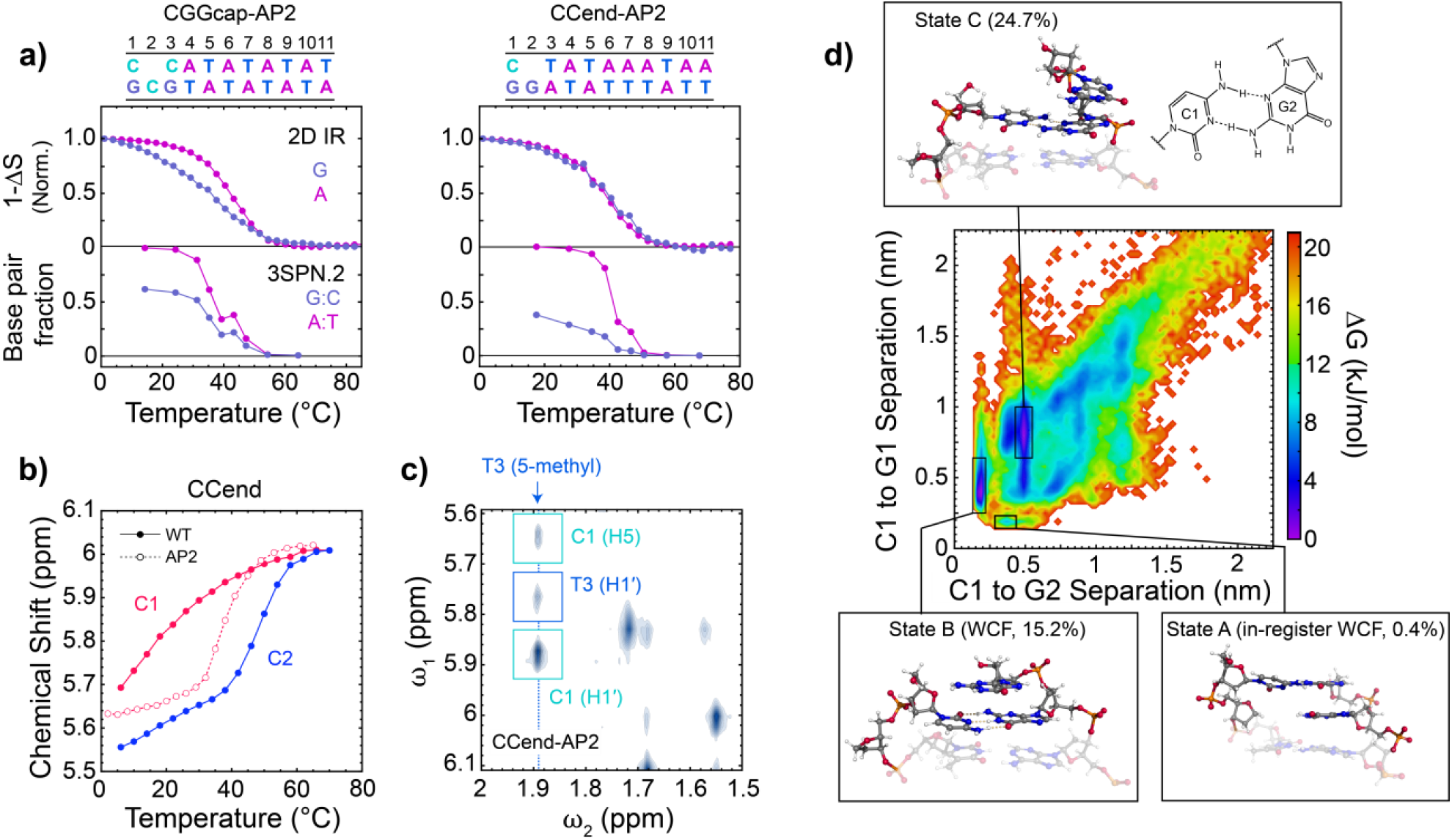
Partial dehybridization and out-of-register base pairing in AP2 duplexes. **(a, top)** Normalized temperature-dependent change in two-dimensional infrared (2D IR) signals relative to the initial temperature, 1 − ∆*S*(*T*) = [*S*(*T*) − *S*(1℃)]/*S*(1℃), at guanine (G) and adenine (A) ring vibrational bands for CGCcap-AP2, and CCend-AP2. Corresponding 2D IR spectra are shown in Fig. S36. **(a, bottom)** Fraction of intact G:C and A:T base pairs from 3SPN.2 MD simulations carried out at 8 – 9 temperatures across the duplex melting transition. Base pairs were assigned using a radial cutoff of 0.7 nm. Low-temperature fraying of the short segment is observed in experiment and simulation for CGCcap-AP2, yet not observed experimentally for CCend-AP2. **(b)** NMR chemical shifts of cytosine H5 nuclei for CCend-WT (filled circles, solid lines) and CCend-AP2 (open circles, dashed lines) determined as a function of temperature from TOCSY measurements. The terminal cytosine (C1) shows low-temperature fraying in CCend-WT whereas the second cytosine (C2) signal primarily changes during duplex dissociation. C1 does not exhibit fraying CCend-AP2 and instead tracks duplex dissociation. **(c)** NOESY cross peaks between H5/H1′ nuclei and 5-methyl protons of thymine for CCend-AP2. Cross-peaks are observed between C1 and T3 (cyan boxes). **(d)** Free-energy surface from 20 × 1 μs all-atom MD simulations of CCend-AP2 using the AMBER-bsc1 force field at 30 °C. The x-axis is the average separation between hydrogen-bonding atoms between C1 and G2 (out-of-register), and the y-axis is the average separation between those atoms in C1 and G1 (in-register). Snapshots of base pairs 1-3 from select free-energy minima are shown, and the population from a given minimum is determined from integrating over the marked rectangles. Additional states are highlighted in Fig. S46. In agreement with experiment, C1 primarily base pairs out-of-register yet contains multiple free-energy minima with distinct base-pairing and stacking configurations.

Comparison of partial-dehybridization signatures across our sequences corroborates thresholds for stable segment hybridization from 3SPN.2 simulations. The identical G:C and A:T melting behavior of CCends-AP4 indicates that the three base-pair segment, 5’-CCT-3’, remains hybridized with the rest of the duplex (Fig. S37). In contrast, the 5’-TAT-3’ segment of GCGcore-AP4 unbinds at temperatures well below the rest of the duplex. These observations support the sequence-dependent trends in Δ*F*_*h*2_ from 3SPN.2 MD simulations and confirm that the length threshold for stable binding (Δ*G*_*h*2_ < 0) of pure A:T segments is in the range of 4 – 7 base pairs at physiological temperatures while that for G:C segments is as low as 2 – 3 base pairs. The nucleation free-energy penalty for the second duplex segment is significantly lower than for the initial duplex nucleation, therefore it is important to note that these values underestimate thresholds for stable hybridization of free duplexes and are most applicable for forming segments that are covalently linked to another bound segment.

The lack of partial dehybridization in CCend-AP2 suggests its terminal base pair is stabilized relative to the other sequences, and we used ^1^H NMR spectroscopy to further examine the dominant duplex configurations of CCend and CGCcap sequences. Melting profiles for individual G:C base pairs were extracted from the temperature-dependent chemical shift of cytosine H5 nuclei isolated using ^1^H-^1^H TOCSY temperature series (Figs. 6b and S38 – S45). Interior cytosines (C2, C3) of CCend-WT and CGCcap-WT show sigmoidal transitions over the temperature range of the duplex melting transition measured with IR spectroscopy, but most of the frequency change in the terminal C1 H5 nuclei occurs at temperatures well below the melting transition. We assign the low-temperature behavior at C1 to fraying of the terminal base pair, which is further supported by the lack of such changes in GCGcore-WT where all cytosines are at the center of the duplex (Fig. S45). The temperature-dependence of the C1 H5 frequency in CCend-AP2 follows the duplex melting transition, suggesting that the terminal G:C base pair is intact and dissociates together with the rest of the duplex as indicated by 2D IR temperature series. The ^1^H-^1^H NOESY spectrum reveals intense cross peaks between C1 and T3 nuclei, indicating that C1 is stacked with T3 in the duplex state and presumably base paired with G2 rather than G1 (Figs. 6c, S40 – S41). This out-of-register base pairing creates a duplex with one stretch of base pairs, circumventing the nucleation penalty for forming a new segment. Similar out-of-register configurations have been proposed previously around AP sites.(28,69) In contrast, the temperature-dependent cytosine H5 frequencies in CGCcap-AP2 indicate partial dehybridization of the G:C cap more than 10 °C below the duplex melting transition. The TOCSY spectra of CGCcap-AP2 also show additional cytosine peaks from 2 to 26 °C that suggest multiple slowly interconverting duplex configurations may be present at low temperature (Figs. S43 – S44), yet the assignment of these structures are still unclear.

All-atom MD simulations provide insight into the possible base-pairing and stacking configurations adopted by CCend-AP2 and CGCcap-AP2. Free-energy surfaces (FESs) constructed from 20 × 1 µs simulations indicate that each sequence adopts a variety of non-canonical configurations that interconvert on sub-microsecond timescales (Figs. S46 – S47). Interpretation of the simulations results must be done with care since the AMBER-bsc1 force is not parameterized for AP sites. While we cannot anticipate accurate predictions of the relative stabilities of the various states, the predictions are nonetheless useful for identifying putative configurational states and their relative propensities to help interpret experimental measurements. Only 1% of CGCcap-AP2 configurations exhibit in-register base pairing between C1 and G1, and C1 instead prefers to stack with C3 and interact with C2. Similar for CCend-AP2, configurations with in-register base pairing between C1 and G1 only make up 0.4% of the population (Fig. 6d). The most populated states are those with C1 paired to G2 and the free deoxyribose group fully extrahelical (states B and C in Fig. 6d). State B (15.2%) is characterized by a Watson-Crick-Franklin (WCF) base pair between C1 and G2 with stacking of G1 over the base pair as often observed for duplexes with dangling ends.(70) The C1 nucleotide rotates approximately 180° in state C (24.7%) and forms a base pair composed of two amino-imino hydrogen bonds, N3(C) to N2H(G) and N4H(C) to N3(G), and G1 stacks only with G2. This type of base pair has previously been observed for mononucleotides binding to RNA templates and in other folded RNA structures.(71,72) Another common configuration involves C1 stacked between bases on the other strand (state D). State B is most consistent with the NOESY spectrum for CCend-AP2 because it is the only configuration where the H5 nuclei of C1 is close enough to the T3 methyl group (2.7 Å vs. 8.8 Å in state C and 8-10 Å in state D) to enable significant dipolar coupling to produce an intense cross-peak. While the NOESY data suggests a configuration like state B has the largest population experimentally, we cannot rule out the presence of states C and D as they interconvert with state B too quickly to be resolved as separate sets of NMR peaks.

### Potential role of out-of-register duplex configurations in repetitive sequences

The stability of the duplex for the repetitive T11 sequence can be affected by out-of-register base pairing, and 3SPN.2 MD simulations confirm the role of these configurations. Figure 7 shows duplex free-energy minima at *n*_*bp*_ = 10 for T11-WT, *n*_*bp*_ = 9 for T11-AP2, *n*_*bp*_ = 7 for T11-AP4, and *n*_*bp*_ = 5 for T11-AP6 corresponding to duplex configurations shifted by one, two, four, and six base-pair indices, respectively. The shifted configurations for T11-AP2 and T11-AP4 are the dominant duplex species even at *T*_*m,MD*_ − 20 °C, but the population of those for T11-WT and T11-AP6 grow with increasing temperature until the duplex fully dissociates (Figs. 7c and S12). Relative to unbound segments of in-register segment-dehybridized configurations found in CGCcap, CCends, and GCGcore sequences, the single-strand overhangs of out-of-register configurations have greater conformational entropy and a lower electrostatic penalty that lead to enhanced stability of the duplex region.(73)

**Figure 7.**
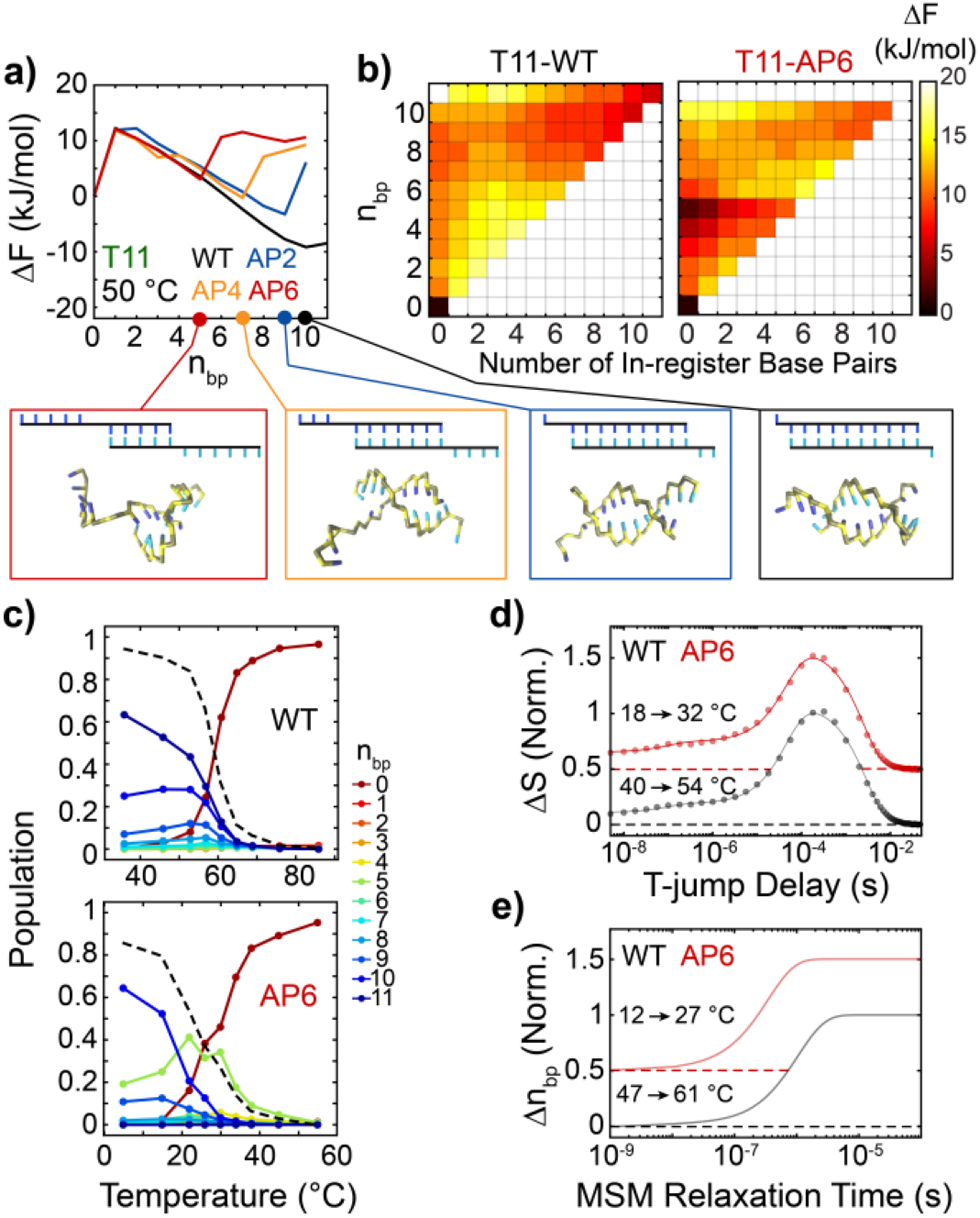
AP-site induced out-of-register base pairing in T11 from 3SPN.2 MD simulations. **(a)** FEPs as a function of *n*_*bp*_ for T11 sequences at 50 °C from 3SPN.2 MD simulations with WTMetaD. Representative structures show out-of-register shifted configurations at duplex free-energy minima. **(b)** 2D free-energy surfaces for (left) T11-WT and (right) T11-AP6 of the number of in-register base pairs vs. *n*_*bp*_. The free-energy minimum at *n*_*bp*_ = 5 for T11-AP6 corresponds to a completely out-of-register configuration. **(c)** Temperature-dependent population profiles of duplexes with *n*_*bp*_ ranging from 0 to 11 for (top) T11-WT and (bottom) T11-AP6. The black dashed line corresponds to the total fraction of intact base pairs. **(d)** Normalized Δ*S*(*t*) time traces at 1605 cm^-1^ (A:T) plotted for the T11-WT and T11-AP6. T-jumps are performed from approximately *T*_*m*_–15°C to *T*_*m*_ for each sequence. Traces are shifted vertically with respect to one another, and dashed lines indicate respective baselines. Solid lines correspond to three-component fits from global lifetime analysis. **(e)** MSM T-jump simulations of the normalized change in A:T base pairs, Δ*n*_*bp*_(*t*), using the same method as in Fig. 5d. Due to overlapping timescales for partial dehybridization to the out-of-register *n*_*bp*_ = 5 state and full-strand dissociation, only a single dehybridization kinetic component is observed in the relaxation profiles.

Out-of-register base pairing is commonly observed in coarse-grained MD simulations and statistical modelling of repetitive DNA oligonucleotides,(35,68,74-76) yet it is challenging to verify the population of such configurations experimentally. The IR spectral features of in- and out-of-register A:T base pairing are expected to be essentially identical, therefore our experimental data does not directly report on the presence of out-of-register base pairing. Additionally, the experimental temperature-dependent melting behavior of T11 sequences does not indicate a significant population of out-of-register configurations and suggest that 3SPN.2 simulations overestimate the thermodynamic stability of out-of-register configurations for T11 AP sequences. The probability of out-of-register hybridization pathways may be overestimated due to smoothening of the free-energy landscape generally induced by coarse-graining.(77) The total change in adenine FTIR and 2D IR signal upon thermal dissociation is nearly constant in T11-WT and each AP sequence, suggesting each duplex contains a similar degree of A:T base pairing at lowest measured temperature in the duplex state (Fig. S34). Further, 3SPN.2 predicts that partial dehybridization of T11-AP4 and T11-AP6 occurs at temperatures below *T*_*m,MD*_, which broadens the overall melting curve (reduces slope at *T*_*m,MD*_ by 20% relative to T11-WT). However, only a 6% reduction in the melting curve slope is observed experimentally.

We also find that MSM and T-jump IR relaxation are insensitive to the kinetics of out-of-register shifting (Figs. 7de and S17). Although a thermally-induced increase of the *n*_*bp*_ = 5 state is predicted for T11-AP6 and 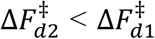, the MSM relaxation timescale for shifting overlaps with full-strand dissociation (Fig. 7e), which was observed previously in similar canonical oligonucleotides.(68) Shifting from the *n*_*bp*_ = 10 to *n*_*bp*_ = 5 state first entails breaking of all in-register contacts followed by formation of five out-of-register base pairs. The barrier height for shifting is likely much larger than indicated by 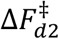 in Fig. 7a, and *n*_*bp*_ is a poor coordinate to describe the shifting transition. As a result, no partial-dehybridization response is observed in MSM relaxation profiles. Similarly, T-jump IR measurements only show responses for terminal fraying and full-strand dissociation and cannot indicate nor rule out a role of out-of-register base pairing in these sequences (Fig. 7d).

## Conclusions

Our study reveals molecular details underpinning the position-dependent duplex destabilization from an AP site within DNA oligonucleotides by combining temperature-dependent IR and NMR spectroscopy, T-jump IR kinetics, and coarse-grained MD simulations. An AP site destabilizes the duplex through a loss of stacking and base-pairing interactions that disrupt cooperative base pairing and introduce a free-energy barrier for nucleating base-pair segments on each side of the AP site. This nucleation barrier leads to a position-dependent duplex destabilization from the AP site. When the AP site is near the termini, the short segment is highly frayed due to the nucleation barrier. As the AP site moves inward by three to six base-pair sites, the short segment has enough stability to bind, and the frayed configuration becomes metastable. At this position, the nucleation barrier is fully encompassed into the duplex and the AP site is maximally destabilizing. The positions over which this transition occurs depends on the nucleobase sequence in the short segment and adjacent to the AP site, which leads to poor prediction of duplex destabilization by the NN model even at a fixed AP-site position. While the transition occurs at short-segment lengths of two to five base pairs depending on the G:C content of the segment, certain short-segment sequences may be able to circumvent the nucleation barrier by forming base pairs out-of-register as observed for CCends-AP2.

This work has focused on AP sites, yet there are numerous non-canonical base pairs, nucleobase modifications, and damaged-induced lesions that impart a comparable energetic penalty to duplex formation 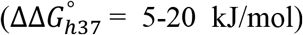.(8,13,78,79) Although they alter local base-pairing and stacking interactions in different ways, such modifications and non-canonical base pairs may disrupt base-pairing cooperativity and follow a position-dependence similar to an AP site. For example, base-pair mismatches impose a sequence-dependent free-energy penalty to duplex formation that follows a similar position-dependence as for AP sites.(8,19,20) Greater reduction in duplex stability occurs as the mismatch moves inward 3 – 4 base pairs from the termini. While the base-pairing geometry of mismatches is highly sequence-dependent, the ‘defect penalty’ used to model their behavior is essentially the same as the nucleation penalty described in our work and supports that the position-dependent thermodynamic and dynamic consequences from destabilizing modifications may be similar to what we find for AP sites.

## Data Availability

Python scripts for generating abasic configurations from intact 3SPN.2 files, performing metadynamics simulations, and reweighting free-energy surfaces are available at https://github.com/mrjoness/abasic-thermo/. Scripts for running equilibrium simulations and building Markov state models are available at https://github.com/mrjoness/abasic-kinetics/. All PLUMED input scripts were submitted to the PLUMED-NEST public repository and are available at https://www.plumed-nest.org/eggs/22/037/.

Initialization files and trajectory data for unbiased coarse-grained simulations used to construct Markov state models, biased coarse-grained simulations over multiple temperatures, and all-atoms simulations of AP2 sequences are uploaded to Zenodo and available at 10.5281/zenodo.8169462. Free-energy profiles along *n*_*bp*_ and *r*_*bp*_, Markov state model relaxation profiles, time- and rate-domain t-HDVE data, FTIR temperature series, 2D IR temperature series, and NMR temperature series are available at 10.5281/zenodo.8174843.

## Supplementary Data

Processing of FTIR melting curves; isothermal titration calorimetry of hybridization; description of two-stretch helix coil model; Analysis of metastable states from 3SPN.2 MD simulations; temperature-dependent FEPs from 3SPN.2 MD simulations; FTIR, 2D IR, and NMR TOCSY and NOESY spectra; analysis of temperature-dependent T-jump IR data; Validation of Markov state models generated from 3SPN.2 MD simulations

## Acknowledgements

This work was supported by the National Institute of General Medical Sciences of the National Institutes of Health (Award No. R01-GM118774) and the National Science Foundation under Grant No. CHE-2152521 and CHE-2155027. B.A. acknowledges support from the NSF Graduate Research Fellowship Program. This work was completed in part with resources provided by the University of Chicago Research Computing Center. We gratefully acknowledge computing time on the University of Chicago high-performance GPU-based cyberinfrastructure supported by the National Science Foundation under Grant No. DMR-1828629.

