## Supplemental Text and Figures for "Molecular insight into how the position of an abasic site and its sequence environment influence DNA duplex stability and dynamics"

#### Table of Contents

**S1. Thermodynamics of duplex hybridization**……………………………………………..…S2
S1.1 Temperature-dependent FTIR spectra
S1.2 Two-state thermodynamic model of duplex hybridization
S1.3 Hybridization enthalpies from isothermal titration calorimetry

**S2. Two-stretch helix-coil model**………………...............................................................S9
S2.1 DNA duplex melting without an abasic site
S2.2 DNA duplex melting with an abasic site
S2.3 Base-pairing probability distributions and free-energy profiles

**S3. Free-energy profiles and thermodynamics from 3SPN.2 simulations**……….……S16
S3.1 Comparison of duplex hybridization thermodynamics from 3SPN.2 and FTIR
S3.2 Base-pairing contributions to local FEP minima
S3.3 Temperature-dependent FEPs and duplex populations

**S4. Temperature-jump IR spectroscopy**……………………………………..……..….S26
S4.1 Temperature-dependent t-HDVE data
S4.2 Global lifetime analysis of t-HDVE data
S4.3 Temperature-dependent trends in $\tau_1$ response
S4.4 Dehybridization and hybridization kinetics from $\tau_3$ response
S4.5 Kinetics of segment-dehybridization

**S5. Construction and validation of Markov state models from 3SPN.2 MD simulations...S37**

**S6. Assessing deviation from two-state melting behavior…………………………………..S40**
S6.1 Evaluating partial dehybridization from total FTIR and 2D IR spectral change
S6.2 Partial dehybridization from FTIR and 2D IR temperature series
S6.3 $^1$H NMR spectroscopy of partial dehybridization in AP2 sequences
S6.4 All-atom MD simulations of AP2 sequences

### S1. Thermodynamics of duplex hybridization

#### S1.1 Temperature-dependent FTIR spectra

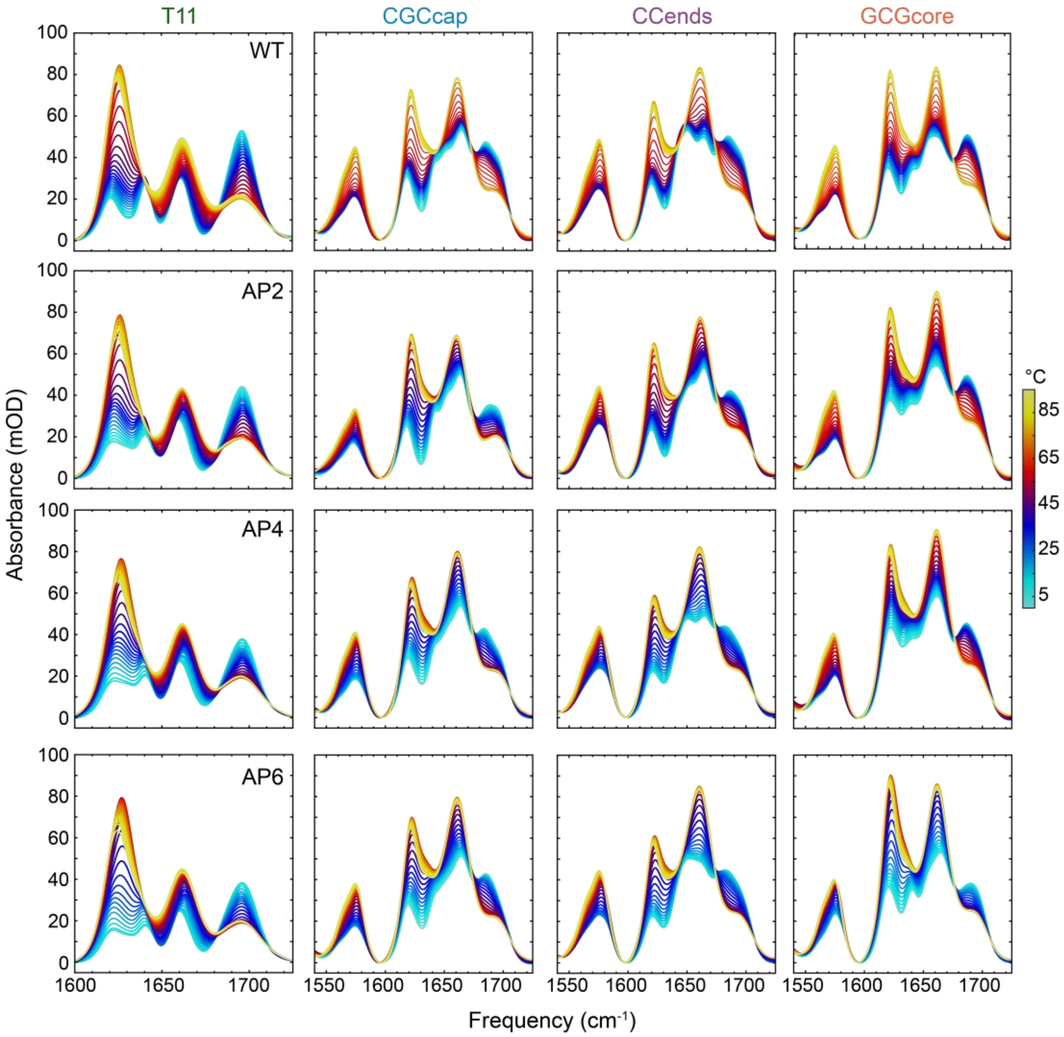


**Figure S1. FTIR temperature series.** FTIR temperature series for T11 (1st column), CGCcap (2nd column), CCends (3rd column), and GCGcore (4th column) sequences measured from 3.8 to 93.3 °C in ~2.6 °C steps. All samples were prepared in deuterated pH* 6.8 400 mM SPB with a total oligonucleotide concentration of 2 mM.

##### S1.2 Two-state thermodynamic model of duplex hybridization

The FTIR temperature series of each DNA sequence studied here exhibit a single sigmoidal melting transition and therefore can be most simply described using a two-state model of the duplex (*D*)-to-single-strand (*S*) transition for non-self-complementary oligonucleotides. We use a standard treatment for optical melting curves that was previously described.(1) In brief, the fraction of intact duplexes ($\theta_D$) is described using a temperature-dependent dissociation constant ($K_d$)

$$\theta_D = 1 + \frac{K_d - \sqrt{K_d^2 + 2c_{tot}K_d}}{c_{tot}} \tag{S1}$$

$c_{tot}$ is the total oligonucleotide concentration. Eq. S1 assumes a 1:1 molar ratio between complementary DNA strands ([$S_1$] = [$S_2$]), and all FTIR measurements in this work were performed under this condition. The temperature dependence of $\theta_D$ and $K_d$ are determined by the hybridization enthalpy ($\Delta H_h^\circ$) and entropy ($\Delta S_h^\circ$) difference between the duplex and single-strand states.

$$K_d(T) = \exp\left[-\frac{-\Delta H_h^\circ(T)}{RT} + \frac{-\Delta S_h^\circ(T)}{R}\right] \tag{S2}$$

In practice, $\Delta H_h^\circ$ and the melting temperature ($T_m$), which here is defined as the temperature where $\theta_D$ = 0.5, are used as free parameters to fit the FTIR melting data.

$$T_m = \frac{\Delta H_h^\circ(T_m)}{\Delta S_h^\circ(T_m) - R\ln(c_{tot}/4)} \tag{S3}$$

The 2$^{nd}$ SVD components from FTIR temperature series were then fit to $\theta_D$ with baselines described by additional slope ($m_D$, $m_S$) and intercept ($b_D$ $b_S$) terms as shown in Fig. S2a.

$$V_{Fit}^{(2)} = \theta_D(L_D - L_S) + L_S \tag{S4}$$

$$L_D = m_D T + b_D \tag{S5a}$$

$$L_S = m_S T + b_S \tag{S5b}$$

Thermodynamic values extracted from the fits are shown in Table S1 and Fig.S3.

**Table S1**. Thermodynamics parameters for duplex melting curves determined from two-state fits to FTIR temperature series 2nd SVD components (Fig. S2a).

| Sequence | | $\Delta H_h^\circ$ (kJ/mol) | $\Delta S_h^\circ$ (J/mol K) | $\Delta G_{h37}^\circ$ (kJ/mol) | $T_m$ (°C) | $\Delta\Delta H_h^\circ$ (kJ/mol) | $\Delta\Delta S_h^\circ$ (J/molK) | $\Delta\Delta G_{h37}^\circ$ (kJ/mol) | $\Delta T_m$ (°C) |
|---|---|---|---|---|---|---|---|---|---|
| T11 | WT | -271 ± 8 | -769 ± 26 | -32.8 ± 4.2 | 52.8 ± 0.2 | - | - | - | - |
| | AP2 | -274 ± 14 | -790 ± 44 | -29.5 ± 7.2 | 48.5 ± 0.3 | 3 ± 32 | 21 ± 51 | 3.3 ± 8.3 | -4.3 ± 0.4 |
| | AP4 | -236 ± 16 | -686 ± 50 | -23.5 ± 8.2 | 42.1 ± 0.5 | 35 ± 36 | 83 ± 56 | 9.3 ± 9.2 | -10.7 ± 0.5 |
| | AP6 | -224 ± 12 | -665 ± 40 | -17.4 ± 6.2 | 34.0 ± 0.5 | 48 ± 30 | 104 ± 48 | 15.4 ± 7.5 | -18.9 ± 0.5 |
| CGCcap | WT | -310 ± 6 | -869 ± 16 | -40.9 ± 2.8 | 59.8 ± 0.1 | - | - | - | - |
| | AP2 | -212 ± 6 | -605 ± 19 | -24.5 ± 3.0 | 44.3 ± 0.2 | 98 ± 8 | 264 ± 25 | 16.3 ± 4.1 | -15.5 ± 0.3 |
| | AP4 | -205 ± 7 | -593 ± 24 | -21.1 ± 3.7 | 39.1 ± 0.3 | 105 ± 9 | 276 ± 29 | 20.0 ± 4.7 | -20.6 ± 0.4 |
| | AP6 | -180 ± 10 | -512 ± 30 | -21.4 ± 4.7 | 40.0 ± 0.6 | 130 ± 11 | 357 ± 34 | 19.5 ± 5.5 | -19.7 ± 0.6 |
| CCends | WT | -342 ± 10 | -970 ± 32 | -41.5 ± 5.3 | 58.1 ± 0.2 | - | - | - | - |
| | AP2 | -269 ± 11 | -770 ± 34 | -30.2 ± 5.6 | 49.5 ± 0.2 | 73 ± 15 | 200 ± 46 | 11.3 ± 7.7 | -8.5 ± 0.3 |
| | AP4 | -164 ± 11 | -469 ± 35 | -18.7 ± 4.8 | 35.2 ± 0.9 | 178 ± 15 | 501 ± 48 | 22.8 ± 7.6 | -22.9 ± 0.9 |
| | AP6 | -183 ± 9 | -535 ± 31 | -17.0 ± 4.8 | 32.6 ± 0.6 | 159 ± 14 | 435 ± 44 | 24.5 ± 7.1 | -25.5 ± 0.6 |
| GCGcore | WT | -284 ± 6 | -779 ± 20 | -42.8 ± 3.3 | 64.4 ± 0.2 | - | - | - | - |
| | AP2 | -279 ± 11 | -767 ± 32 | -41.4 ± 5.7 | 63.1 ± 0.3 | 5 ± 13 | 12 ± 40 | 1.4 ± 6.6 | -1.3 ± 0.4 |
| | AP4 | -210 ± 7 | -576 ± 21 | -31.7 ± 3.5 | 55.8 ± 0.3 | 74 ± 9 | 203 ± 29 | 11.1 ± 4.8 | -8.6 ± 0.4 |
| | AP6 | -140 ± 13 | -396 ± 42 | -17.1 ± 6.5 | 31.5 ± 1.7 | 145 ± 14 | 384 ± 46 | 25.7 ± 7.1 | -32.9 ± 1.8 |

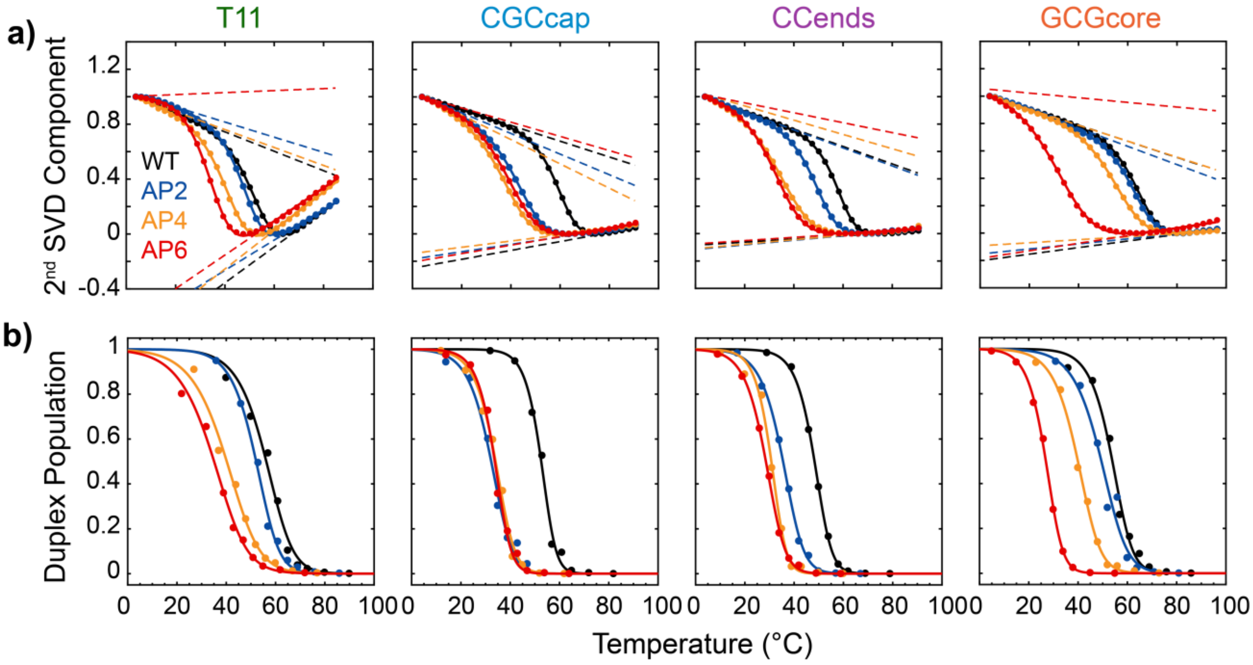


**Figure S2. Fitting of melting curves from FTIR temperature series and 3SPN.2 MD simulations to two-state model. (a)** Second component from singular value decomposition (SVD) of FTIR temperature series fit to two-state thermodynamic model (solid lines, eq. S4). Dashed lines indicate low ($L_D$) and high ($L_S$) baselines from the fits. **(b)** Probability of oligonucleotides being in the duplex ensemble as a function of temperature extracted from 3SPN.2 MD simulations with WTMetaD using the approach described previously.(1) Data are fit to eq. S1 (solid lines).

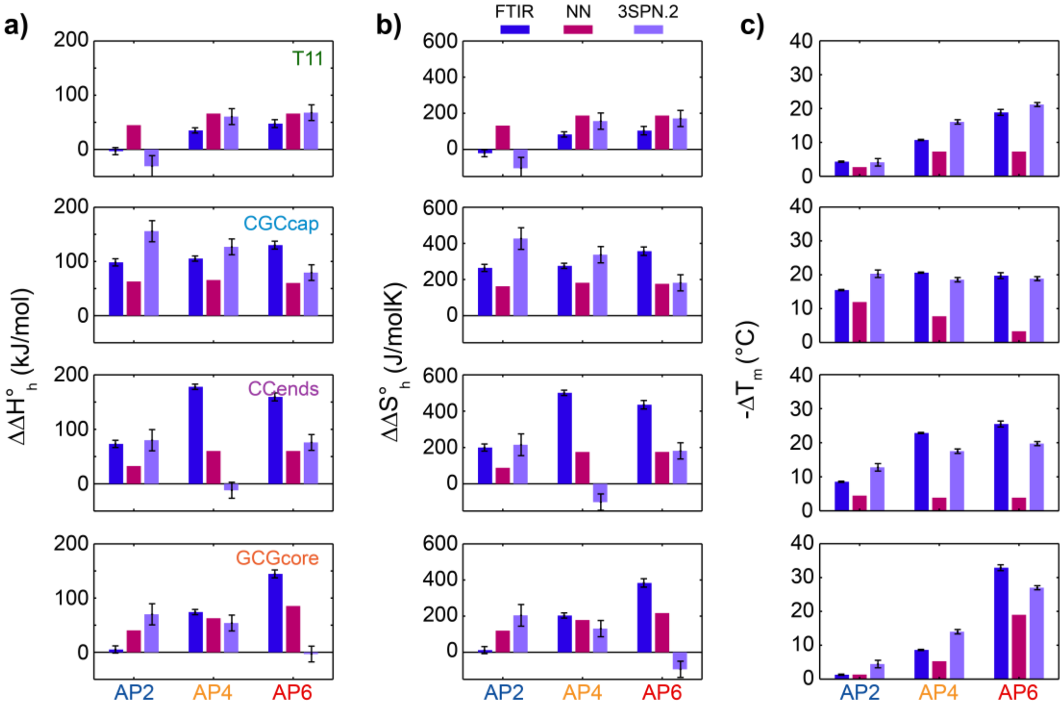

**Figure S3. Duplex dehybridization thermodynamics from two-state analysis.** Drop in duplex dehybridization enthalpy (**a**, $\Delta\Delta H_h^{\circ}$), entropy (**b**, $\Delta\Delta S_h^{\circ}$), and melting temperature (**c**, $\Delta T_m$) for each AP sequence with respect to the WT sequence from fits to the FTIR melting curve (eq. S4) and compared with values from Santa Lucia's nearest-neighbor (NN) model(2,3) and those generated from 3SPN.2-determined melting curves. FTIR and 3SPN.2 error bars correspond to 95% confidence intervals from two-state fits.

##### S1.3 Hybridization enthalpies from isothermal titration calorimetry

Isothermal titration calorimetry (ITC) measurements were performed using a MicroCal iTC200 (Malvern Panalytical). Single-strand oligonucleotides were prepared in the cell with a concentration of 10-30 μM and the complement strand was placed in the syringe with a concentration of 100-300 μM. All samples were prepared in non-deuterated pH 6.8 400 mM SPB and degassed under vacuum for >20 min at the experimental temperature prior to each measurement. Sample conditions for each measurement are listed in Table S1.

ITC measurements started with an initial 0.4 μL injection followed by 19 2 μL aliquot injections of the titrant oligonucleotide solution into the cell. Injections of 4 s duration were performed every 180 s, and the syringe needle constantly stirred the cell solution with a spin rate of 1000 rpm. The power required to re-equilibrate the sample and reference cells was monitored as a function of time. The power profile of each injection was integrated over time to obtain the change in heat from the injection (ΔQ). The heat of dilution was measured for the titrant at each experimental temperature and subtracted from ΔQ after integration. Curves of ΔQ vs molar ratio of titrant and cell oligonucleotide were fit to a single site binding model(4) with free parameters for binding enthalpy ($\Delta H_h^{\circ}$), binding constant ($K_a$), and reaction stoichiometry ($n$).

$$\Delta Q_i = Q_i - Q_{i-1} + \frac{V_{inj}}{V_o}\left[\frac{Q_i + Q_{i-1}}{2}\right] \tag{S6}$$

$$Q_i = \frac{nM_{c,i}V_o\Delta H^{\circ}}{2}\left[1+\frac{T_{c,i}}{nM_{c,i}}-\frac{1}{nK_aM_{c,i}}-\sqrt{\left(1+\frac{T_{c,i}}{nM_{c,i}}+\frac{1}{nK_aM_{c,i}}\right)^2-\frac{4T_{c,i}}{nM_{c,i}}}\right] \tag{S7}$$

$Q_i$ is the total heat exchanged after injection $i$ of titrant. $M_{c,i}$ and $T_{c,i}$ are the concentration of cell oligonucleotide and titrant oligonucleotide, respectively, in the cell after injection $i$. $V_o$ is the initial volume of sample in the cell, which was 200 μL for all measurements, and $V_{inj}$ is the volume of titrant used in each injection (2 μL). ITC measurements were performed at multiple temperatures for WT sequences to determine the change in heat capacity from the single-strand to duplex state ($\Delta C_p$, Fig. S4).

ITC thermograms for each sequence are shown in Fig. S5. $\Delta H_h^\circ$ extracted from fits of the thermograms are compared with values extracted from FTIR temperature series. The FTIR $\Delta H_h^\circ$ values are corrected to the ITC measurement temperature using $\Delta C_p$.

$$\Delta H_h^\circ(T) = \Delta H_h^\circ(T_m) + \Delta C_p(T - T_m) \tag{S8}$$

After correction, good agreement is found between ITC and FTIR $\Delta H_h^\circ$ values (Fig. 5b).

**Table S2.** Experimental conditions used for ITC measurements shown in Fig. S5. All samples were prepared in non-deuterated 400 mM pH 6.8 sodium phosphate buffer (SPB).

| Sequence | T (°C) | Syringe Strand | [Syringe] (μM) | Cell Strand | [Cell] (μM) |
|---|---|---|---|---|---|
| | | | **T11** | | |
| WT | 25 | AAAAAAAAAAA | 180 | TTTTTTTTTTT | 15 |
| AP2 | 20 | AAAAAAAAAAA | 180 | T_TTTTTTTTT | 15 |
| AP4 | 15 | AAAAAAAAAAA | 160 | TTT_TTTTTTT | 15 |
| AP6 | 10 | AAAAAAAAAAA | 180 | TTTTT_TTTTT | 15 |
| | | | **CGCcap** | | |
| WT | 25 | ATATATATGCG | 120 | CGCATATATAT | 10 |
| AP2 | 15 | ATATATATGCG | 140 | C_CATATATAT | 15 |
| AP4 | 10 | ATATATATGCG | 220 | CGC_TATATAT | 20 |
| AP6 | 10 | ATATATATGCG | 245 | CGCAT_TATAT | 20 |
| | | | **CCends** | | |
| WT | 25 | GGATATATAGG | 100 | CCTATATATCC | 10 |
| AP2 | 17 | GGATATATAGG | 100 | C_TATATATCC | 10 |
| AP4 | 10 | GGATATATAGG | 220 | CCT_TATATCC | 20 |
| AP6 | 10 | GGATATATAGG | 220 | CCTAT_TATCC | 20 |
| | | | **GCGcore** | | |
| WT | 25 | ATATCGCTATA | 110 | TATAGCGATAT | 10 |
| AP2 | 25 | ATATCGCTATA | 110 | T_TAGCGATAT | 10 |
| AP4 | 18 | ATATCGCTATA | 160 | TAT_GCGATAT | 15 |
| AP6 | 10 | ATATCGCTATA | 2000 | TATAG_GATAT | 200 |

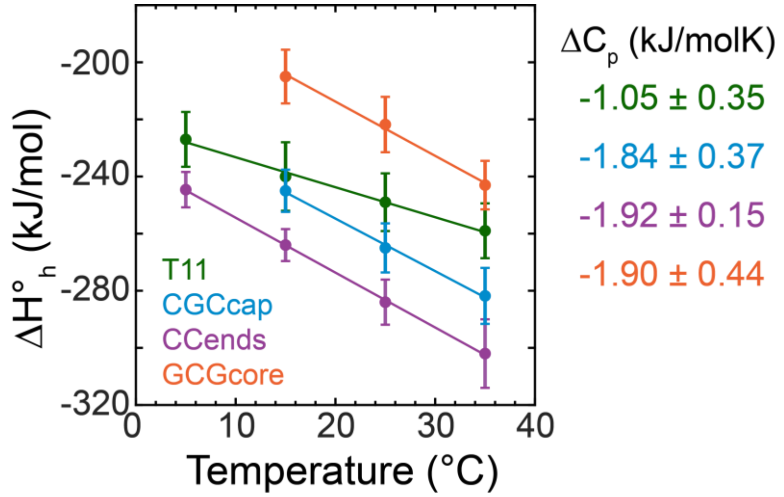


**Figure S4. Determination of $\Delta C_p$ from ITC.** $\Delta H_h^{\circ}$ determined with ITC as a function of temperature for T11-WT, CGCcap-WT, CCends-WT, and GCGcore-WT sequences. Vertical error bars indicate 95 % confidence intervals from nonlinear least squares fitting of ITC thermograms. Solid lines correspond to linear fits $\Delta H_h^{\circ}$ data, and the slope ($\Delta C_p$) is listed for each sequence.

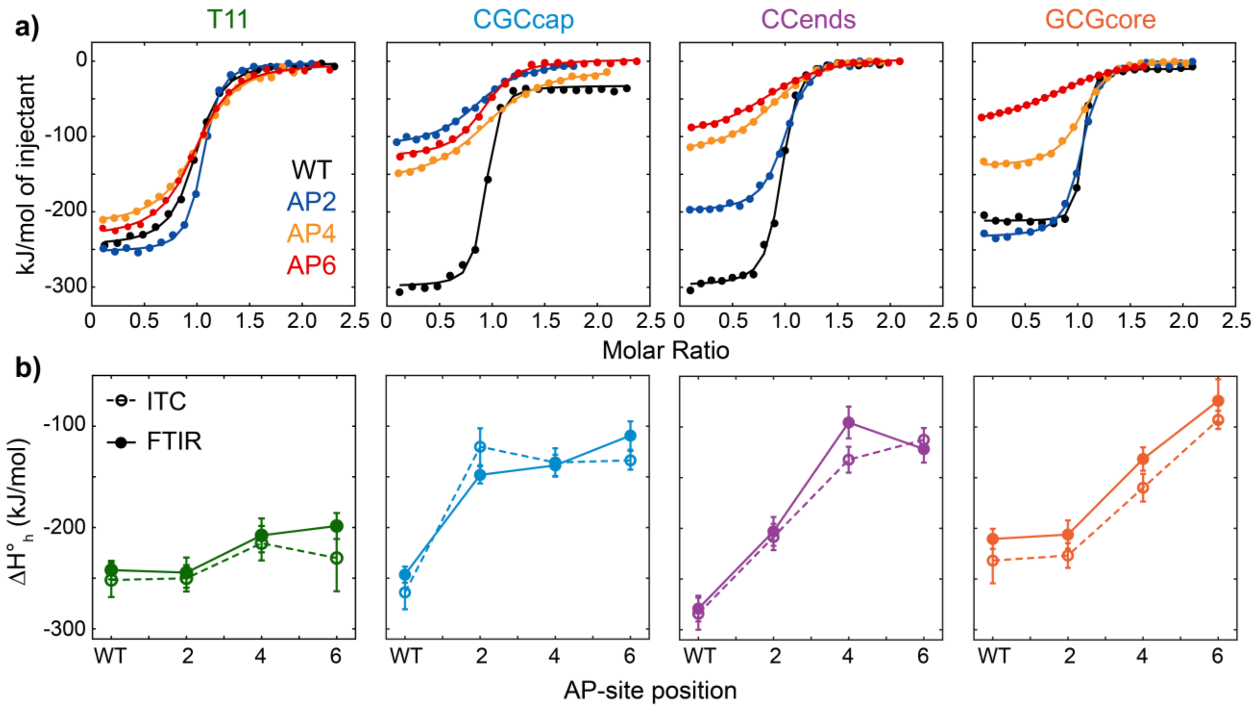


**Figure S5. DNA duplex hybridization monitored with ITC (a)** ITC thermograms expressed as molar heat of injectant vs. molar ratio between cell and titrant oligonucleotides. Measurements were performed in non-deuterated 400 mM pH 6.8 SPB using variable sample concentration and cell temperatures ranging from 10 to 25 °C depending on the respective binding constant (Table S1). Data are fit to a two-state model (solid lines, eq. S6). **(b)** $\Delta H_h^{\circ}$ determined from ITC (open circle with dashed line) compared with $\Delta C_p$-corrected values from FTIR temperature series (filled circle with solid line). Error bars for ITC indicate average 95% confidence intervals from fits of ITC thermograms and those of FTIR are propagated from ITC and FTIR fit errors.

#### S2. Two-stretch helix-coil model

To gain insight into the sequence-dependent melting behavior of DNA duplexes containing an AP site, we need to go beyond to the two-state thermodynamic description of DNA hybridization (Section S1.2) and assess the characteristics of the duplex state. Along this line, we employ a cooperative helix-coil transition model based on the perfect-matching helix-coil models of Poland and Scheraga and others.(5,6) Helix-coil models describe the statistics of base pairing in the DNA duplex and have been extended for improved prediction of DNA melting behavior.(7) Base-pairing thermodynamics are primarily described by a nucleation parameter ($\sigma$) that contains the statistical weight for forming the first base pair in an unbound region as well as an equilibrium constant ($s$) for each base pairing formation step adjacent to an already formed base pair.

$$s_i = e^{-\Delta G^{\circ}_{\mathrm{int},i}/RT} \tag{S9}$$

$$\sigma_i = e^{-\Delta G^{\circ}_{\mathrm{nuc},i}/RT} \tag{S10}$$

$\Delta G^{\circ}_{int}$ and $\Delta G^{\circ}_{nuc}$ are the free energies changes for base-pair formation and nucleating a stretch of base pairs. To fully account for DNA sequence effects, $s$ and σ will be different for each base-pair site $i$ within the duplex.

##### S2.1 DNA duplex melting without an abasic site

The most general form of the base-pairing partition function, $Z_{int,D}$, describes the statistics of base pairing configurations for a duplex with $N$ possible base pairs and accounts for all possible dissociation bubble configurations:

$$Z_{\mathrm{int},D}(N) = \sum_{n_{bp}=0}^{N} \sum_{\nu=0}^{\nu_{\max}} \left[ \prod_{j=1}^{n_{bp}} s_{i,j} \prod_{\ell=1}^{\nu} \sigma_{i,\ell} \right] \tag{S11}$$

For a given number of intact base pairs, $n_{bp}$, there are base-pairing configurations with variable number of continuous base-pairing stretches, $\nu$, that depend on $n_{bp}$ and $N$ and the maximum number of possible base-pair stretches ($\nu_{max}$). The energy of a given microstate is determined by products of $s$ across each intact base pair site, $j$, and σ across each stretch nucleation site, $\ell$. While eq. S11 accounts for all possible base-pair configurations, it is too general to be of practical use and can be significantly simplified by making a couple of key assumptions. First, we assume that the thermodynamics of the duplex-to-single-strand transition can largely be captured by a single $s$ parameter, $\langle s \rangle$, that is an average over all in-register base-pairing interactions, and that σ is independent of base-pairing position. In practice, $\langle s \rangle$ is computed from the average nearest-neighbor (NN) free-energy parameter across the sequence.

$$\langle s \rangle = e^{-\Delta G^{\circ}_{\text{int}}/RT} = \frac{\sum_{i}^{N-1} \Delta H^{\circ}_{NN,i} - T \sum_{i}^{N-1} \Delta S^{\circ}_{NN,i}}{N-1} \tag{S12}$$

In eq. S12, $\Delta H^{\circ}_{NN}$ and $\Delta S^{\circ}_{NN}$ come from Santa Lucia's NN parameters for duplex DNA with a correction for counterion concentration.(2,8) The use of a single average $s$ will partially account for sequence-dependent behavior of full dissociation, but it cannot properly capture pre-melting effects such as terminal fraying. The simplified form of $Z_{int,D}$ with $\langle s \rangle$ and site-independent σ is:

$$Z_{\text{int},D}(N) = \sum_{n_{bp}=0}^{N} \sum_{\nu=0}^{\nu_{\max}} \left[ W\left(N, n_{bp}, \nu\right) \langle s \rangle^{n_{bp}} \sigma^{\nu} \right] \tag{S13}$$

In eq. S13, a degeneracy term, $W$, corresponds to the number of microstates with given values of $N$, $n_{bp}$, and $\nu$. Equation S13 can be further simplified by neglecting microstates with multiple stretches of base pairs. The entropic penalty of forming short loops of dissociated base pairs makes configurations with multiple base pairing stretches highly unfavorable in short canonical sequences like those studied in this work ($N$ = 11). Previous statistical models and simulations have observed negligible population of configurations with loops in short canonical sequences,(9-11) and experiments have indicated that a minimum of ~20 A:T base pairs flanked by G:C regions are needed to generate non-negligible bubble populations.(12) Therefore, we limit $Z_{int,D}$ to only consider single stretch configurations ($\nu$ = 1) with in-register base pairing.(5,6,13) For a single base-pairing stretch, the degeneracy factor, $W$, is simplified to $N - n_{bp}$+1.

$$Z_{\text{int},D}(N) = 1 + \sigma \sum_{n_{bp}=1}^{N} \kappa^{n_{bp}} (N - n_{bp} + 1) \langle s \rangle^{n} \tag{S14}$$

$\kappa$ is an entropic correction factor that is used to shift the calculated $T_m$ for WT sequences to match experimental values. The choice of $\kappa$ will depend on σ. From $Z_{int,D}$, the average fraction of intact base pairs per duplex ($\theta_{int}$) can be determined.

$$\theta_{\text{int}} = \frac{\langle s \rangle}{N} \frac{\partial \ln Z_{\text{int},D}}{\partial \langle s \rangle} \tag{S15}$$

Short duplexes typically undergo full-strand dissociation at temperatures below the point at which $\theta_{int}$ ~ 0 due to non-negligible contributions from the external partition functions ($Z_{ext,i}$) of the single-strand and duplex species. The dissociation equilibrium constant ($K_d$) defined in eq. S2 can be re-cast in terms $Z_{ext,i}$ and $Z_{int,i}$ for the single-strands and duplex.

$$K_d = \frac{[S_1][S_2]}{[D]} = \frac{Z_{\text{int},S1}^{N_{S1}} Z_{ext,S1} Z_{\text{int},S2}^{N_{S2}} Z_{ext,S2}}{Z_{\text{int},D}^{N_D} Z_{ext,D}} \tag{S16}$$

$N_{S1}$ and $N_{S2}$ are the number of each single-strand molecules in solution. The external partition functions accounts for the greater translational entropy in the single-strand state compared to the duplex as well as the increased stability of the duplex as $c_{tot}$ increases. Similar to a lattice gas model,(5) we calculate the external partition functions using the number of possible ways to arrange single-strand and duplex DNA molecules on a 3D lattice. The volume of a single lattice site ($V_{ss}$) is set to that of a single-strand molecule where $V_{ss} = 4\pi R_G^3/3$. We use $R_G$ = 1.47 nm for $N$ = 11 and $[Na^+]$ = 600 mM based on the experimentally determined length and ionic strength scaling relationships for the radius of gyration ($R_G$) of single-strand DNA.(14) We neglect any minor sequence-dependence to $R_G$ and use a the same value of $V_{ss}$ for all oligonucleotides in this work. $Z_{ext,S1}$ and $Z_{ext,S2}$ are determined from the number of ways of placing $N_{S1/S2}$ single-strand molecules on a 3D lattice with $M$ sites, which in the dilute limit ($N_{S1/S2} << M$) is:

$$Z_{ext,S1/S2} \approx \frac{M^{N_{S1/S2}}}{N_{S1/S2}!} \quad \text{when} \quad N_{S1/S2} << M \tag{S17}$$

$M$ is defined by $M = V/V_{ss}$ where $V$ is the total system volume. Duplexes are defined when a single-strand occupies any of the 6 lattice sites adjacent to another single strand. $Z_{ext,D}$ then corresponds to the number of ways to arrange the number of duplexed oligonucleotides, $N_D$, on $M$ lattice sites where $N_D$ are restricted to the 6 adjacent sites around an already occupied site.

$$Z_{ext,D} \approx \frac{M^{N_D} 6^{N_D}}{N_D!} \quad \text{when } N_D << M \tag{S18}$$

We can put the external partition functions in terms of known parameters by first determining the chemical potential ($\mu$) of each species $i$.

$$\mu_i = -k_B T\left(\partial \ln Z_{tot,i} / \partial N_i\right) \text{ where } Z_{tot,i} = Z_{ext,i} Z_{\text{int},i}^{N_i} \tag{S19}$$

Combining eq. S19 with the equilibrium definition $\mu_{S1} + \mu_{S2} = \mu_D$, the following relation can be derived.

$$\frac{N_{S1} N_{S2}}{N_D} = \frac{M Z_{\text{int},S1} Z_{\text{int},S2}}{6 Z_{\text{int},D}} \tag{S20}$$

Using eq. S20 and relating the number of molecules in the system to concentration, $N_i = \mathrm{N}_A [i] V$, we can re-express $K_d$ (eq. S16).

$$K_d = \frac{1}{\gamma} \frac{Z_{\text{int},S1} Z_{\text{int},S2}}{Z_{\text{int},D}} \text{ where } \gamma = 6 c^{\circ} \mathrm{N}_A V_{ss} \tag{S21}$$

$\mathrm{N_A}$ is Avogadro's constant and $c^{\circ}$ is the standard state concentration of 1 M. Our treatment of the internal partition functions exclusively consider base pairing configurations, therefore $Z_{int,S1}$ and $Z_{int,S2}$ each have a statistical weight of 1 relative to $Z_{int,D}$ and eq. S21 can be further simplified.

$$K_d = \frac{1}{\gamma Z_{\text{int},D}} \tag{S22}$$

Just as for the two-state thermodynamic model described in Section S1.2, the fraction of oligonucleotide strands in the duplex state ($\theta_{ext}$) is determined by $K_d$ and $c_{tot}$.

$$\theta_{ext} = 1 + \frac{K_d - \sqrt{K_d^2 + 2c_{tot}K_d}}{c_{tot}} \tag{S23}$$

Eqs. S22 and S23 can be combined to give an expression for $\theta_{ext}$ in terms of $Z_{int,D}$.

$$\theta_{ext} = 1 + \frac{1 - \sqrt{1 + 2c_{tot}\gamma Z_{\text{int},D}}}{c_{tot}\gamma Z_{\text{int},D}} \tag{S24}$$

**S2.2 DNA duplex melting with an abasic site**

Our helix-coil model of DNA duplexes containing an AP site is illustrated in Fig. 2. We model the AP site as a defect in the duplex that splits base pairing into two possible stretches of length $N_1$ and $N_2$, where $N_1 + N_2 = N - 1$. For simplicity, each stretch is considered independent and contains its own $\langle s \rangle$ and σ parameters. The $\langle s \rangle$ parameter for each stretch is determined from average NN parameters across each stretch.(2)

$$\left\langle s_j \right\rangle = e^{-\Delta G^{\circ}_{\text{int},j}/RT} = \frac{\sum_{i}^{N_j-1} \Delta H^{\circ}_{NN,i} - T\sum_{i}^{N_j-1} \Delta S^{\circ}_{NN,i}}{N_j - 1} \quad j = 1,2 \text{ when } N_j > 1 \tag{S25a}$$

$$\left\langle s_j \right\rangle = e^{-\Delta G^{\circ}_{\text{int},j}/RT} = \Delta H^{\circ}_{NN} - T\Delta S^{\circ}_{NN} \quad j = 1,2 \text{ when } N_j = 1 \tag{S25b}$$

When a stretch has only a single base pair ($N_1$ or $N_2 = 1$), $\langle s \rangle$ is approximated by the NN parameter for a single dinucleotide step. By the design of $\langle s \rangle$, then, $Z_{int,D}$ can be described as a product of internal partition functions for stretches 1 and 2.

$$Z_{\text{int},D1} = 1 + \sigma_1 \sum_{n_{bp,1}=1}^{N_1} \kappa^{n_{bp,1}} (N_1 - n_{bp,1} + 1) \left\langle s_1 \right\rangle^{n_{bp,1}} \tag{S26a}$$

$$Z_{\text{int},D2} = 1 + \sigma_2 \sum_{n_{bp,2}=1}^{N_2} \kappa^{n_{bp,2}} (N_2 - n_{bp,2} + 1) \left\langle s_2 \right\rangle^{n_{bp,2}} \tag{S26b}$$

$$Z_{\text{int},D} = Z_{\text{int},D1} Z_{\text{int},D2} \tag{S26c}$$

The form of $\theta_{int}$ is split into a term for each stretch while $\theta_{ext}$ still follows eq. S23.

$$\theta_{\text{int}} = \frac{\langle s_1 \rangle}{N_1 + N_2} \frac{\partial \ln Z_{\text{int},D1}}{\partial \langle s_1 \rangle} + \frac{\langle s_2 \rangle}{N_1 + N_2} \frac{\partial \ln Z_{\text{int},D2}}{\partial \langle s_2 \rangle} \quad \text{(S27)}$$

A key effect of splitting the duplex into two stretches is that the total number of possible base-pairing arrangements ($W_{tot}$) is reduced. For a single base-pairing stretch, $W_{tot}$ is determined from the sum over $W$ in eq. S14.

$$W_{tot}(N) = \frac{N(N+1)}{2} \quad \text{(S28)}$$

For sequences containing an AP site, $W_{tot}$ is computed from the number of possible base-pairing configurations in each stretch.

$$W_{tot}(N_1, N_2) = \frac{N_1(N_1+1)}{2} + \frac{N_2(N_2+1)}{2} \quad \text{(S29)}$$

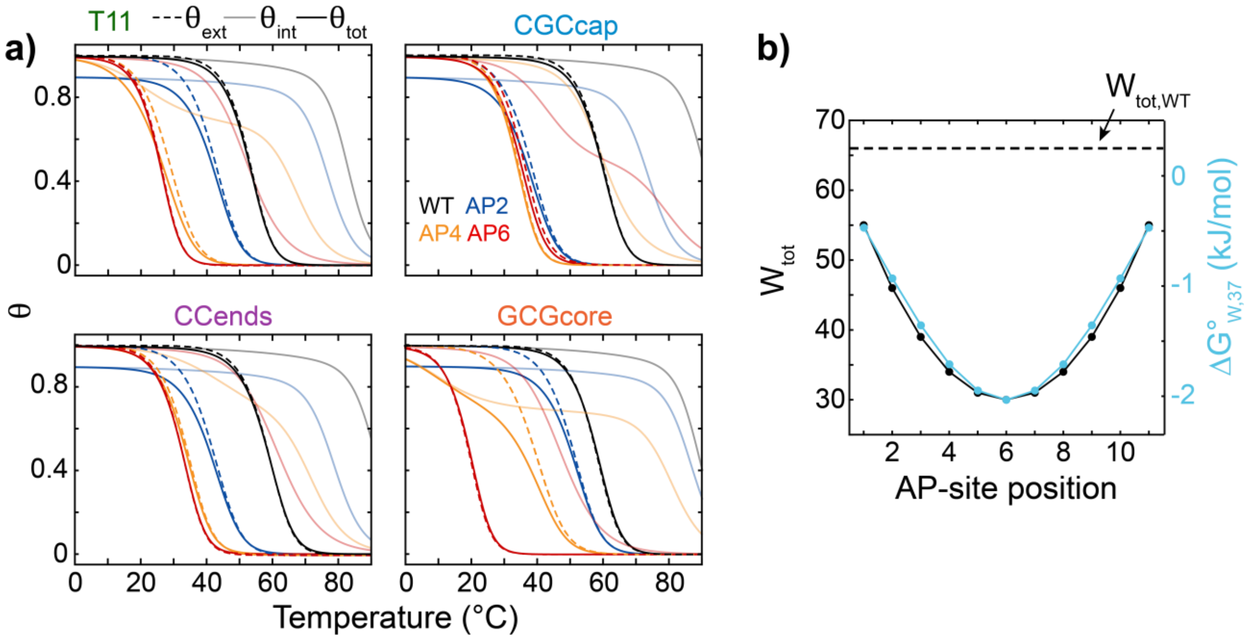


**Figure S6. Melting profiles from two-stretch helix-coil model. (a)** External ($\theta_{ext}$, dashed line), internal ($\theta_{int}$, light solid line) and total base-pair ($\theta_{tot} = \theta_{int}\theta_{ext}$, dark solid line) melting curves from eqs. S15, S24, S27. A single nucleation parameter ($\sigma = \sigma_1 = \sigma_2$) of $10^{-4}$ was used for each sequence. An entropic scaling factor (κ) of 2.1, 2.3, 2.3, 2.6 was applied to T11, CGCcap, CCends, and GCGcore sequences, respectively, to match the WT sequence $T_m$ to the experimental value. **(b)** Number of possible in-register base-pairing configurations ($W_{tot}$) as a function of AP-site position from Eqs. S28-S29. The dashed line indicates $W_{tot}$ for the WT duplex. $W_{tot}$ decreases as an AP site is shifted further from the duplex termini. The free-energy penalty for the reduction in $W_{tot}$ at 37 °C, $\Delta G^0_{W,37} = -RTln(W_{tot,AP}/W_{tot,WT})$, is shown in blue.

##### S2.3 Base-pairing probability distributions and free-energy profiles

The probability distribution across $n_{bp}$ can be determined for WT sequences using the statistical weight for a given $n_{bp}$ divided by the internal partition function. The probability of having $n_{bp}$ base pairs is scaled by $\theta_{ext}$ to account for full strand dissociation.

$$P(N, n_{np}) = \frac{\sigma(N - n_{bp} + 1)\langle s \rangle^{n_{bp}}}{Z_{\text{int},D}(N)} \theta_{ext} \text{ for } n_{bp} \geq 1 \quad \text{(S30a)}$$

$$P(N,0) = 1 - \sum_{n_{bp}=1}^{N} P(N, n_{bp}) \quad \text{(S30b)}$$

In section S2.2, sequences containing an AP site were described in terms of independent stretches with $n_{bp.1}$ and $n_{bp,2}$ base pairs. To determine the probability distribution across $n_{bp} = n_{bp,1} + n_{bp,2}$, we must take into account all possible combinations of $n_{bp,1}$ and $n_{bp,2}$ that sum to $n_{bp}$.

$$P(N, n_{bp}) = \theta_{ext} \sum_{n_{bp,1}=x}^{y} P_1(N_1, n_{bp,1}) P_2(N_2, n_{bp} - n_{bp,1}) \text{ for } n_{bp} \geq 1 \quad \text{(S31)}$$

$$\text{where, } P_m(N_m, n_{bp,m}) = \frac{\sigma_m(N_m - n_{bp,m} + 1)\langle s_i \rangle^{n_{bp,i}}}{Z_{\text{int},Dm}(N_m)} \text{ for } m = 1 \text{ or } 2$$

and

| | $x$ | $y$ |
|---|---|---|
| $n_{bp} \leq N_1$ & $n_{bp} \leq N_2$ | $0$ | $n_{bp}$ |
| $n_{bp} \leq N_1$ & $n_{bp} > N_2$ | $n_{bp} - N_2$ | $n_{bp}$ |
| $n_{bp} > N_1$ & $n_{bp} \leq N_2$ | $0$ | $N_1$ |
| $n_{bp} > N_1$ & $n_{bp} > N_2$ | $n_{bp} - N_2$ | $N_1$ |

$n_{bp}$ probability distributions are shown for each sequence at 37 °C in Fig. S7.

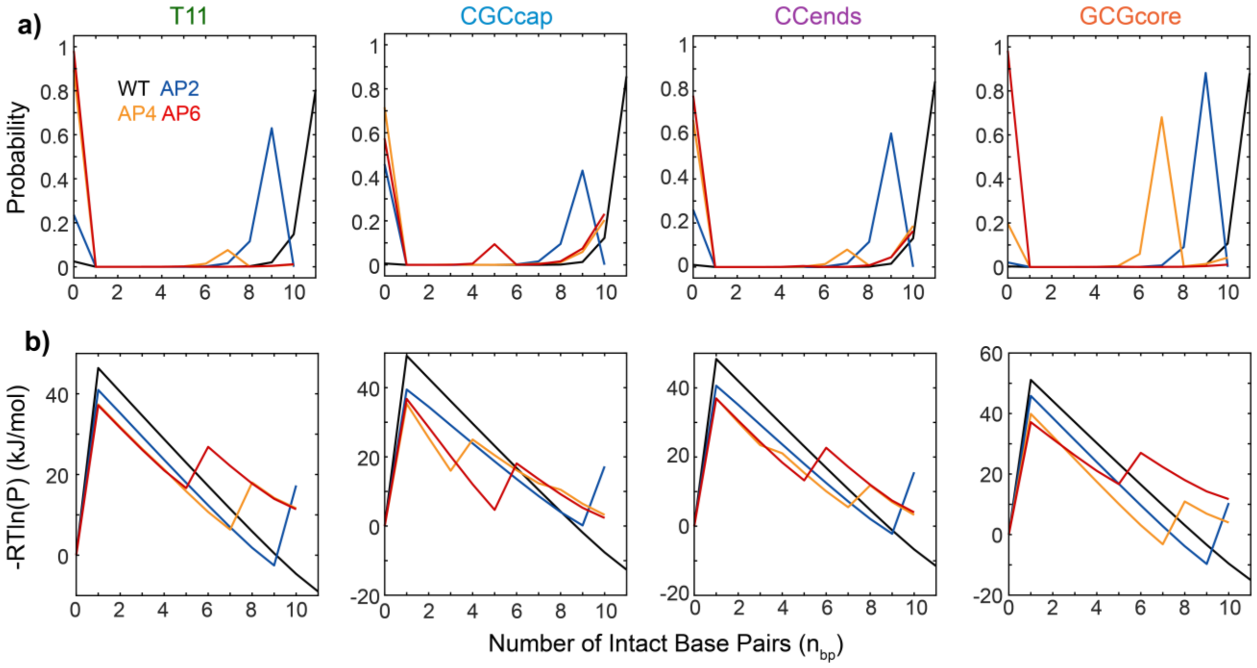


**Figure S7. Probability distributions and free-energy profiles from two-stretch helix-coil model. (a)** Probability distributions as a function of the number of intact base pairs ($n_{bp}$) at 37 °C determined from the helix-coil model (eqs. S30-S31). $\sigma$, $\sigma_1$, and $\sigma_2$ were set to $10^{-4}$ for all sequences and κ was set to 2.1, 2.3, 2.3, and 2.6 for T11, CGCcap, CCends, and GCGcore sequences, respectively. **(b)** Free-energy profiles determined from the probability distributions (P) in **(a)** by $-RTln(P)$, where R is the ideal gas constant.

### S3. Free-energy surfaces and thermodynamics from 3SPN.2 simulations

#### S3.1 Comparison of duplex dehybridization thermodynamics from 3SPN.2 and FTIR

Duplex melting curves were determined the 3SPN.2 MD simulations with WTMetaD using the same method as previously reported.(1) The values $T_m$ and $\Delta G^{\circ}_{h37}$ obtained from 3SPN.2 MD simulations and FTIR temperature series show strong positive correlation, but the those for $\Delta H^{\circ}_{h}$ and the melting curve derivative at $T_m$, $(d\theta/dT)_{T_m}$, are in quite poor agreement and not well correlated (Fig. S9). Discrepancies between 3SPN.2 and FTIR $\Delta H^{\circ}_{h}$ and $(d\theta/dT)_{T_m}$ are particularly severe for AP sequences. In general, sequences containing G:C base pairs are predicted by the 3SPN.2 simulations to have lower $T_m$ and sharper melting transitions than those measured by FTIR while T11 sequences tend to have higher $T_m$ and broader melting curves. 3SPN.2 has primarily been parameterized at low salt concentrations (<200 mM) and may not predict $T_m$ as accurately at the 600 mM salt concentration used for simulations and experiments in this work.(15) Discrepancies in the duplex melting transition sharpness are less surprising because 3SPN.2 is primarily parameterized against $T_m$ and $\Delta G^{\circ}_{h37}$,(15) but the width of the melting transition is largely determined by $\Delta H^{\circ}_{h}$ and $\Delta S^{\circ}_{h}$. As such, the experimental comparisons lend confidence to the computational predictions of $T_m$ and $\Delta G^{\circ}_{h37}$, but the width of the experimental melting transitions are not well reproduced by the computational model.

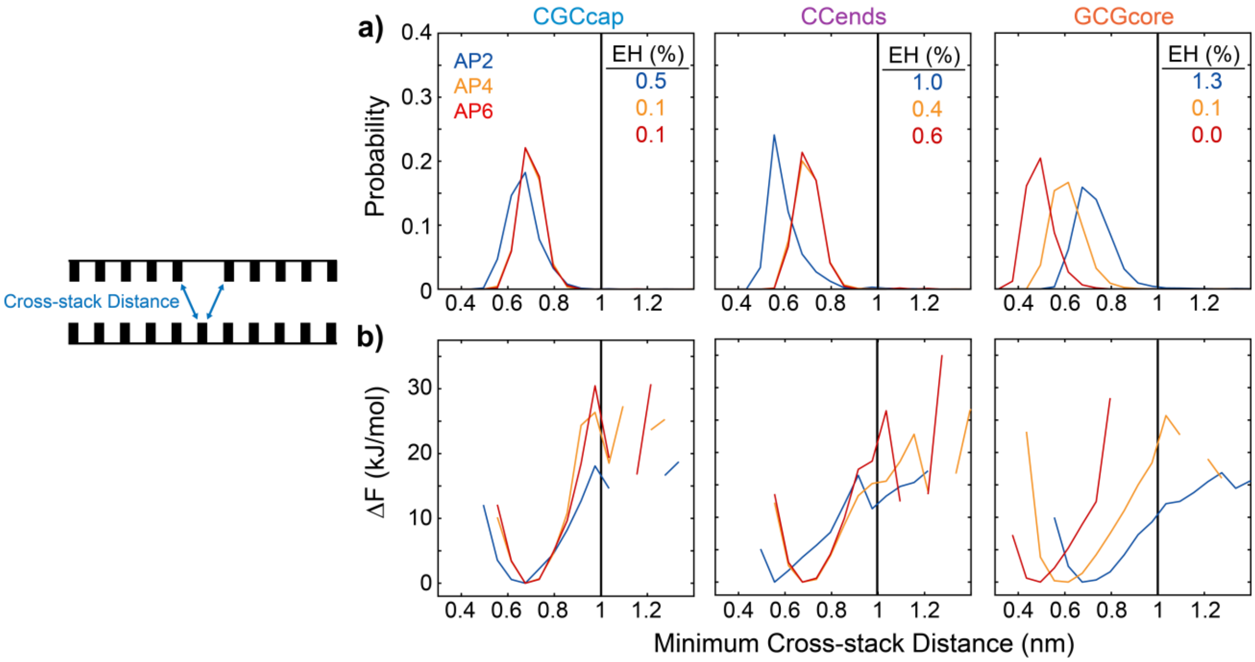


**Figure S8. Configuration of the nucleobase opposite the AP-site in 3SPN.2 MD simulations (a)** Probability distributions for the minimum cross-stacking distance between the nucleobase opposite the AP site and nucleobases adjacent to the AP site (schematic shown on the left). Populations include all configurations from WTMetaD simulation with $n_{bp} \geq 9$ for AP4 and AP6 sequences and $n_{bp} = 10$ for AP2 sequences. **(b)** Corresponding FEPs plotted in ΔF relative to the intrahelical free-energy minimum. The nucleobase opposite the AP site is extrahelical when the minimum cross-stacking distance is greater than 1 nm, and these configurations are minor for all sequences but AP2 with EH populations (EH) of 0.5 – 1.3%.

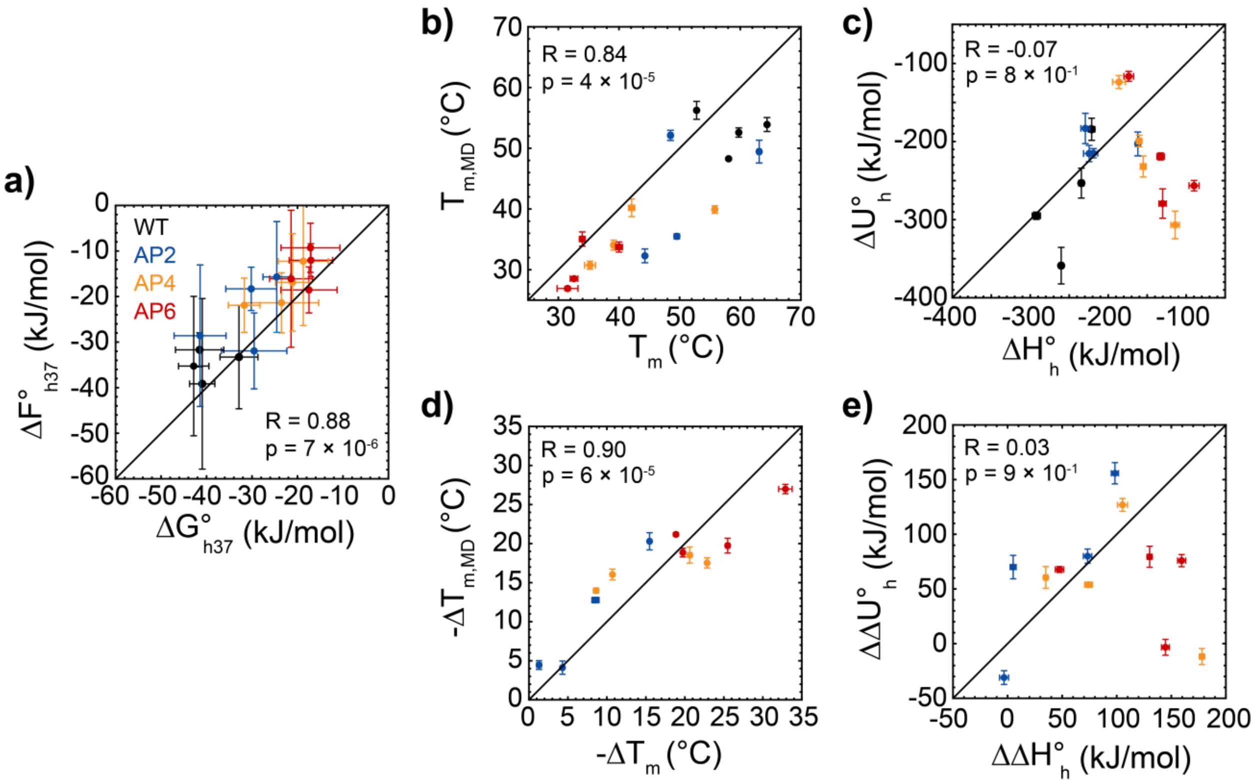


**Figure S9. Comparison between duplex hybridization thermodynamic parameters extracted from FTIR and 3SPN.2 MD simulations.** Scatter plots are shown for **(a)** $\Delta G^{\circ}_{h37}$ vs. $\Delta F^{\circ}_{h37}$, **(b)** $T_m$vs. $T_{m,MD}$, **(c)** $\Delta H^{\circ}_{h}$ vs. $\Delta U^{\circ}_{h}$, **(d)** $\Delta T_m$ vs. $\Delta T_{m,MD}$ and **(e)** $\Delta\Delta H^{\circ}_{h}$ vs. $\Delta\Delta U^{\circ}_{h}$. In each plot, the experimental FTIR results are presented on the x-axis and the computational 3SPN.2 results are presented on the y-axis. Error bars are propagated from 95% confidence intervals from two-state fits to FTIR and 3SPN.2 melting curves. Pearson correlation coefficient (R) and p value is reported for each plot. Black solid lines are shown along the diagonal. Good correlation is found between experimental and model dehybridization free energy and melting temperature, but correlation for dehybridization enthalpy/internal energy is poor.

#### S3.2 Base-pairing contributions to metastable FEP minima

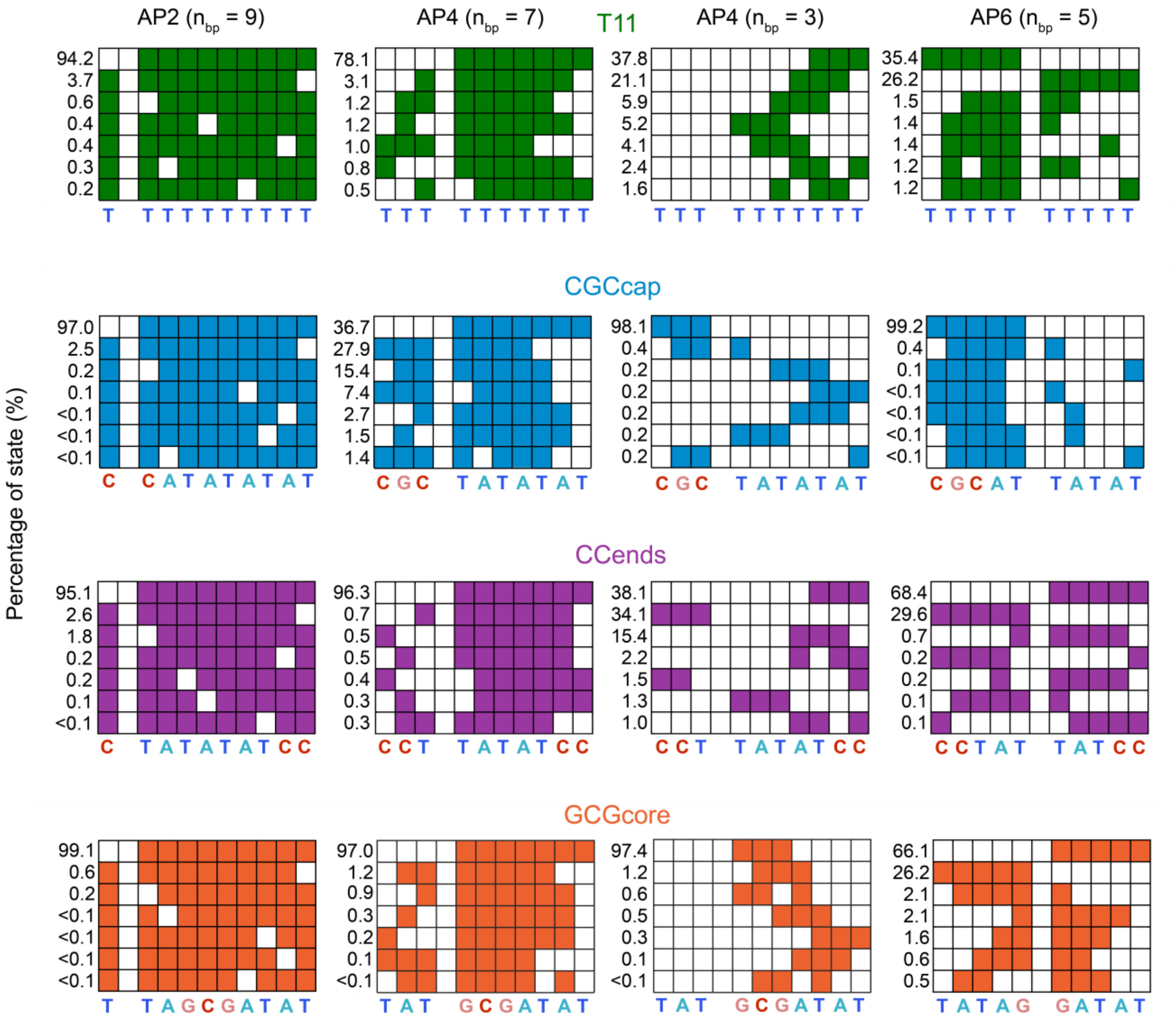


**Figure S10. Base-pair configurations contributing to local FEP minima.** Seven most probable fully in-register base-pairing configurations contributing to the (left) $n_{bp}$ = 9 state of AP2 sequences, (left-center) $n_{bp}$ = 7 state of AP4 sequences, (right-center) $n_{bp}$ = 3 state of AP4 sequences, and (right) $n_{bp}$ = 5 state of AP6 sequences from FEPs in Fig. 4a. The probability of a given configuration (%) is listed on the left. Local free-energy minima along $n_{bp}$ are dominated by configurations that only contain intact base pairs on one side of the AP-site.

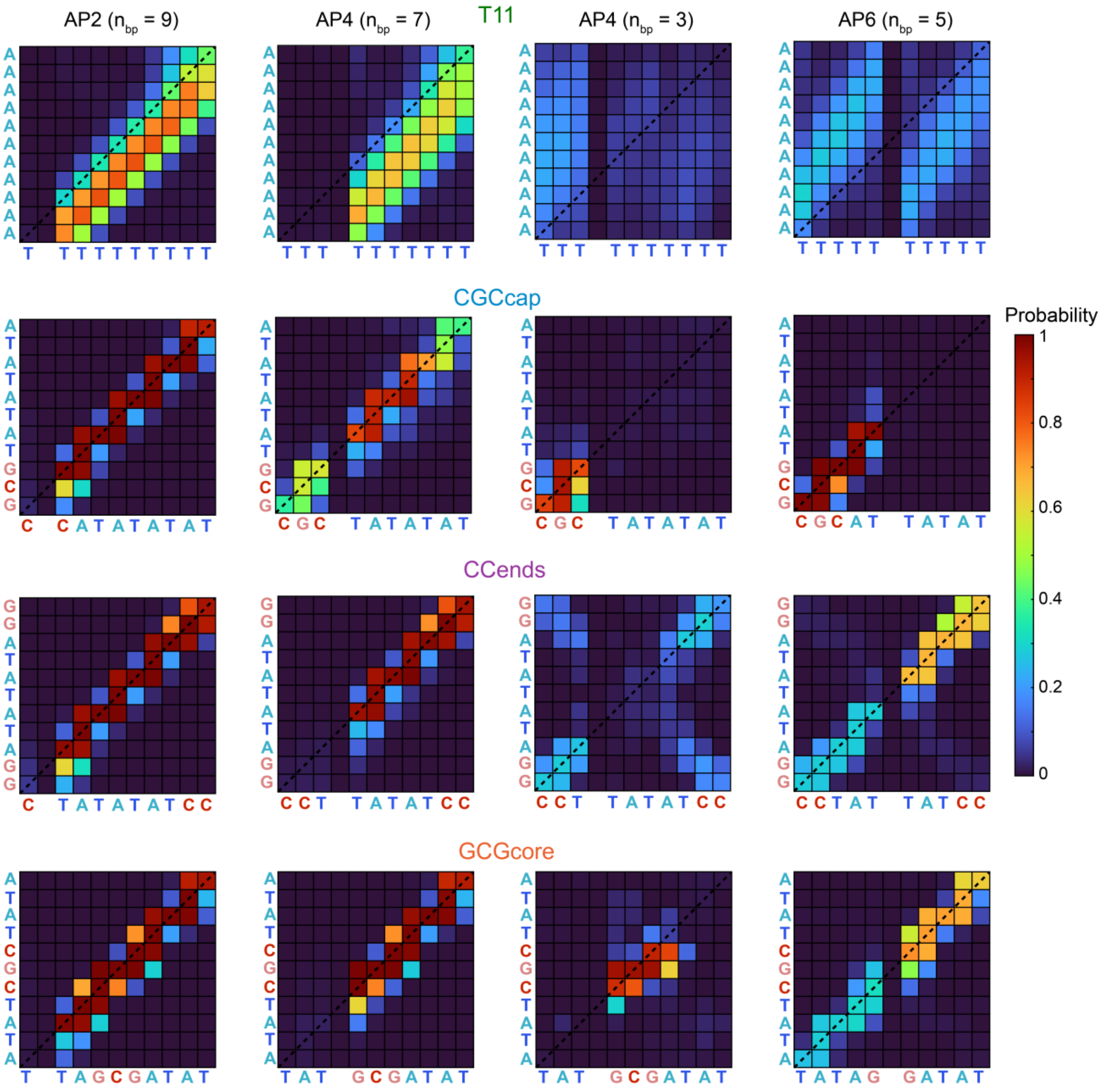


**Figure S11. Contact maps of metastable states.** Contact maps indicating the probability for each nucleobase intermolecular distance to be below 0.7 nm in the (left) $n_{bp}$ = 9 state of AP2 sequences, (left-center) $n_{bp}$ = 7 state of AP4 sequences, (right-center) $n_{bp}$ = 3 state of AP4 sequences, and (right) $n_{bp}$ = 5 state of AP6 sequences from FEPs in Fig. 4a. Squares along the diagonal (dashed line) correspond to probabilities for in-register contacts. Metastable states for CGCcap, CCends, and GCGcore sequences are dominated by in-register base pairing whereas configurations for metastable states in T11 sequences are primarily out-of-register.

##### S3.3 Temperature-dependent FEPs and duplex populations

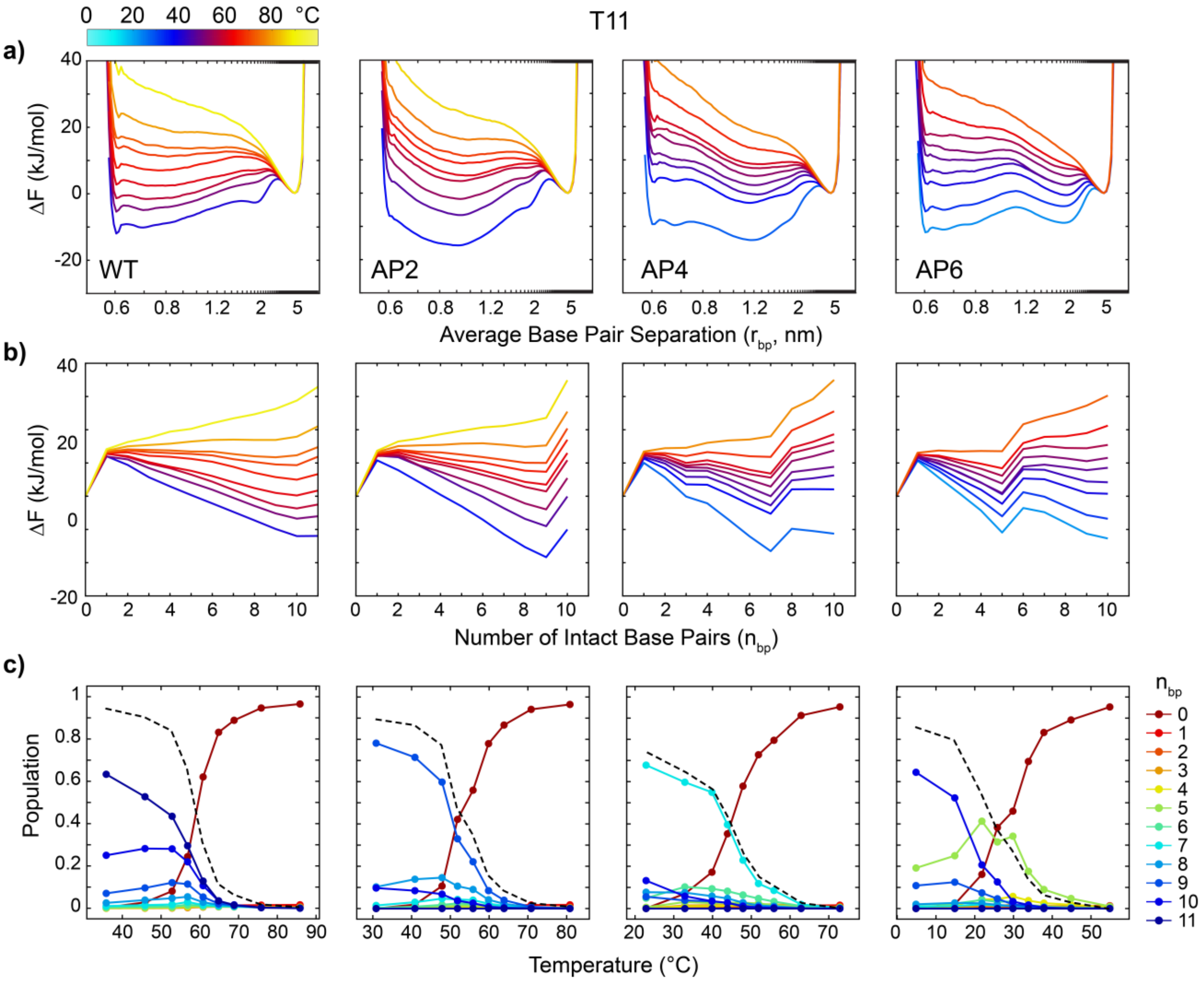


**Figure S12. Temperature-dependent free-energy profiles for T11 sequences determined from 3SPN.2 simulations with WTMetaD.** Free-energy profiles are shown as a function of **(a)** the average separation of all in-register base pairs ($r_{bp}$) and **(b)** the number of intact base pairs ($n_{bp}$) at temperatures from $T_m$ – 20 to $T_m$ + 20 °C. The location of the free-energy minimum at $r_{bp}$ = 5.0 nm in panel (a) is attributed to the 8.5 nm side length of the periodic cubic simulation box that limits the maximum separation of the DNA strands to 7.28 nm. The same considerations apply for FEPs of other sequences shown in Figs. S13 – S15. FEPs are aligned such that the arbitrary zero of free energy is at the free-energy minimum of the single-strand state. **(c)** Population of duplex species with different $n_{bp}$ as a function of simulation temperature. Black dashed lines indicate the total fraction of intact base pairs (equivalent to $\theta_{tot}$).

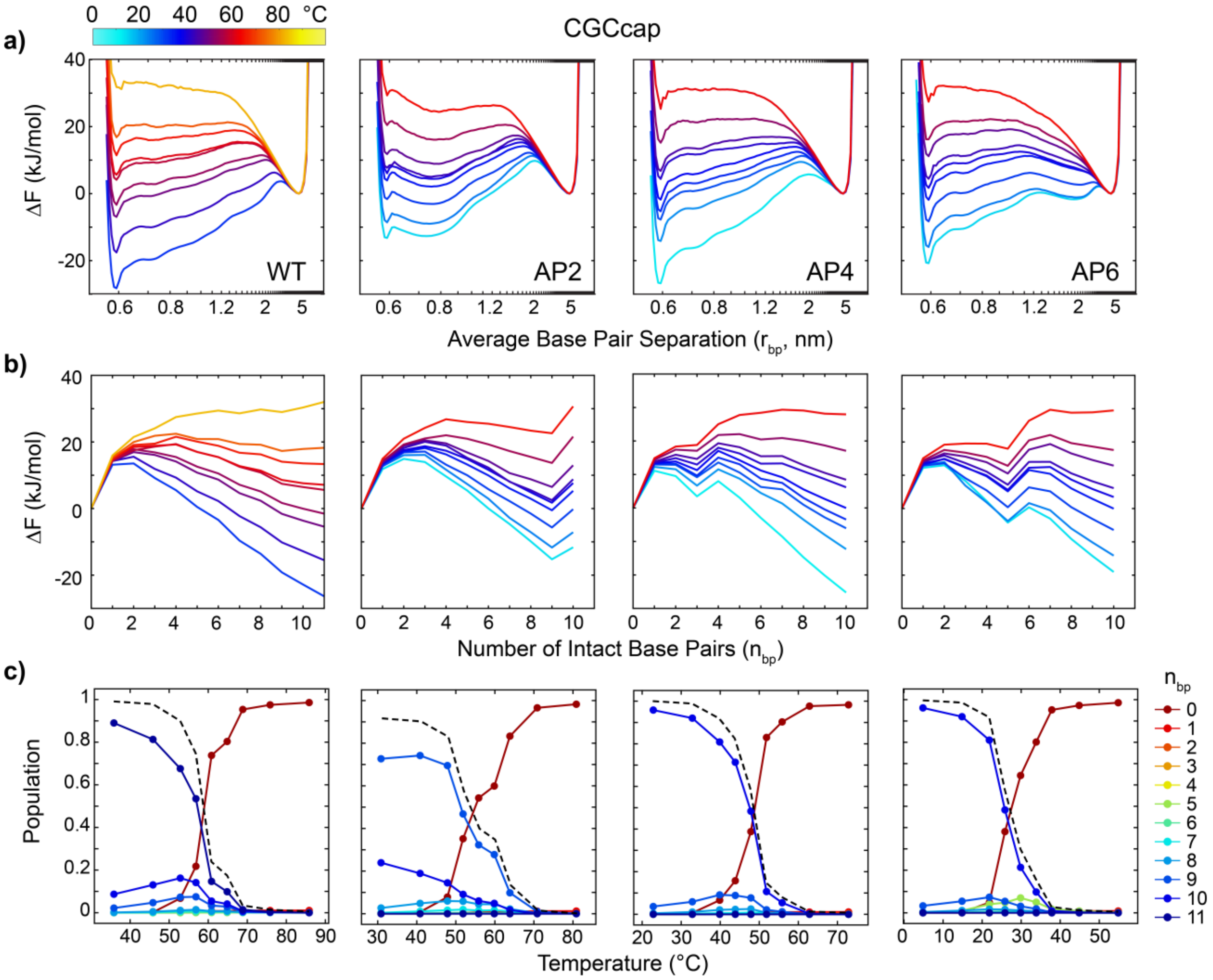


**Figure S13. Temperature-dependent free-energy profiles for CGCcap sequences determined from 3SPN.2 simulations with WTMetaD.** Free-energy profiles are shown as a function of (a) $r_{bp}$ and (b) $n_{bp}$ at temperatures from $T_m$ – 20 to $T_m$ + 20 °C. (c) Population of duplex species with different $n_{bp}$ as a function of simulation temperature. Black dashed lines indicate the total fraction of intact base pairs (equivalent to $\theta_{tot}$).

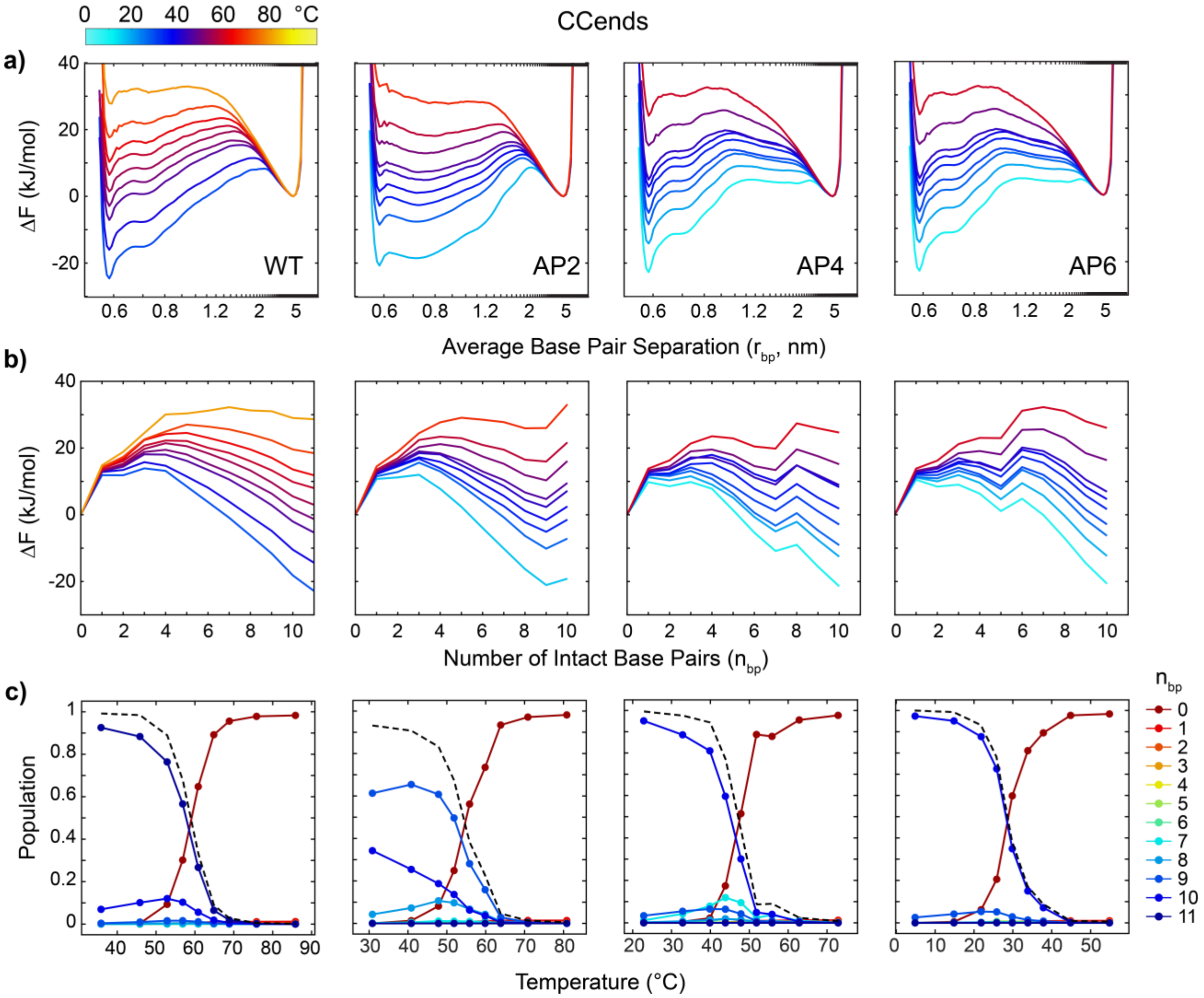


**Figure S14. Temperature-dependent free-energy profiles for CCends sequences determined from 3SPN.2 simulations with WTMetaD.** Free-energy profiles are shown as a function of (a) $r_{bp}$ and (b) $n_{bp}$ at temperatures from $T_m$ – 20 to $T_m$ + 20 °C. (c) Population of duplex species with different $n_{bp}$ as a function of simulation temperature. Black dashed lines indicate the total fraction of intact base pairs (equivalent to $\theta_{tot}$).

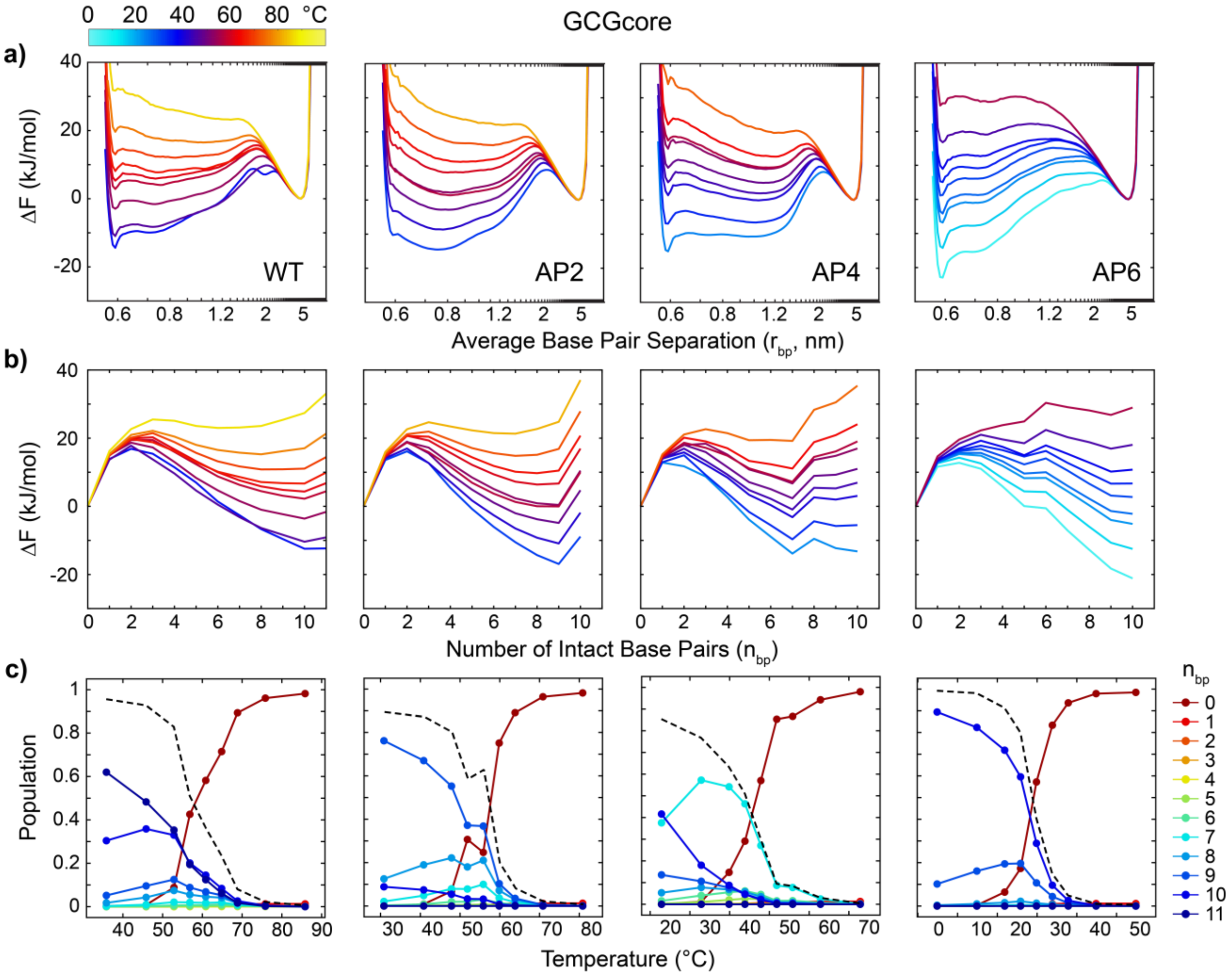


**Figure S15. Temperature-dependent free-energy profiles for GCGcore sequences determined from 3SPN.2 simulations with WTMetaD.** Free-energy profiles are shown as a function of (a) $r_{bp}$ and (b) $n_{bp}$ at temperatures from $T_m$ – 20 to $T_m$ + 20 °C. (c) Population of duplex species with different $n_{bp}$ as a function of simulation temperature. Black dashed lines indicate the total fraction of intact base pairs (equivalent to $\theta_{tot}$).

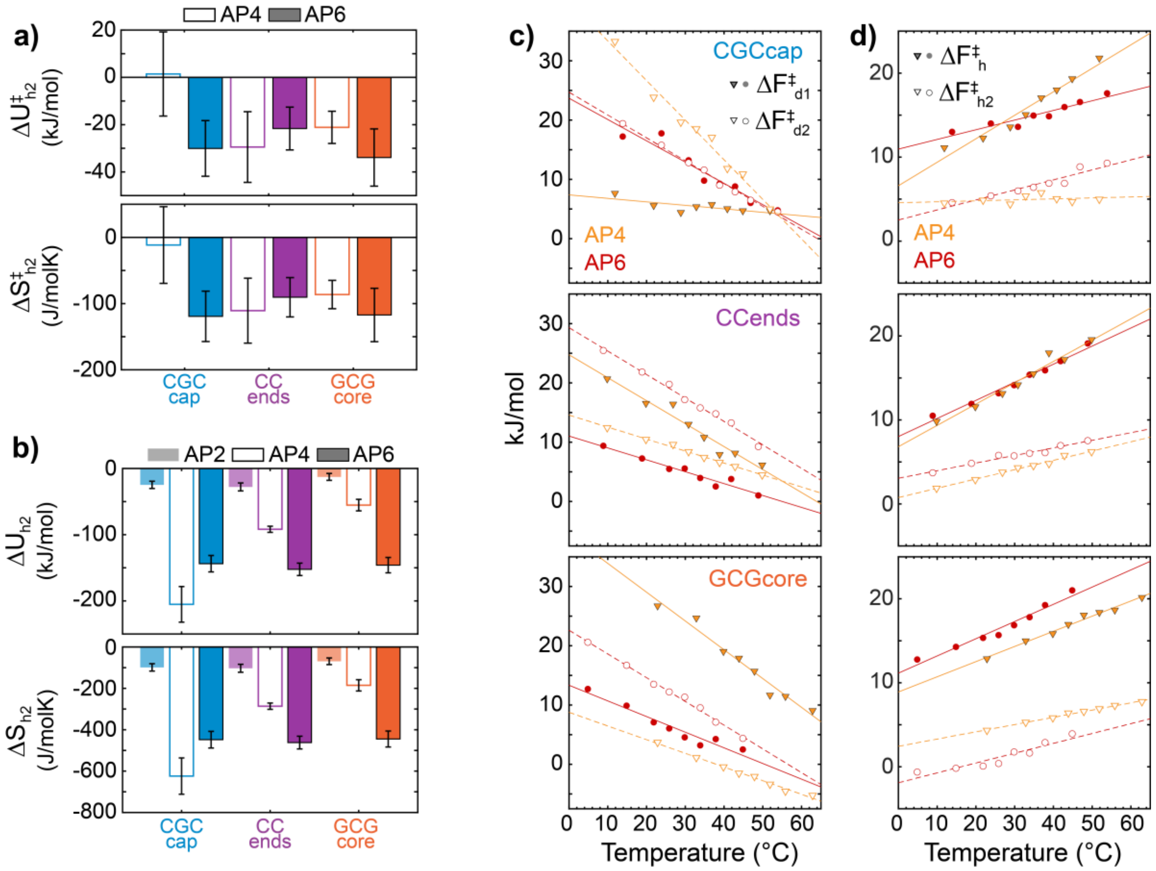


**Figure S16. Temperature dependence of $\Delta F_{h2}$, $\Delta F^{\ddagger}_{h2}$, $\Delta F^{\ddagger}_{d1}$, and $\Delta F^{\ddagger}_{d2}$ from FEPs. (a)** Internal-energy ($\Delta U^{\ddagger}_{h2}$) and entropic ($\Delta S^{\ddagger}_{h2}$) barriers for nucleation of the second base-pair segment determined from linear fits of the temperature-dependence of $\Delta F^{\ddagger}_{h2}$ (Fig. 4b), $\Delta F^{\ddagger}_{h2} = \Delta U^{\ddagger}_{h2} - T\Delta S^{\ddagger}_{h2}$. **(b)** Internal-energy ($\Delta U_{h2}$) and entropy ($\Delta S_{h2}$) change for hybridization of the second base-pair segment determined from linear fits of the temperature dependence of $\Delta F_{h2}$ (Fig. 4c). Error bars correspond to 95% confidence intervals from fits. **(c)** Temperature-dependence of $\Delta F^{\ddagger}_{d1}$ (closed symbols and solid lines) and $\Delta F^{\ddagger}_{d2}$ (open symbols and dashes lines) for AP4 and AP6 sequences**.** **(d)** Temperature-dependence of $\Delta F^{\ddagger}_{h}$ (closed symbols and solid lines) and $\Delta F^{\ddagger}_{h2}$ (open symbols and dashed lines) for AP4 and AP6 sequences**.**

#### S4. Temperature-jump IR spectroscopy

##### S4.1 Temperature-dependent t-HDVE data

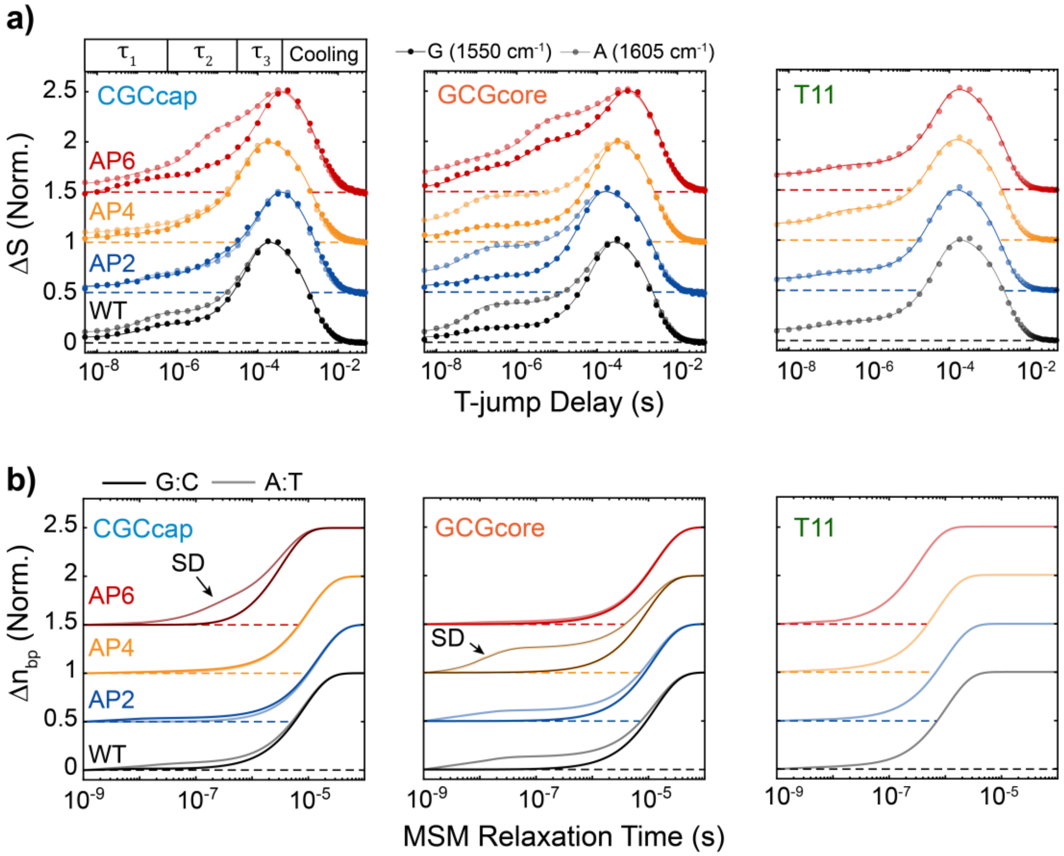


**Figure S17. T-jump IR and MSM T-jump relaxation profiles of CGCcap, GCGcore, and T11 sequences. (a)** Normalized difference T-jump IR (t-HDVE) time traces plotted as the normalized change relative to the maximum of the initial temperature spectrum, $\Delta S(t) = [S(t) - S(T_i)]/\max[S(T_i)]$. Time traces probed at 1550 cm$^{-1}$ (dark) report on changes in G:C base pairing and those at 1605 cm$^{-1}$ (light) report on changes in A:T base pairing. T-jumps are performed from approximately $T_m$–15°C to $T_m$ for each sequence. Traces are shifted vertically with respect to one another, and dashed lines indicate respective baselines. Solid lines correspond to three- or four-component fits from global lifetime analysis (Section S5.2). **(b)** Markov state model (MSM) T-jump simulations of the normalized change in intact G:C (dark) and A:T (light) base-pairs, $\Delta n_{bp}(t) = [n_{bp}(t) - n_{bp}(T_i)]/[n_{bp}(T_f) - n_{bp}(T_i)]$, from $T_{m,MD}$–15°C to $T_{m,MD}$. Changes corresponding to segment-dehybridization (SD) are indicated. Traces are shifted vertically with respect to one another, and dashed lines indicate respective baselines. Black/gray solid lines for CGCcap-AP6 and GCGcore-AP4 correspond to fits to a sum of two exponential components.

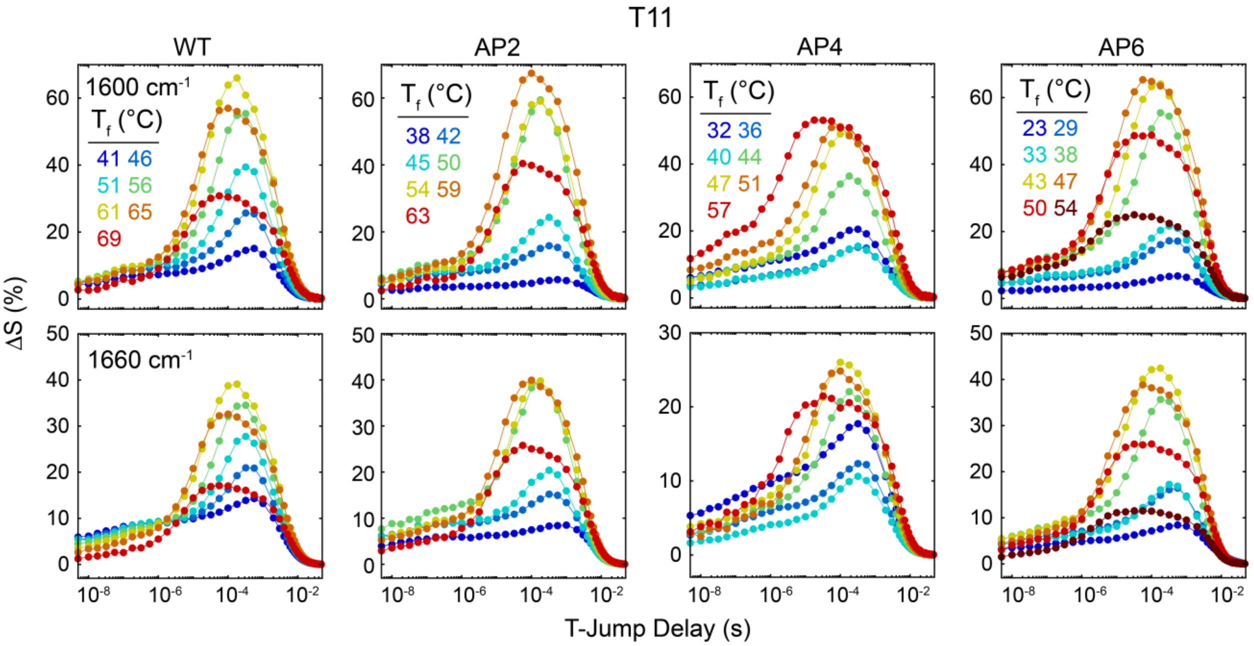


**Figure S18. Temperature-dependent T-jump IR data for T11 sequences.** Temperature-dependent t-HDVE time traces for T11 sequences shown at probe frequencies of 1600 (adenine ring mode) and 1660 $cm^{-1}$ (thymine carbonyl mode).

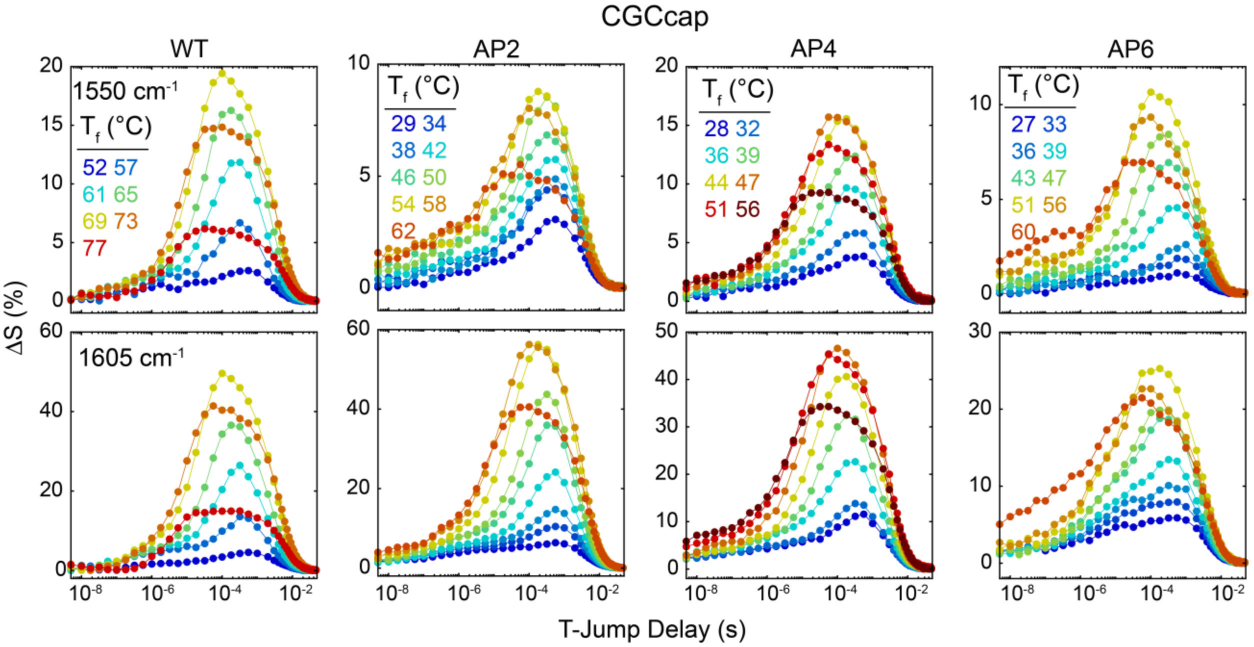


**Figure S19. Temperature-dependent T-jump IR data for CGCcap sequences.** Temperature-dependent t-HDVE time traces for CGCcap sequences shown at probe frequencies of 1550 and 1605 $cm^{-1}$.

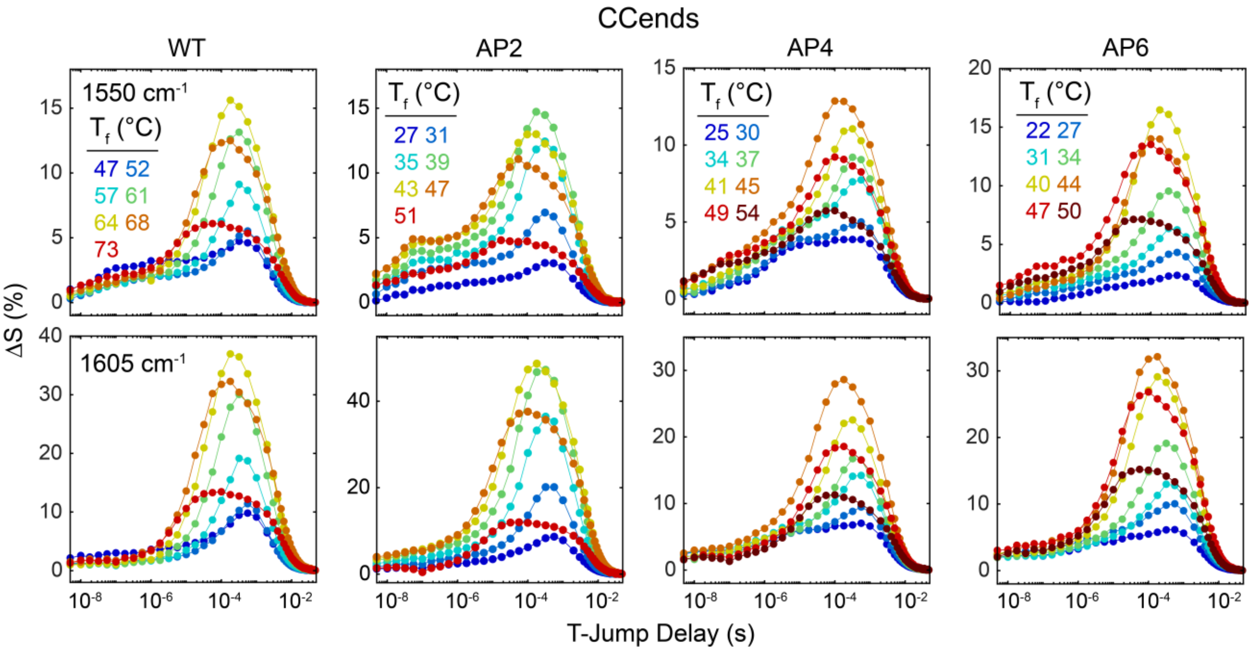


**Figure S20. Temperature-dependent T-jump IR data for CCends sequences.** Temperature-dependent t-HDVE time traces for CCends sequences shown at probe frequencies of 1550 and 1605 $cm^{-1}$.

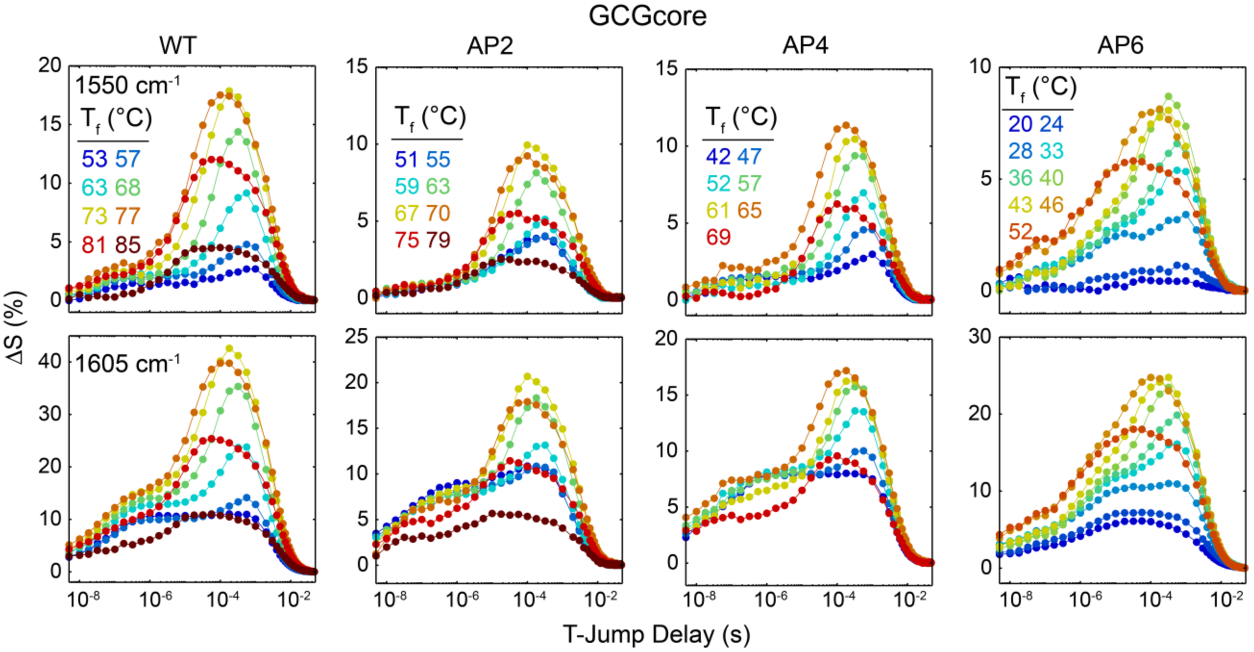


**Figure S21. Temperature-dependent T-jump IR data for GCGcore sequences.** Temperature-dependent t-HDVE time traces for GCGcore sequences shown at probe frequencies of 1550 and 1605 $cm^{-1}$.

##### S4.2 Global lifetime analysis of t-HDVE data

We applied global lifetime analysis to the t-HDVE data ($\psi_{t-HDVE}$) to extract kinetic components, as described previously.(1) Three or four exponential components ($N_{comp}$) are applied based on examination of the rate-domain (Fig. S22) and time-domain data.

$$\psi_{t-HDVE}\left(\omega,\tau_{TJ}\right)=\sum_{n=1}^{N_{comp}} A_n\left(\omega\right)\exp\left(-\frac{\tau_{TJ}}{\tau_n}\right) \quad \text{(S32)}$$

The first two or three components account for the T-jump response while the final component accounts for thermal relaxation and re-hybridization of the sample. Global fitting was performed by minimizing the objective function (*R*) in eq. S33 using a nonlinear least-squares solver where *I* is an identity matrix, *C* is the matrix of time-dependent concentration profiles, and $C^+$ denotes the pseudoinverse of *C*.(16)

$$R=\left\|\left(I-CC^+\right)\psi_{t-HDVE}\right\|^2 \quad \text{(S33)}$$

Figure S23 shows the DAS for each sequence determined from three or four component global lifetime fitting. As expected, the DAS and associated time constants reveal similar information to the rate distributions in Fig. S22. All sequences exhibit a first component with time constant from tens to hundreds of nanoseconds. Previous studies of short oligonucleotide dissociation with T-jump IR demonstrated that the structural dynamics on this timescale can primarily be assigned to terminal base-pair fraying.(11,17-20) The first component spectra in Fig. S23 (DAS 1) are consistent with this assignment and more details are discussed in Section S3.5. A second component (DAS 2) with a time constant ranging from hundreds of nanoseconds to many microseconds is only needed for CCends-AP4, CGCcap-AP4, CGCcap-AP6, CCends-AP6, and GCGcore-AP6. The final T-jump component (DAS 3) occurs from a few to hundreds of microseconds depending on the temperature and sequence and corresponds to full-strand dissociation and association.

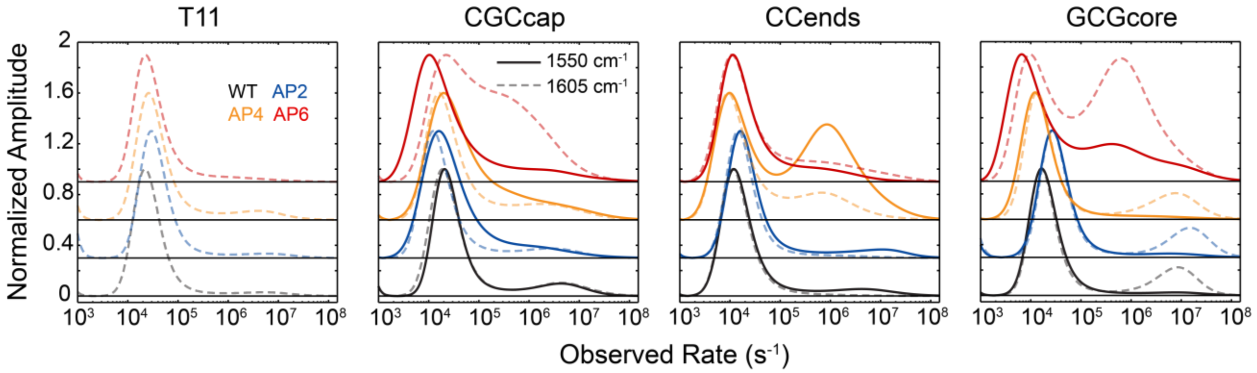


**Figure S22. T-jump IR rate distributions.** Rate-domain traces of t-HDVE data at 1550 cm$^{-1}$ (dark solid line) and 1605 cm$^{-1}$ (light dashed line) determined from an inverse-Laplace transform of the time-domain data using a maximum entropy method (MEM-iLT).(20,21) Data correspond to the same temperature conditions as in Figs. 5 and S17. The number of maxima in the rate traces were used to inform global lifetime fitting of the t-HDVE data.

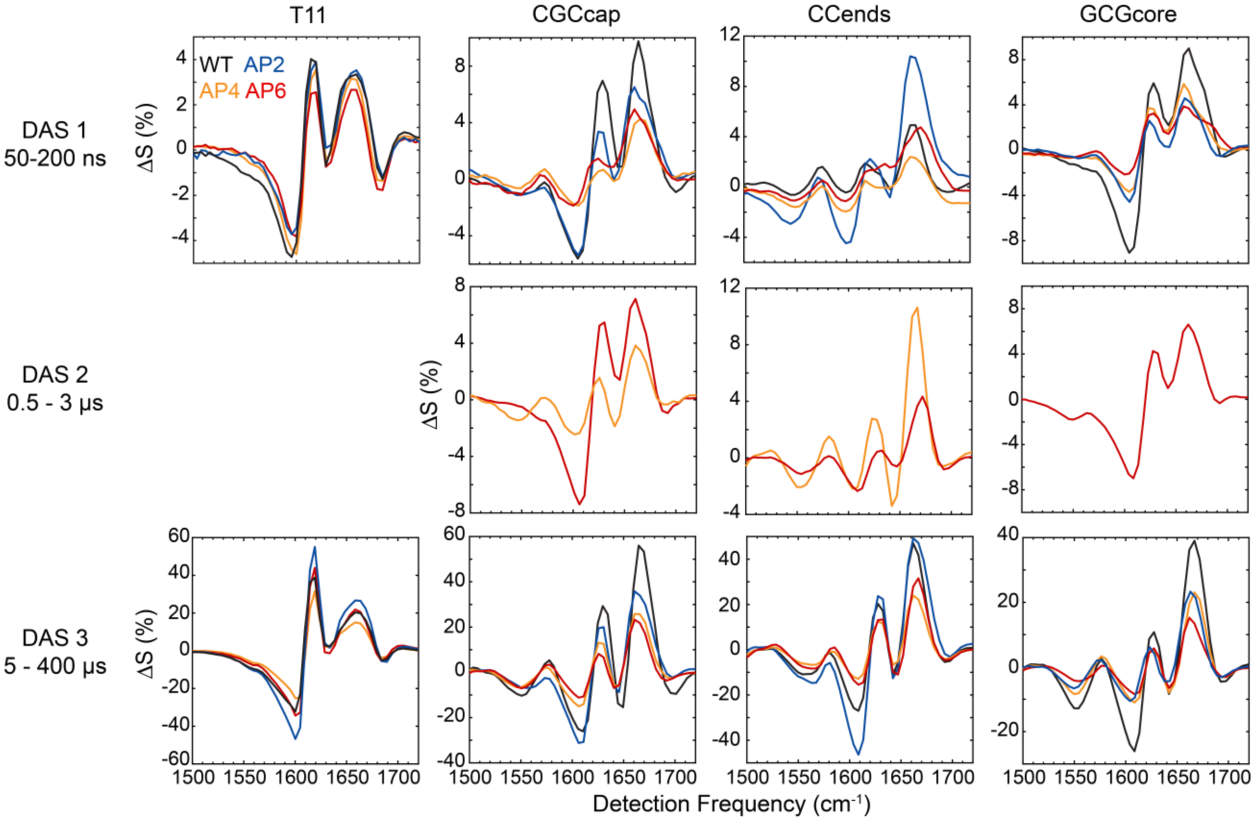


**Figure S23. Decay-associated spectra (DAS) determined from global lifetime analysis of t-HDVE data for all sequences.** DAS corresponding to thermal relaxation are not shown. DAS are shown for a T-jump with $T_f$ near the respective duplex melting temperature. (top) The first spectral component (DAS 1) has a time constant ranging from 50-200 ns and is observed for most sequences. (middle) An intermediate component (DAS 2) is only observed for CGCcap-AP4, CGCcap-AP6, CCends-AP4, CCends-AP6, and GCGcore-AP6 at select temperatures with a time constant ranging from 500 ns to 3 μs. (bottom) DAS 3 corresponds to complete duplex dissociation and is observed for all sequences on a timescale ranging from a 5 to 400 μs.

##### S4.3 Temperature-dependent trends in $\tau_1$ response

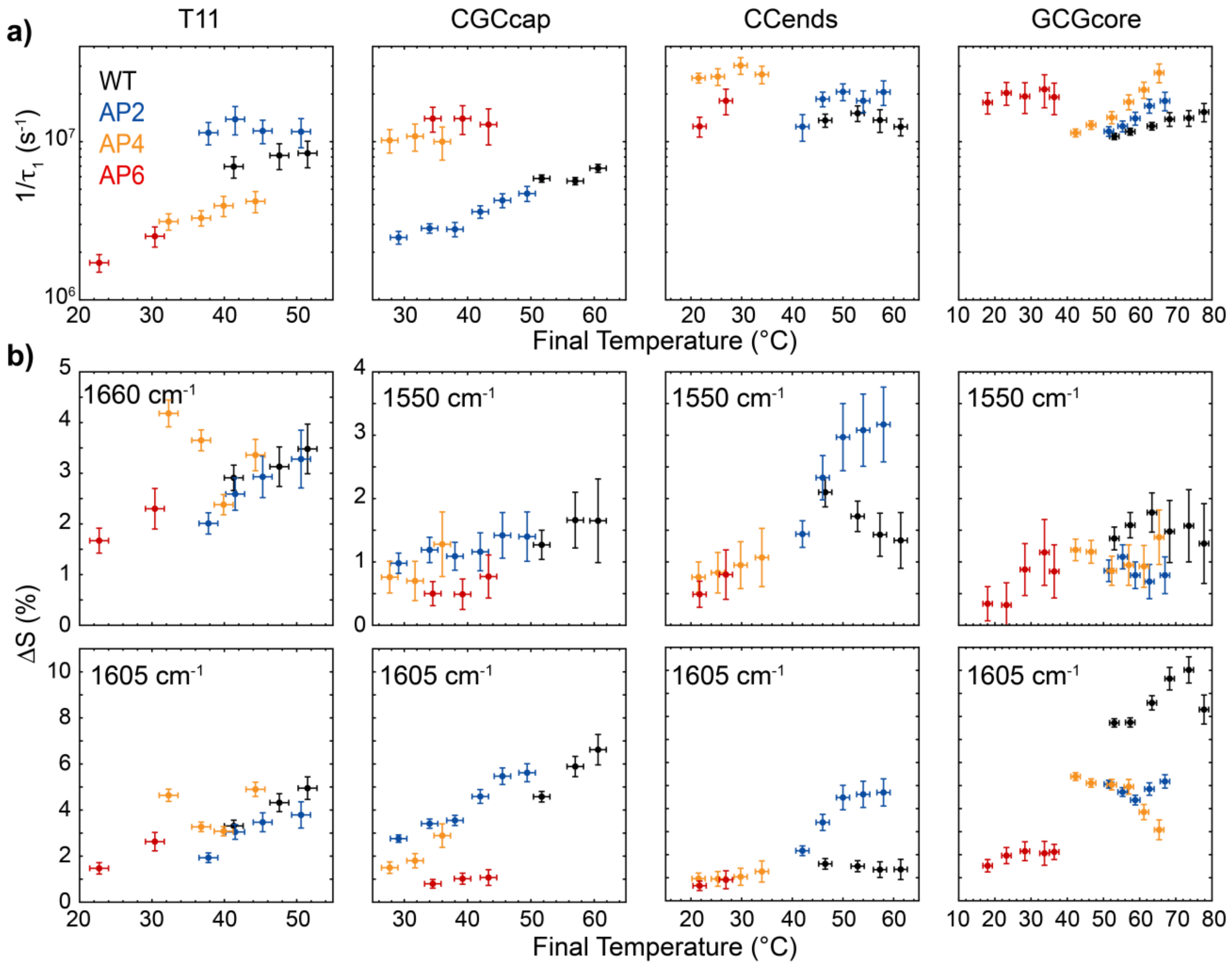


**Figure S24. Rate and amplitude of $\tau_1$ response. (a)** Rate of first component ($1/\tau_1$) for all sequences determined from global lifetime fitting of t-HDVE data. Signal amplitude during $\tau_1$ is negligible at many measurement temperatures. **(b)** DAS1 amplitude at denoted frequencies for all sequences. Vertical error bars correspond to 95% confidence intervals from global lifetime fitting and horizontal error bars are the standard deviation in T-jump magnitude. For CGCcap, CCends, and GCGcore sequences, 1550 $cm^{-1}$ primarily reports on changes in G:C base pairing and 1605 reports on changes in A:T base pairing.

##### S4.4 Dehybridization and hybridization kinetics from $\tau_3$ response

The temperature-dependence of the T-jump response during $\tau_3$ (Fig. S25) provides insight into how an AP-site alters the kinetics of duplex dissociation and association. Dissociation ($k_d$) and hybridization ($k_h$) rate constants were derived by applying a two-state relaxation kinetics model to the observed rates ($1/\tau_3$) (eq. S34). FTIR melting curves (Figs. 1 and 3) were used to determine single-strand concentrations ($[S_1]$ and $[S_2]$) and the dissociation constant ($K_d$) at each $T_f$.(22)

$$1/\tau_3 = k_d + k_h\left(\left[S_1\right]_{T_f} + \left[S_2\right]_{T_f}\right) \tag{S34}$$

$$1/\tau_3 = \frac{k_d K_{D,T_f}}{K_{d,T_f} + \left[S_1\right]_{T_f} + \left[S_2\right]_{T_f}}, \text{ where } K_{d,T_f} = \frac{k_{d,T_f}}{k_{h,T_f}} \tag{S35}$$

Although many of the AP4 and AP6 sequences exhibit three-state relaxation kinetics overall, we can still apply a two-state treatment to $1/\tau_3$. In this case, reports on the average dehybridization kinetics between the duplex ensemble after terminal fraying and segment-dehybridization during $\tau_1$ and $\tau_2$ and the single-strand ensemble. The temperature-dependence of $k_d$ and $k_h$ (Fig. S26) can be described by a Kramers-like expression in the high-friction limit to extract enthalpic ($\Delta H^\ddagger$) and entropic ($\Delta S^\ddagger$) barriers where the temperature-dependence of the $D_2O$ solvent viscosity ($\eta$) is taken into account.(23)

$$k_{d/h}\left(T\right) = \frac{k_b T}{\hbar}\frac{\eta_{37}}{\eta\left(T\right)}\exp\left(-\frac{\Delta H^{\dagger}_{d/h} - T\Delta S^{\dagger}_{d/h}}{RT}\right) \tag{S36}$$

In eq. S36, we assume a transmission coefficient of 1, yet the true value is likely much smaller.(24) Overestimating the transmission coefficient leads to a reduction in $\Delta S^\ddagger$ and therefore an increase in free-energy barrier ($\Delta G^\ddagger$) for both dehybridization and hybridization. However, assuming the transmission coefficient or speed limit of dehybridization and hybridization are independent of sequence, the relative trends of $\Delta S^\ddagger$ and $\Delta G^\ddagger$ across sequence and AP-site position are still informative.

Consistent with previous studies, we find a large enthalpic dehybridization barrier ($\Delta H_d^\ddagger$ >150 kJ/mol, Fig. S26) for all sequences. Incorporation of an AP-site has negligible effect or reduces $\Delta H_d^\ddagger$ by as much as 120 kJ/mol ($\Delta\Delta H_d^\ddagger$). $\Delta G_{d37}^\ddagger$ and $\Delta G_{d37}^\circ$ are well correlated with a Pearson correlation coefficient (R) of 0.98 and a slope of 0.90 ± 0.08 (Fig. S41e). Slopes of $\Delta G_{d37}^\ddagger$ vs. $\Delta G_{d37}^\circ$ near 1 have been observed across canonical oligonucleotides of various sequence or environmental conditions,(25,26) which suggests that the change in $\Delta G_{d37}^\ddagger$ is primarily due to destabilization of the duplex rather than a change in transition state energy. In contrast, the hybridization free-energy barrier ($\Delta G^{\ddagger}_{h37}$) shows poor correlation with $\Delta G^{\circ}_{d37}$ (R = 0.35) and a slope of -0.10 ± 0.08. $\Delta G^{\ddagger}_{h37}$ only varies by ~6 kJ/mol across all sequences, such that all values are within the error of each other. $\Delta G^{\ddagger}_{h37}$ is generally more weakly dependent on sequence or modification than $\Delta G^{\ddagger}_{d37}$,(27,28) but has been shown to increase slightly from an AP site.(29) Most of our T-jump measurements are performed at temperatures where $k_d$ dominates $1/\tau_3$ such that our determination of $k_h$ is rather indirect and model-dependent. Therefore, a more direct measurement of $k_h$ is required to accurately assess small changes from an AP-site.

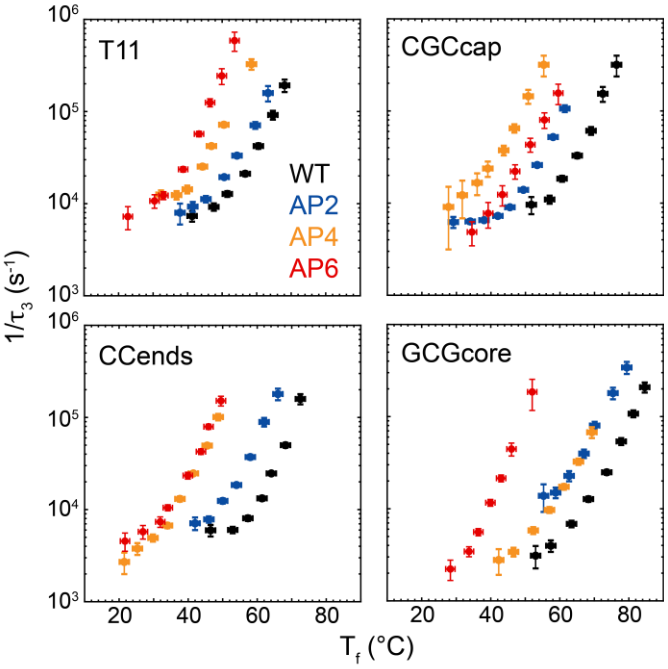


**Figure S25. Rate of $\tau_3$ response.** Observed rates for full-duplex dissociation ($1/\tau_3$) determined from global lifetime fitting of the t-HDVE data as a function of $T_f$. Vertical error bars indicate 95% confidence intervals from global fits and horizontal error bars correspond to the measured standard deviation in T-jump magnitude.

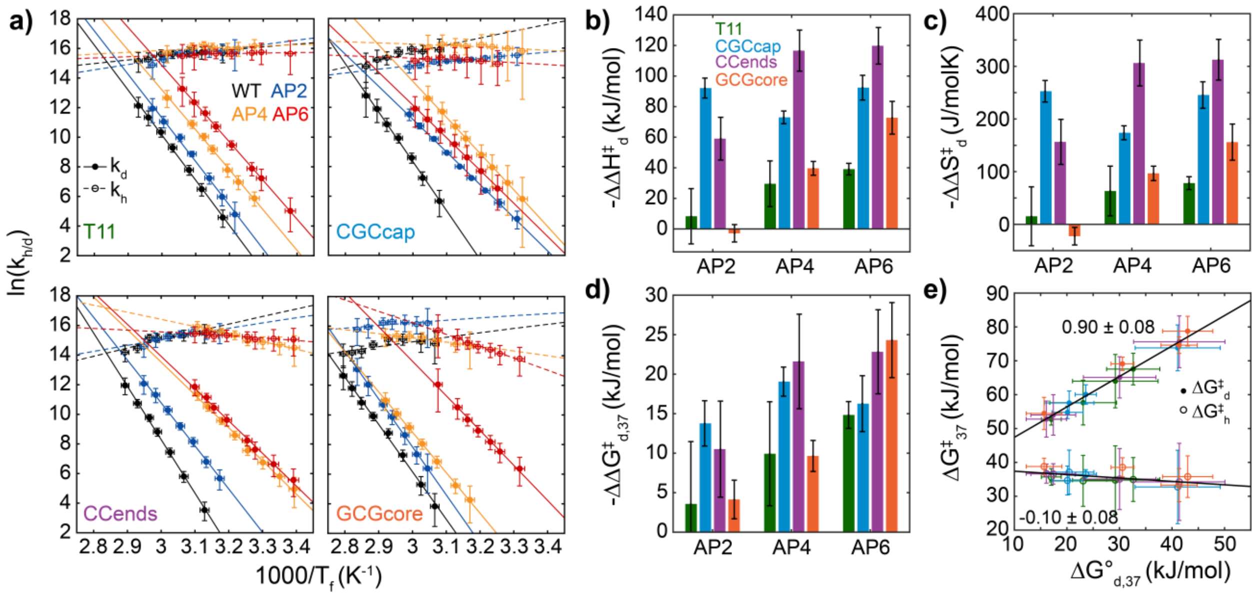


**Figure S26. Kinetic of dehybridization and association. (a)** Temperature-dependence of dehybridization ($k_d$, filled circles) and hybridization ($k_h$, open circles) rate constants determined from two-state analysis of ($1/\tau_3$, Fig. S25). Temperature-trends are fit to eq. S36 (solid and dashed lines). Vertical error bars indicate 95% confidence intervals propagated from global fits and FTIR melting curves and horizontal error bars correspond to the measured standard deviation in T-jump magnitude. Change relative to WT sequence in **(b)** enthalpic ($\Delta\Delta H_d^{\ddagger}$), **(c)** entropic ($\Delta\Delta S_d^{\ddagger}$), and **(d)** dehybridization free-energy barrier at 37 °C ($\Delta\Delta G_{d,37}^{\ddagger}$) extracted from fits to eq. S36 in **(a).** Error bars correspond to 95% confidence intervals from fit. **(e)** Scatter plots for $\Delta\Delta G_{d,37}^{\circ}$ vs. $\Delta G_{d,37}^{\ddagger}$ (closed circles) and $\Delta G_{d,37}^{\circ}$ vs. the free-energy hybridization barrier ($\Delta G_{h,37}^{\ddagger}$, open circles) for all sequences with linear fits (black solid lines).

#### S4.5 Kinetics of segment-dehybridization

The fraction of A:T and G:C character in the $\tau_2$ response ($\tau_1$ for GCGcore-AP4) were determined using the same method described for AP6 sequences previously.(1) First, we calculate the percentage of total dehybridization amplitude change ($P_{G/A}$) that occurs during $\tau_2$ using the ratio of DAS2 to DAS2+DAS3 amplitude at 1550 cm$^{-1}$ for G:C and 1605 cm$^{-1}$ for A:T.

$$P_{G/A} = \frac{A_{2,G/A}}{A_{2,G/A} + A_{3,G/A}} \tag{S37}$$

$P_{G/A}$ values for each AP4 and AP6 sequence are shown in Fig. S42b. Then, the A:T and G:C character ($C$) are determined by the ratio of $P_{G/A}$ to the sum of $P_G$ and $P_A$ weighted by the number of A:T ($N_A$) and G:C ($N$-$N_A$) base pairs in the sequence. $N$ is the total number of starting base pairs in the sequence, which is assumed to be 10 for sequences containing an AP-site.

$$C_{A:T} = \frac{N_A P_A}{(N - N_A) P_G + N_A P_A} \tag{S38a}$$

$$C_{G:C} = \frac{(N - N_A) P_G}{(N - N_A) P_G + N_A P_A} = 1 - C_{A:T} \tag{S38b}$$

The values of $C_{A:T}$ are shown in Fig. S27c and are nearly the same, within experimental error, across the temperature ranges measured. The mean value of $C_{A:T}$ and $C_{G:C}$ across measured $T_f$ points are plotted in Fig. 5a.

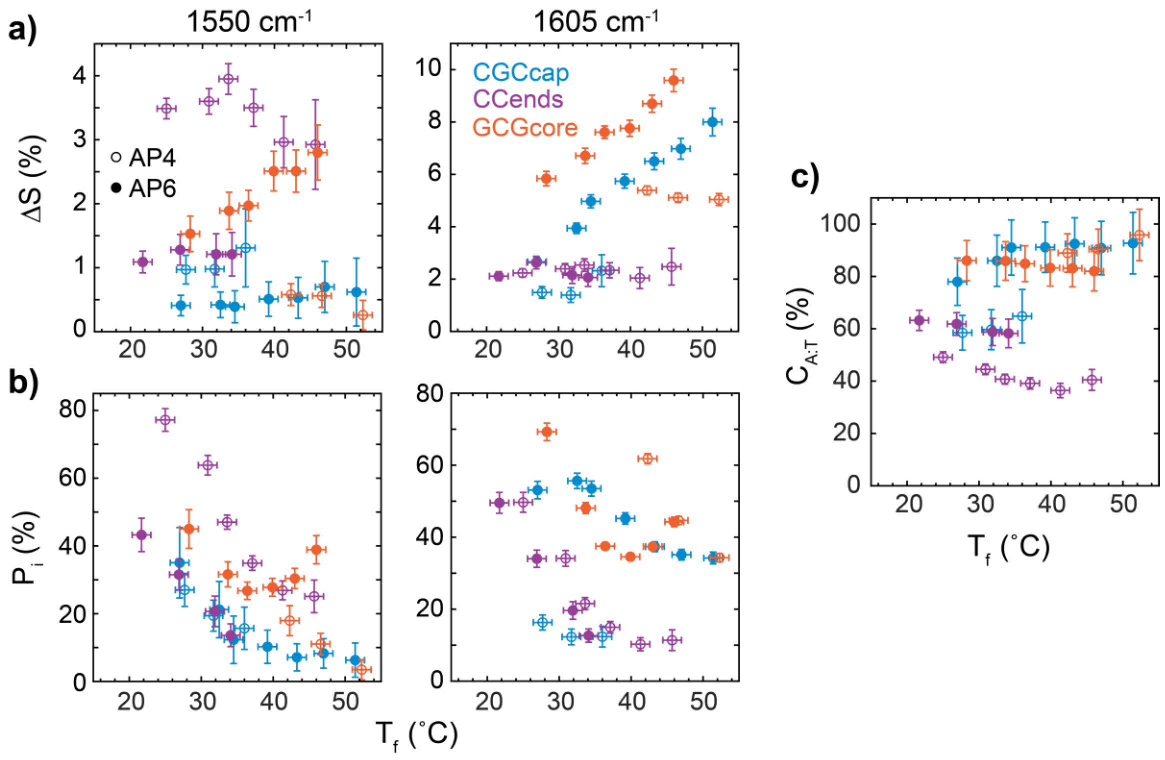


**Figure S27. Determination of A:T/G:C character in $\tau_2$ response. (a)** Amplitudes of second component (DAS 2) probed at 1550 and 1605 cm$^{-1}$ from global lifetime fitting of t-HDVE data for AP4 (open circles) and AP6 (closed circles) sequences as a function of $T_f$. **(b)** Amplitudes in **(a)** divided by the sum of DAS 2 and DAS 3 amplitudes (eq. S37). **(c)** A:T character of $\tau_2$ response at each $T_f$ determined from eq. S40a.

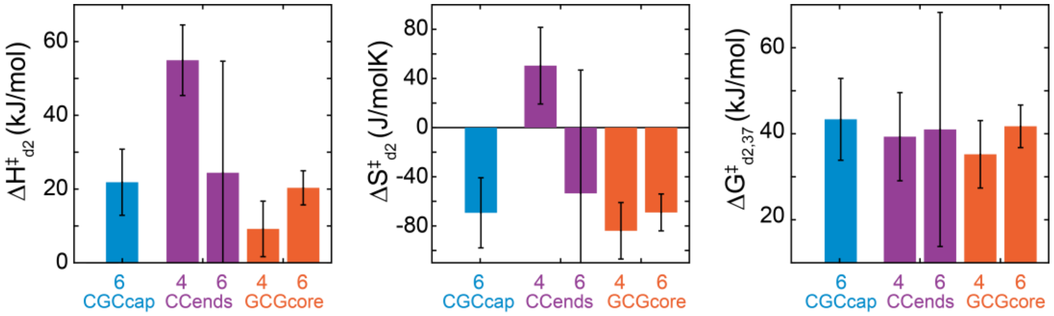


**Figure S28. Kinetics of segment-dehybridization.** Enthalpic ($\Delta H^{\ddagger}_{d2}$), entropic ($\Delta S^{\ddagger}_{d2}$), and free-energy barrier at 37 °C ($\Delta G^{\ddagger}_{d2,37}$) for segment-dehybridization determined from fits of $1/\tau_2$ (Fig. 5c) to eq. S36. Data for GCGcore-AP4 comes from $1/\tau_1$. Error bars correspond to 95% confidence intervals in fit.

**Table S3.** Calculated thermodynamics of segment-dehybridization from Santa Lucia's NN model.(2) Values are corrected to a sodium ion concentration of 600 mM.(8)

| **Sequence** | | **Stretch** | $\Delta H^{\circ}_{d}$ **(kJ/mol)** | $\Delta S^{\circ}_{d}$ **(J/molK)** | $\Delta G^{\circ}_{d,37}$ **(kJ/mol)** |
|---|---|---|---|---|---|
| **CGCcap** | AP4 | 5′-CGC-3′ | 84.9 | 227.7 | 14.3 |
| | | 5′-TATATAT-3′ | 171.1 | 510.1 | 12.9 |
| | AP6 | 5′-CGCAT-3′ | 150.6 | 409.4 | 23.7 |
| | | 5′-TATAT-3′ | 110.9 | 334.3 | 7.2 |
| **CCends** | AP4 | 5′-CCT-3′ | 65.7 | 183.4 | 8.8 |
| | | 5′-TATATCC-3′ | 187.9 | 539.9 | 20.4 |
| | AP6 | 5′-CCTAT-3′ | 125.9 | 359.1 | 14.6 |
| | | 5′-TATCC-3′ | 127.6 | 364.1 | 14.7 |
| **GCGcore** | AP4 | 5′-TAT-3′ | 50.6 | 158.5 | 1.5 |
| | | 5′-GCGATAT-3′ | 200.4 | 554.1 | 28.6 |
| | AP6 | 5′-TATAG-3′ | 113.4 | 336.4 | 9.1 |
| | | 5′-GATAT-3′ | 115.0 | 337.7 | 10.3 |

### S5. Validation of Markov state models from 3SPN.2 MD Simulations

#### S5.1 Construction and validation of state-reversible VAMPnets (SRV) MSMs

**Table S4.** Simulation temperature used for unbiased 3SPN.2 simulations to build MSMs

| Sequence | WT (K) | AP2 (K) | AP4 (K) | AP6 (K) |
|---|---|---|---|---|
| T11 | 338 | 334 | 325 | 320 |
| CGCcap | 330 | 312 | 310 | 312 |
| CCends | 327 | 315 | 308 | 307 |
| GCGcore | 334 | 329 | 321 | 303 |

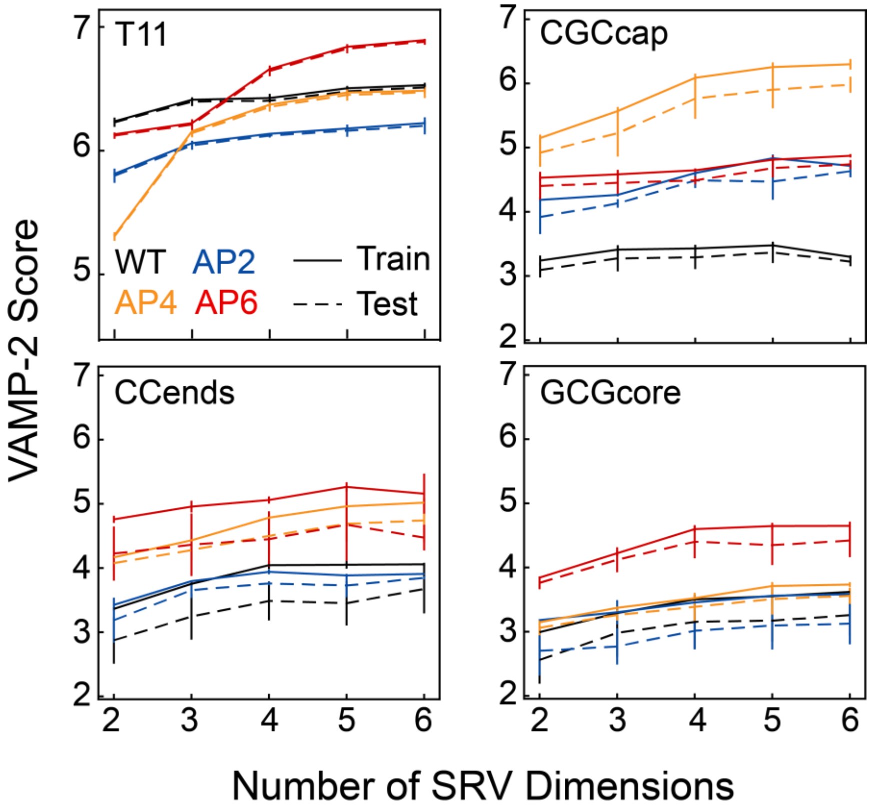


**Figure S29. VAMP-2 scoring with 5-fold cross-validation across all systems as a function of embedding dimensions.** A plateau in the training and testing scores at or before five dimensions indicates a near optimal embedding space across systems. A decrease in VAMP-2 beyond five dimensions for some sequences may be indicative of overfitting. Error bars correspond to the standard deviation computed over the cross-validation partitions.

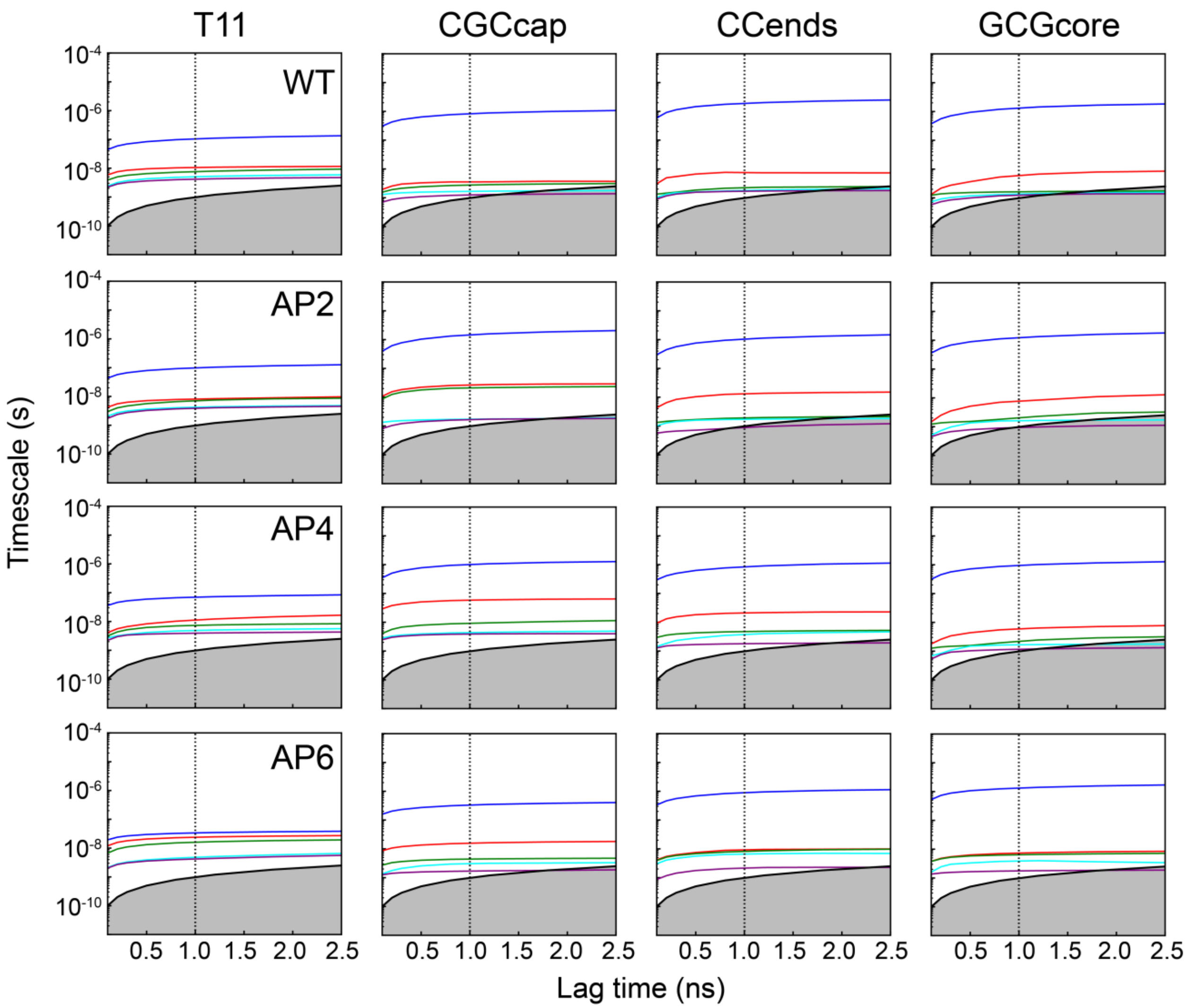


**Figure S30.** Implied timescale convergence plots as a function of lag time for each system after SRV dimensionality reduction and k-means clustering. At a lag time of 1 ns (vertical dashed line), we note that most slow processes have converged and that many fast processes are still resolvable. As noted previously(11) the leading slow mode corresponds to the overall hybridization/dissociation and converges much slower than higher order modes.

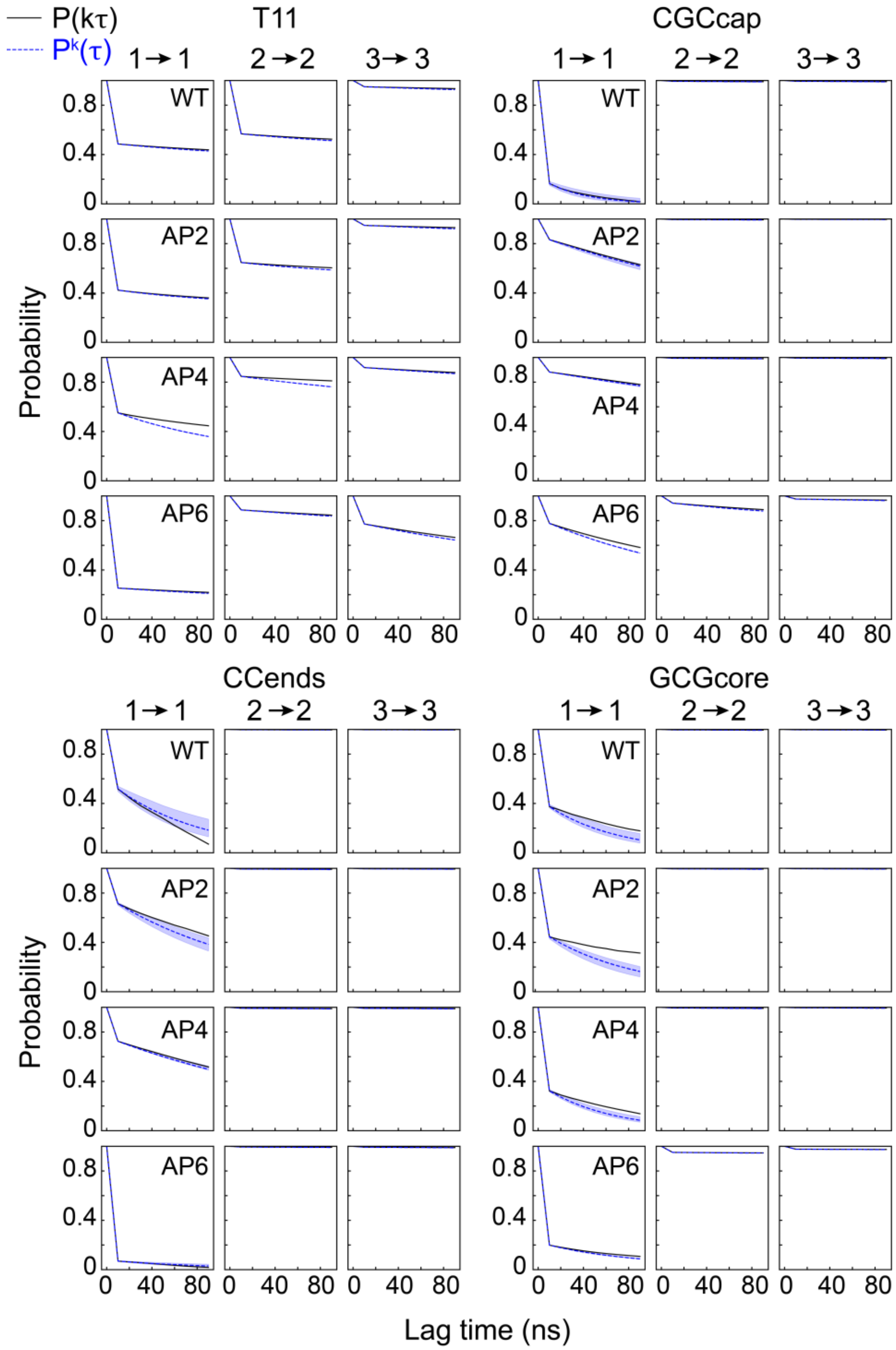


**Figure S31.** Chapman-Kolmogorov tests for SRV-MSMs constructed for all sequences using a lag time of 1 ns.(30) For each system, the self-transition probabilities for 3 PCCA+ sets are shown. Strong agreement between probabilities predicted from k applications of an MSM constructed at a lag time of 1 ns ($P^k(\tau)$, dashed blue line) and those computed from an MSM built at a lag time of k ns ($P(k\tau)$, black solid line) indicate robust Markovian behavior out to at least 100 ns. The blue shaded region indicates the estimated standard deviation in $P^k(\tau)$. Systems that show deviation for the metastable state (1 → 1) tend not to cluster well into coarse-grained states.

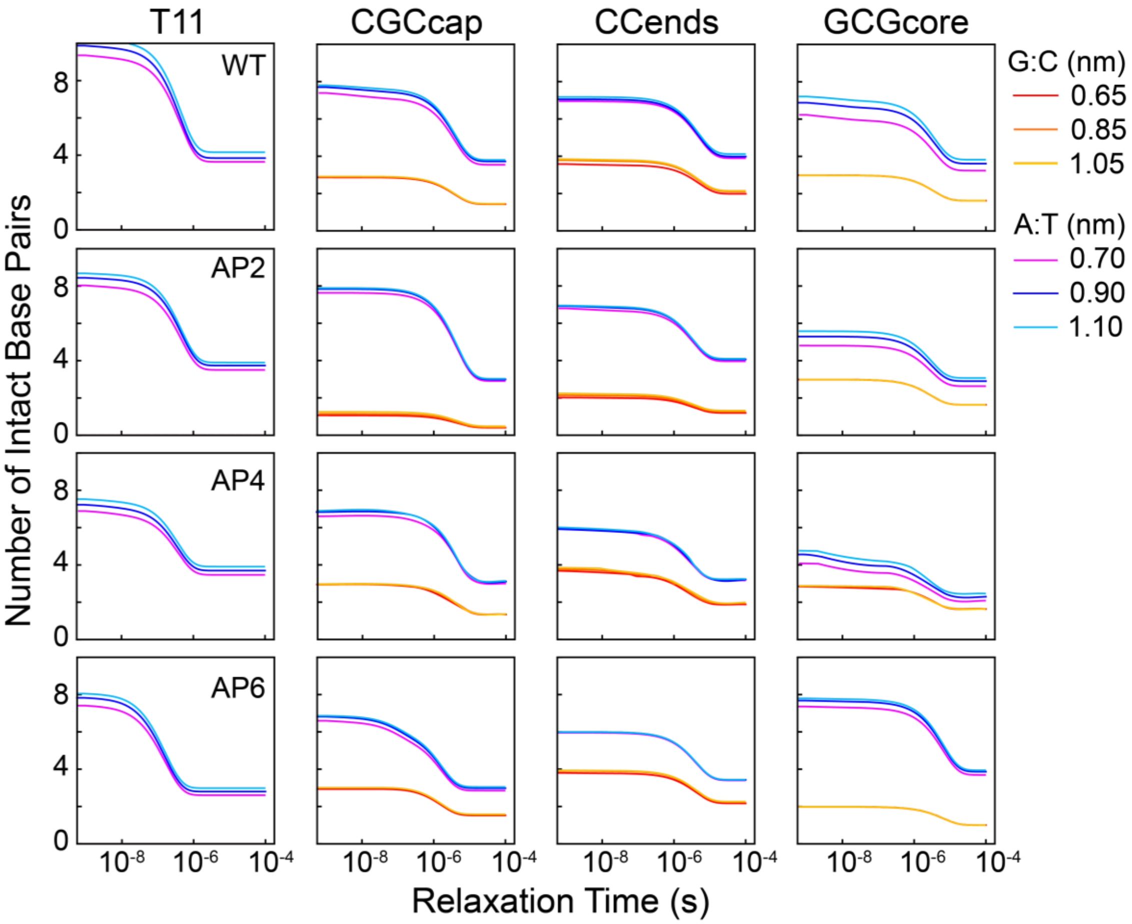


**Figure S32.** Simulated T-jump relaxation profiles in terms of the number of intact A:T (purple-blue) and G:C (red-yellow) base pairs using various base-pairing distance cutoff values. Cutoff is represented as the distance in nm greater than the equilibrium A:T (0.60 nm) or G:C (0.55 nm) center-of-mass distance. All profiles and responses are qualitatively consistent for cutoff values between 0.1 – 0.5, with a systematically lower response over time for lower cutoff values.

#### S6. Assessing deviation from two-state melting behavior

##### S6.1 Evaluating partial dehybridization from total FTIR and 2D IR spectral change

While FTIR temperature series started at 3.8 °C, which is well below the transition midpoint for full strand dehybridization, it is possible that some sequences are partially dehybridized even at 3.8 °C. Both the HC model and 3SPN.2 MD simulations applied in this work predict that AP2 and some AP4 sequences are partially dehybridized at low temperature. We compare the 3.8 °C FTIR spectra and 90 – 3.8 °C difference spectra for each sequence (Fig. S33) to assess the degree of base-pair dissociation at 3.8 °C. Figure S34 compiles the change in absorbance at select frequencies from 3.8 to 90 °C for sequences containing an AP site relative to their respective WT sequence ($\Delta A/\Delta A_{WT}$). The difference in absorbance between 3.8 and 90 °C ($\Delta A$) at the chosen frequencies is dominated by loss of base pairing and stacking. Therefore, we expect that an AP-site will either have a negligible effect on $\Delta A$ ($\Delta A/\Delta A_{WT}$ ~ 100 %) or reduce it ($\Delta A/\Delta A_{WT}$ < 100 %) due to the loss of a chromophore at the specified frequency or elevated dissociation at 3.8 °C. It should be noted that the value of $\Delta A/\Delta A_{WT}$ does not measure the absolute amount of base-pair disruption but instead only that relative to the respective WT sequence. Figure S34 also reports $\Delta A/\Delta A_{WT}$ values determined from 2D IR spectroscopy, which are largely consistent with the FTIR values.

For T11, the largest signal changes occur at the 1625 $cm^{-1}$ adenine ring band and the 1695 $cm^{-1}$ thymine carbonyl band. Regardless of AP-site position, $\Delta A/\Delta A_{WT}$ falls within 85 – 95 % at 1625 $cm^{-1}$, suggesting that the A/A stacking across the duplex is not significantly perturbed. In contrast, T11-AP2 reduces $\Delta A$ at 1695 $cm^{-1}$ and a larger reduction is observed for T11-AP4. The loss of a T nucleobase alone should reduce $\Delta A/\Delta A_{WT}$ by 9%, but instead $\Delta A/\Delta A_{WT}$ is 84% (2D IR value of 92%) for T11-AP2, 60% (68%) for T11-AP4, and 64% (90%) for T11-AP6 are observed. As shown in the 3SPN.2 MD simulations (Fig. 7), it is also possible that an AP site stabilizes out-of-register base-pairing configurations that result in overall loss of A:T base pairing relative to the WT sequence at 3.8 °C. However, $\Delta A/\Delta A_{WT}$ of the A ring mode band at 1625 $cm^{-1}$ is also expected to drop proportionally to the population of out-of-register duplex configurations, yet it is almost the same for all T11 sequences. Therefore, it is more likely that the reduction in $\Delta A$ at 1695 $cm^{-1}$ for T11-AP4 and T11-AP6 result from significant perturbations to nearby T/T stacking interactions rather than out-of-register shifting.

CGCcap, CCends, and GCGcore sequences are more difficult to analyze due to congestion from adenine, thymine, guanine, and cytosine vibrational bands, but the change in guanine ring modes at 1575 $cm^{-1}$ and adenine ring mode at 1625 $cm^{-1}$ offer insight into the amount of A:T and G:C base pairing at 3.8 °C. The spectral change at 1662 $cm^{-1}$ is also shown but is more complicated to interpret due to overlapping changes in G and T carbonyl bands. Starting with CGCcap, we expect $\Delta A/\Delta A_{WT}$ to vary depending on which nucleobase is removed for the AP-site. Removal of a guanine in CGCcap-AP2 leads to $\Delta A/\Delta A_{WT}$ of 72% (2D IR value of 60%) at 1575 $cm^{-1}$ and 105% (103%) at 1625 $cm^{-1}$. There are only three G:C base pairs in CGCcap, so removal of a guanine alone is expected to give a $\Delta A/\Delta A_{WT}$ of 66% at 1575 $cm^{-1}$. Although the FTIR and 2D IR values differ by 10%, the rough agreement between this expected value and experiment suggests that the terminal G:C base pair adjacent to the AP-site is mostly base paired at 3.8 °C. CGCcap-AP4 and CGCcap-AP6 modifications only produce minor changes in $\Delta A/\Delta A_{WT}$ at each of reported frequencies. The reduction of $\Delta A/\Delta A_{WT}$ at 1625 $cm^{-1}$ is comparable with that expected from removing an adenine nucleobase (12%). The trends in $\Delta A/\Delta A_{WT}$ across AP-site position are similar for CCends and CGCcap. CCends-AP2 has a $\Delta A/\Delta A_{WT}$ of 75% (70%) at 1575 $cm^{-1}$, which is consistent with losing one of its four G:C base pair interactions and suggests that the terminal G:C base pair next to the AP site is mostly intact at 3.8 °C. The AP4 and AP6 sequences only exhibit $\Delta A/\Delta A_{WT}$ values at 1625 $cm^{-1}$ of 90% (82%) and 93% (88%), respectively, again indicating that the loss of signal change is primarily due to a reduction in adenine signal. The value of $\Delta A/\Delta A_{WT}$ at 1575 $cm^{-1}$ is 87% (85%) for CCends-AP4 and 90% (95%) for CCends-AP6, but this is inconsistent with negligible reduction in $\Delta A/\Delta A_{WT}$ at 1662 $cm^{-1}$. The discrepancy could result from different sensitivity of the 1575 $cm^{-1}$ guanine ring mode and 1662 $cm^{-1}$ guanine carbonyl mode to base pairing or stacking.

Unlike CGCcap and CCends, GCGcore only has terminal A:T base pairs which are likely more unstable adjacent to an AP-site than terminal G:C base pairs. For GCGcore-AP2, $\Delta A/\Delta A_{WT}$ at 1625 $cm^{-1}$ is 84% (75%), which may indicate fraying of the terminal A:T base pair next to the AP-site at 3.8 °C. The signatures of low temperature base-pair disruption are clearer for GCGcore-AP4. $\Delta A/\Delta A_{WT}$ at 1625 $cm^{-1}$ is 75% (66%), indicating significant loss of A:T base pairing relative to the WT sequence. This partial dissociation likely comes from fraying of base pairs 1 – 3 on the 5´ side of the AP site that is also observed as an additional melting component (Fig. S36). A reduction of signal change at 1575 and 1662 $cm^{-1}$ is also observed and likely results from the absence of stacking between the adenine and guanine at the $4^{th}$ and $5^{th}$ positions of the duplex.

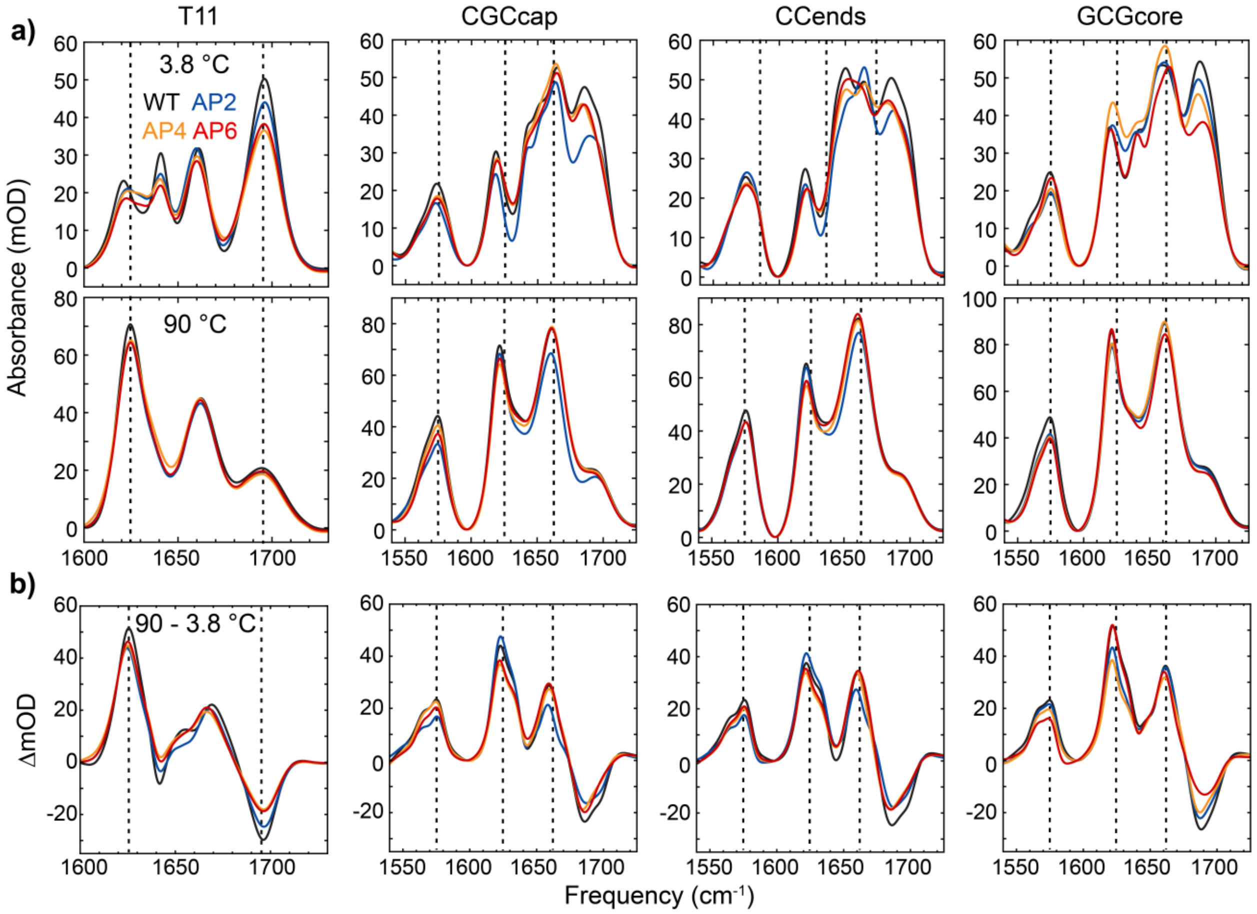


**Figure S33. FTIR spectra of duplex and single-strand states. (a)** Comparison of duplex (3.8 °C) and fully-dissociated oligonucleotides (90 °C) FTIR spectra for each sequence. **(b)** Difference spectra between 90 and 3.8 °C.

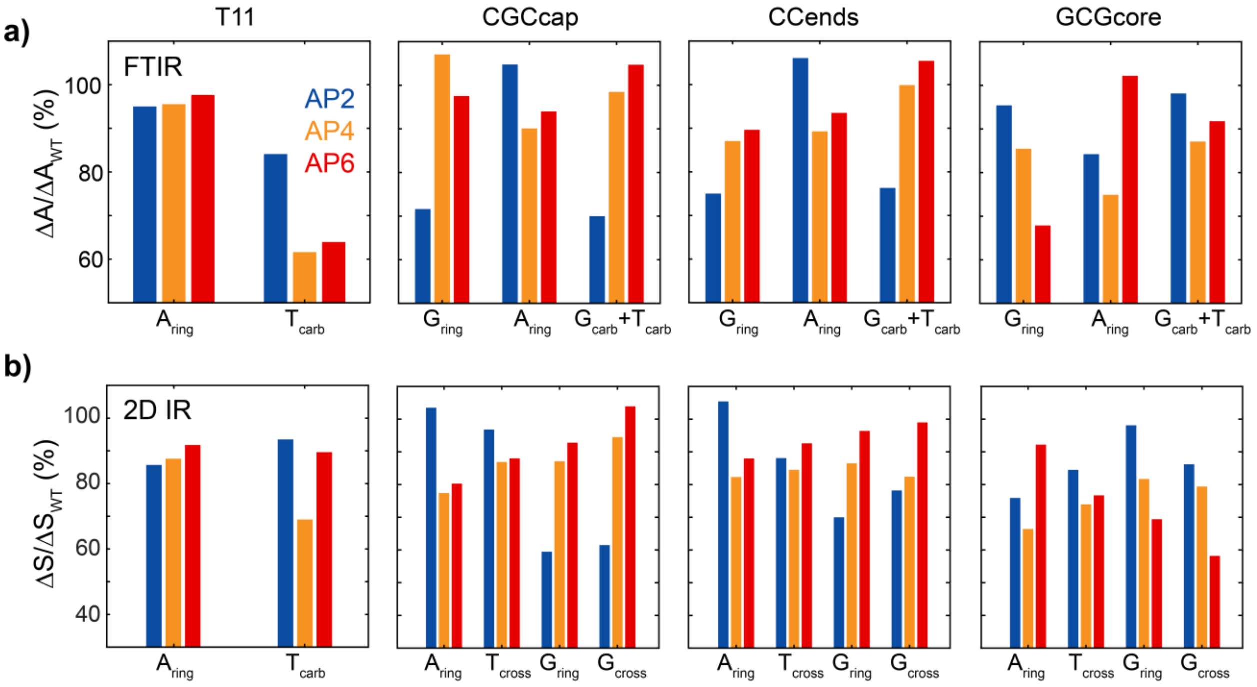


**Figure S34. Total FTIR and 2D IR temperature series spectral change. (a)** Change in FTIR absorbance from 3.8 to 90 °C relative to the WT sequence ($\Delta A/\Delta A_{WT}$) at select frequencies: 1625 $cm^{-1}$ ($A_{ring}$), 1695 $cm^{-1}$ ($T_{carb}$), 1575 $cm^{-1}$ ($G_{ring}$), 1662 $cm^{-1}$ ($G_{carb}$+ $T_{carb}$). **(b)** Change in integrated area of 2D IR spectrum relative to WT sequence ($\Delta S/\Delta S_{WT}$) for given spectral regions. Diagonal regions: 1605 – 1630 $cm^{-1}$ ($A_{ring}$), 1550 – 1590 $cm^{-1}$ ($G_{ring}$), and 1680 – 1700 $cm^{-1}$ ($T_{carb}$). Cross peak regions: 1620 & 1665 $cm^{-1}$ ($T_{cross}$) and 1575 & 1662 $cm^{-1}$ ($G_{cross}$).

#### S6.2 Partial dehybridization from FTIR and 2D IR temperature series

Thermal melting curves extracted from the 2nd SVD component of FTIR temperature series are dominated by a single sigmoidal melting transition (Fig. S2), yet deviations from all-or-nothing melting behavior may be revealed from separately monitoring the temperature-dependence of guanine, adenine, and cytosine vibrational bands. Such information may be extracted from FTIR temperature series. However, each vibrational band possess different low and high temperature linear baselines, particularly the guanine and adenine ring modes, making direct comparison of these regions difficult (Fig. S35). Partial dehybridization at low temperatures is often difficult to distinguish from large amplitude baselines. Previous studies have demonstrated that the relative baseline signal change of carbonyl and ring modes to base-pair disruption is much smaller in pump-probe and 2D IR spectroscopy than in FTIR,(31) presumably stemming from the fourth order scaling of the of the transition dipole moment ( $\mu^4$ ) in the signal of the latter. We make a similar observation for the sequences studied in this work where 2D IR melting curves are dominated by base-pair disruption and low and high temperature baselines are nearly flat (Fig. 6 & S37).

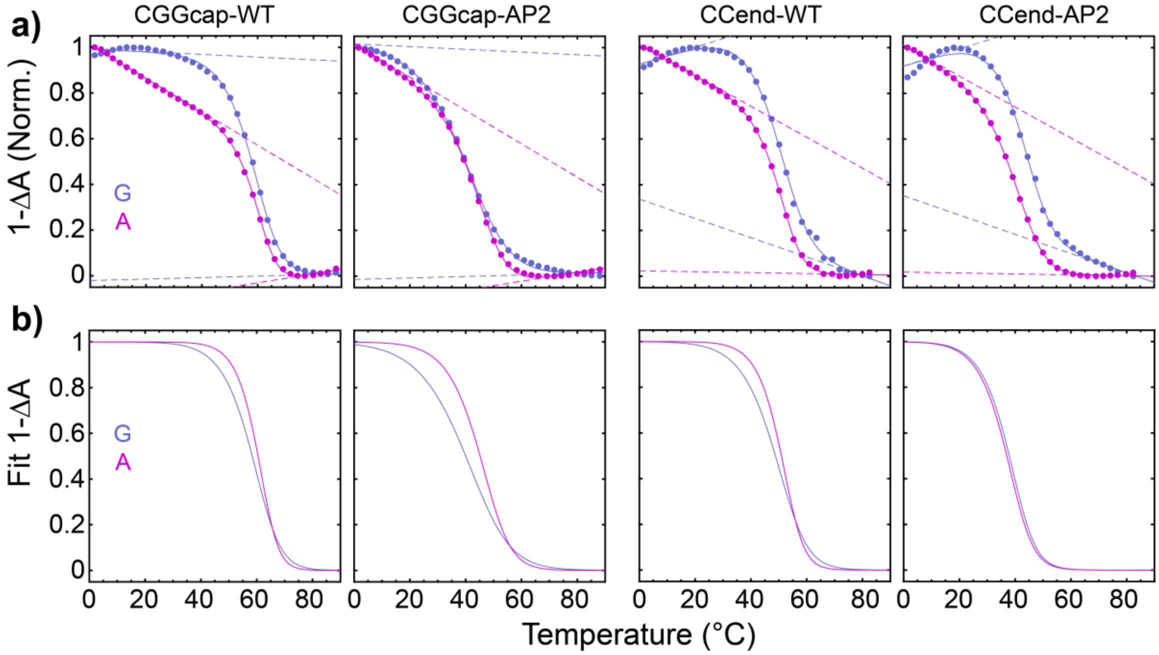


**Figure S35. G:C and A:T melting curves from FTIR. (a)** Normalized change in absorbance at 1575 $cm^{-1}$ (G) and 1625 $cm^{-1}$ (A) for CGCcap and CCend WT and AP2 sequences. Comparison is complicated by the significantly different low-temperature baseline slope observed at G and A bands. Solid lines correspond to two-state fits (eq. S1) using upper and lower baselines (dashed lines). The baseline slopes of the AP2 sequences are fixed to that of the respective WT sequence. **(b)** Fit melting curves at each frequency. The large deviation between G and A melting curves for CGCcap-AP2 and lack of deviation for CCend-AP2 are consistent with the 2D IR melting curves in Fig. 6.

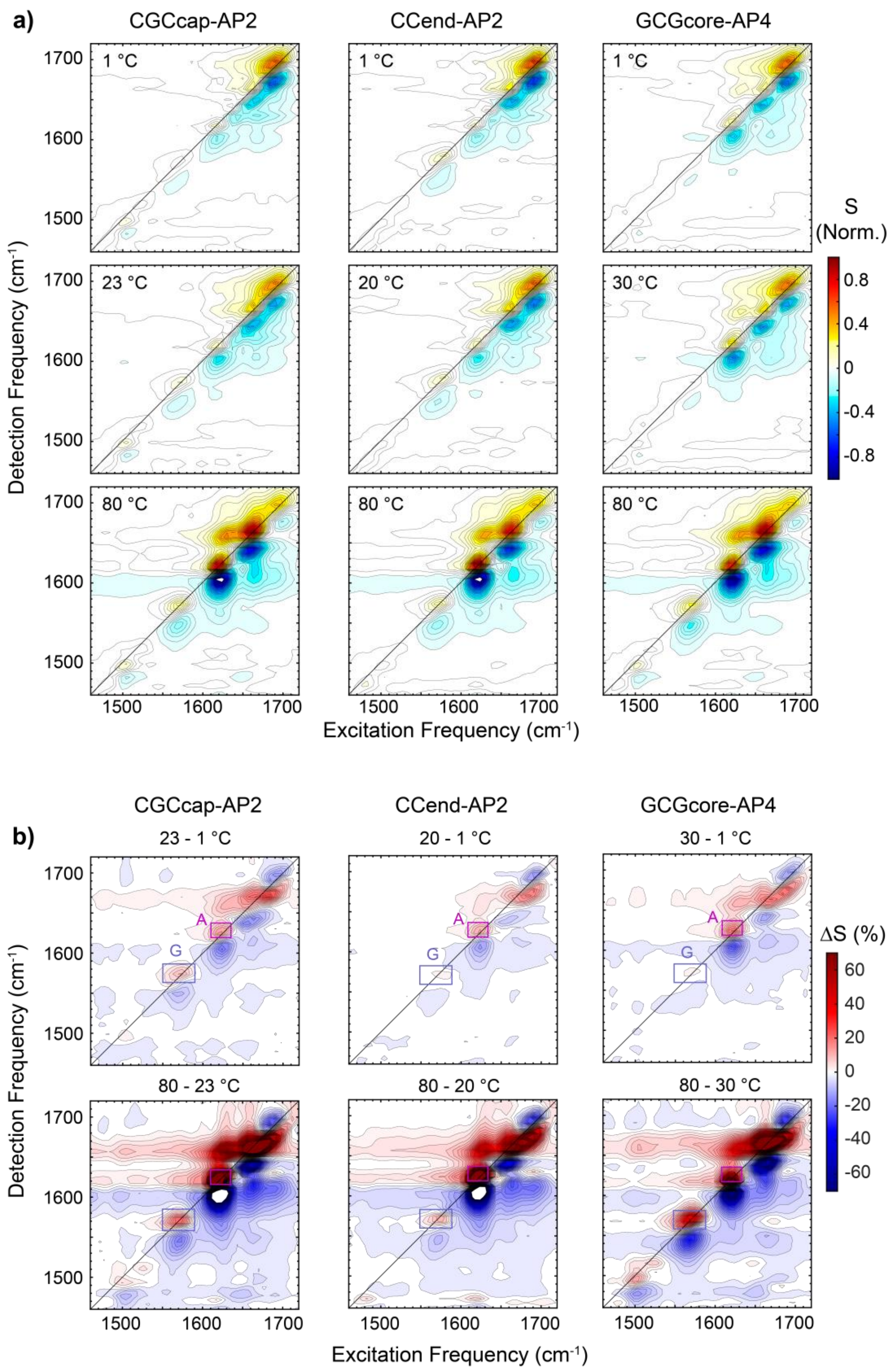


**Figure S36. Partial dehybridization observed with 2D IR. (a)** 2D IR spectra of CGCcap-AP2, CCend-AP2, and GCGcore-AP4 at low (1 °C), intermediate (20 – 30 °C), and high temperature (80 °C).

Spectra are normalized relative to the maximum of the respective 80 °C spectrum. **(b)** 2D IR difference spectra of (top) low-temperature (1 to ~23-30 °C) and (bottom) high-temperature (~23-30 to 80 °C) changes. Spectra are plotted as the difference relative the maximum of the 1 °C, $\Delta S(T) = [S(T_2) - S(T_1)]/S(1°C)$. Contours are plotted with uniform 2.1% spacing. Adenine (A) and guanine (G) ring mode regions are indicated for each sequence.

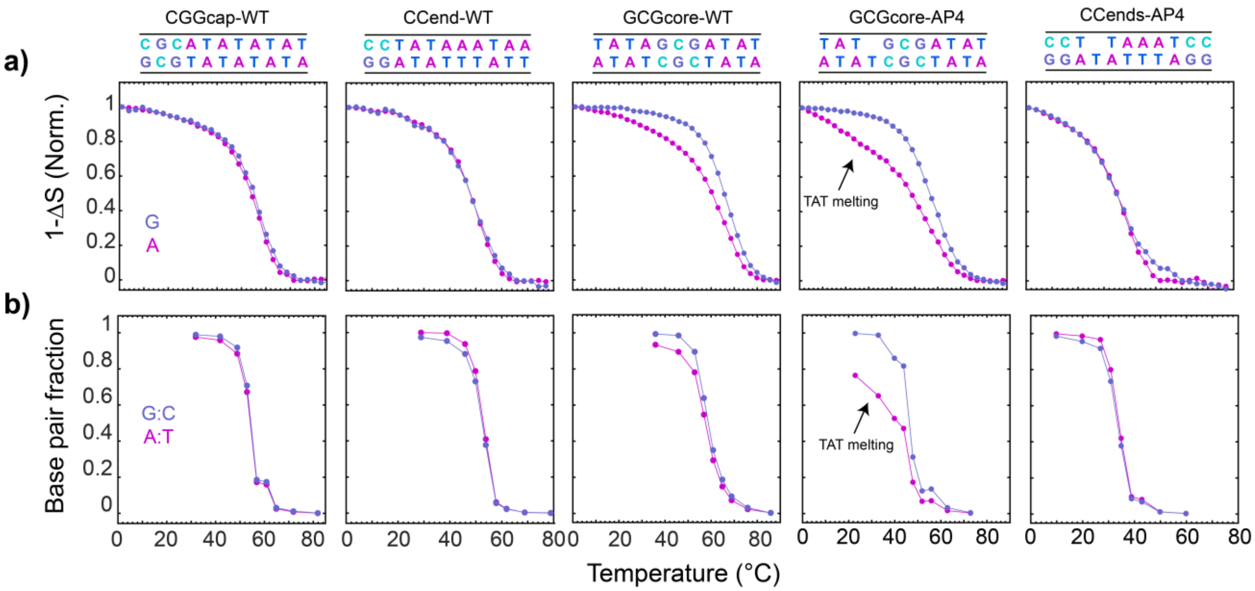


**Figure S37. 2D IR and 3SPN.2 G:C and A:T melting curves for additional sequences. (a)** Normalized temperature-dependent change in 2D IR signals at cytosine (C), guanine (G), and adenine (A) ring vibrational bands for CGCcap-WT, CCend-WT, GCGcore-WT, GCGcore-AP4, and CCends-AP4. **(b)** Fraction of intact G:C and A:T base pairs from temperature-dependent 3SPN.2 MD simulations using a radial base-pair cutoff of 0.7 nm. Experiment and simulation only show significant partial dehybridization in GCGcore-WT (terminal fraying) and GCGcore-AP4 (TAT segment melting and terminal fraying of other end).

##### S6.3 $^1$H NMR spectroscopy of partial dehybridization in AP2 sequences

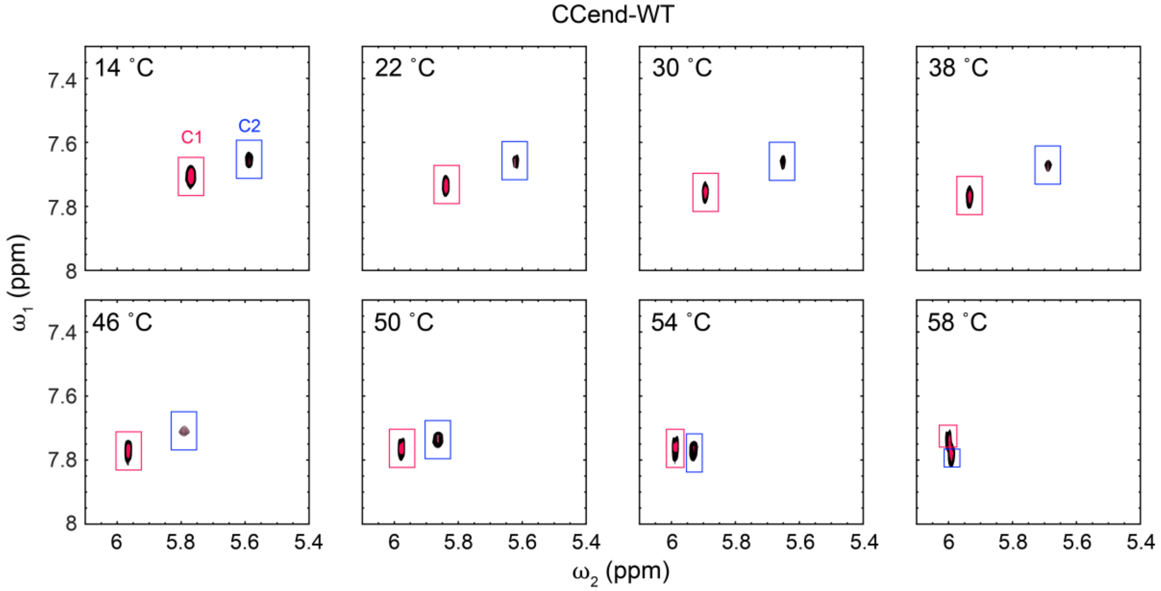


**Figure S38. TOCSY temperature series of CCend-WT.** TOCSY temperature series showing cross peak between H6 and H5 nuclei of C1 (red) and C2 (blue) in CCend-WT. H5 chemical-shift profiles are shown in Fig. 6b.

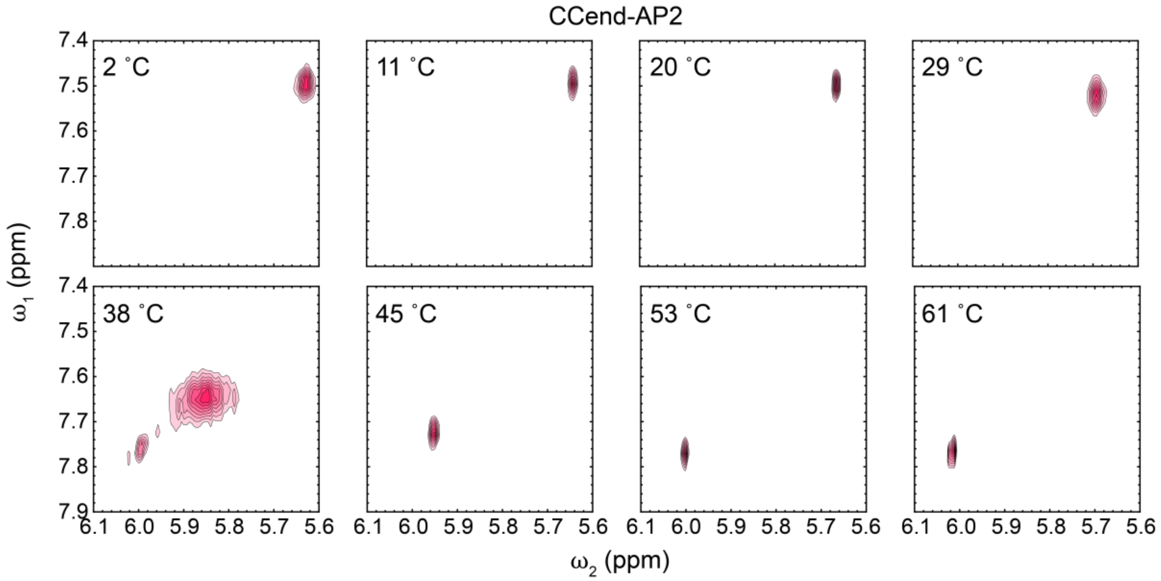


**Figure S39. TOCSY temperature series of CCend-AP2.** TOCSY temperature series showing the cross peak between H6 and H5 nuclei of C1 in CCend-AP2. Significant broadening is observed at 38 °C, which is near the duplex melting temperature. The H5 chemical-shift temperature profile is shown in Fig. 6b.

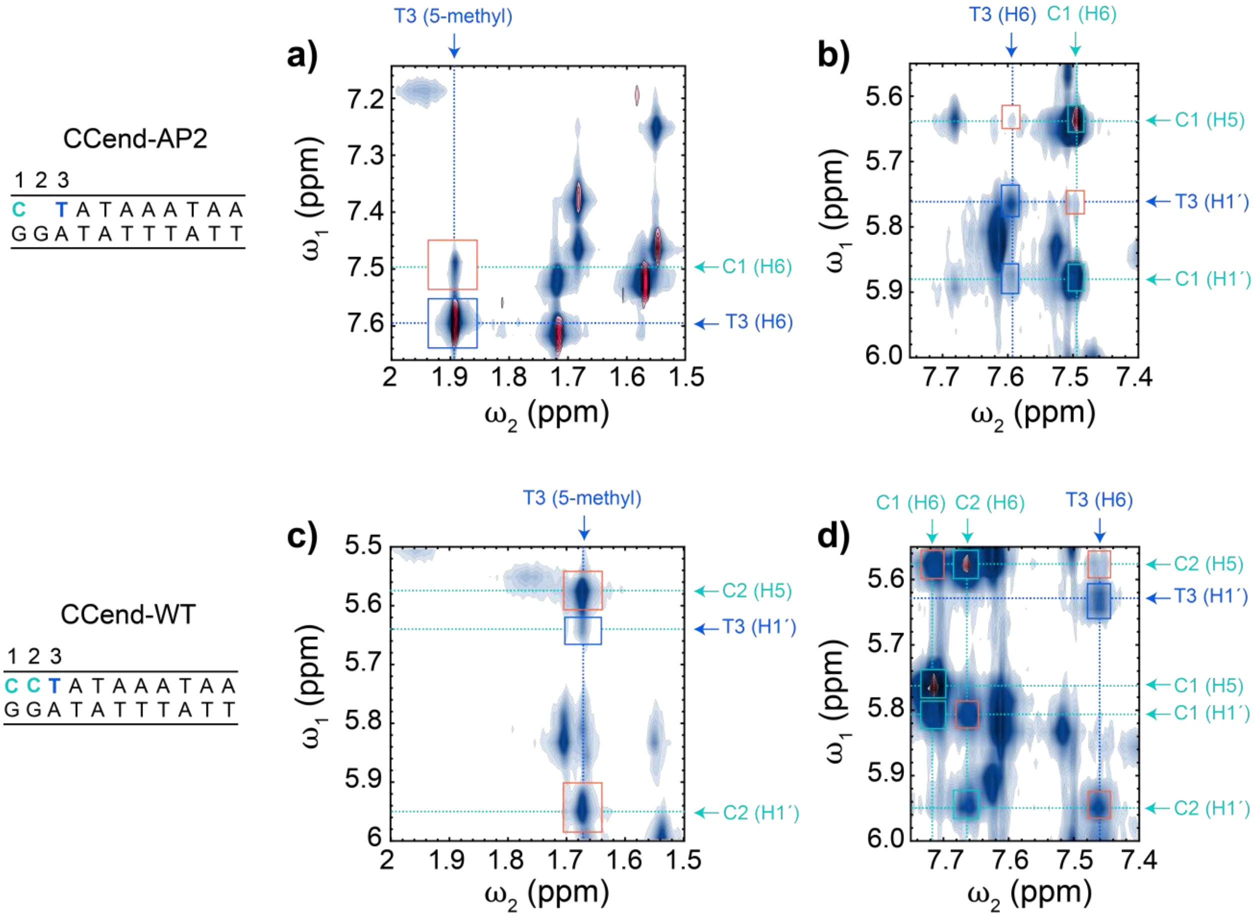


**Figure S40. ¹H-¹H cross-peaks in CCend-WT and CCend-AP2. (a)** NOESY cross-peaks (blue contours) between H6 nuclei (7.15 – 7.65 ppm) and 5-methyl protons of thymine (1.5 – 2.0 ppm) of CCend-AP2 in deuterated solution at 15 °C. The TOCSY spectrum is overlaid in red contours. NOESY and TOCSY spectra were measured at mixing times of 200 ms and 20 ms, respectively. **(b)** NOESY and TOCSY cross peaks between H6 and H5/H1´ (5.55 – 6.0 ppm). Cross-peaks **(c)** between H5/H1′ nuclei and 5-methyl protons and **(d)** between H6 and H5/H1′ are shown for CCend-WT. Orange boxes denote interbase cross-peaks while blue and cyan boxes correspond to intrabase thymine and cytosine cross-peaks, respectively. Numerous cross-peaks between C1 and T3 in CCend-AP2 support the assignment in Fig. 6. In CCend-WT, cross-peaks are instead observed between C2 and T3.

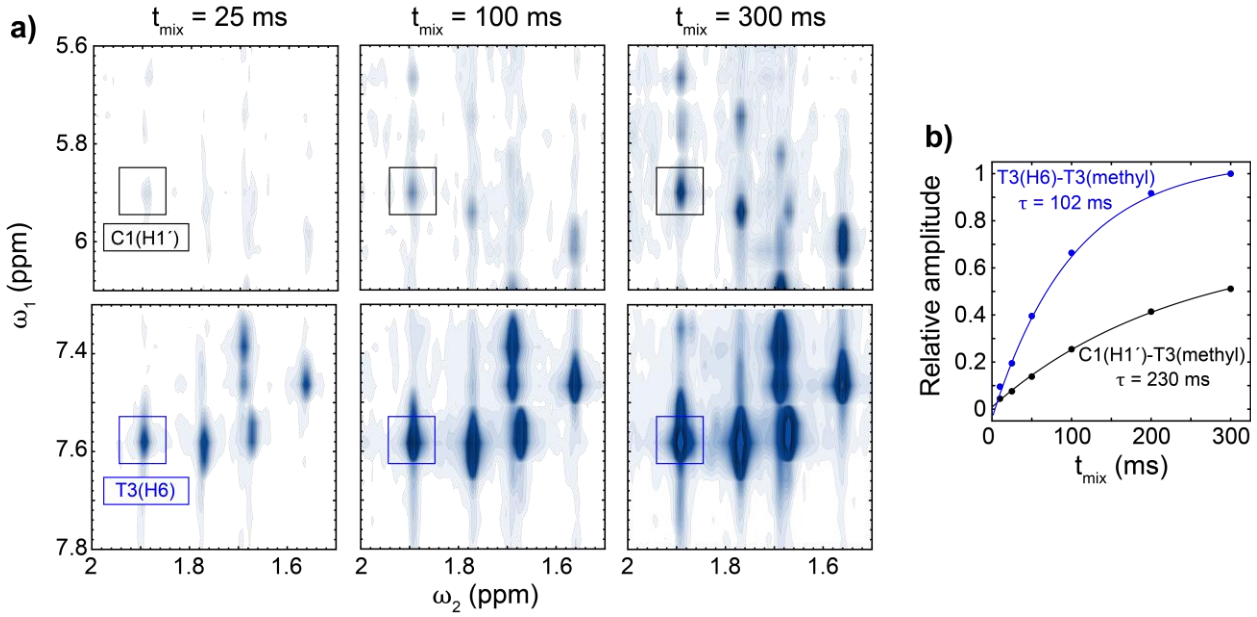


**Figure S41. NOESY mixing time series of CCend-AP2. (a)** NOESY spectra of CCend-AP2 using $t_{mix}$ = 25, 100, and 300 ms at 15 °C highlighting the cross-peaks between aromatic protons of C1 and T3 with the 5-methyl protons of T3. **(b)** Integrated amplitude of T3(H6) to T3(methyl) and C1(H1′)-T3(methyl) cross-peaks as a function of $t_{mix}$. Each trend is fit to a single-exponential rise with the reported time constant ($\tau$). The exponential dependence suggests that each cross-peak arises from direct dipolar coupling instead of indirect coupling through spin diffusion.

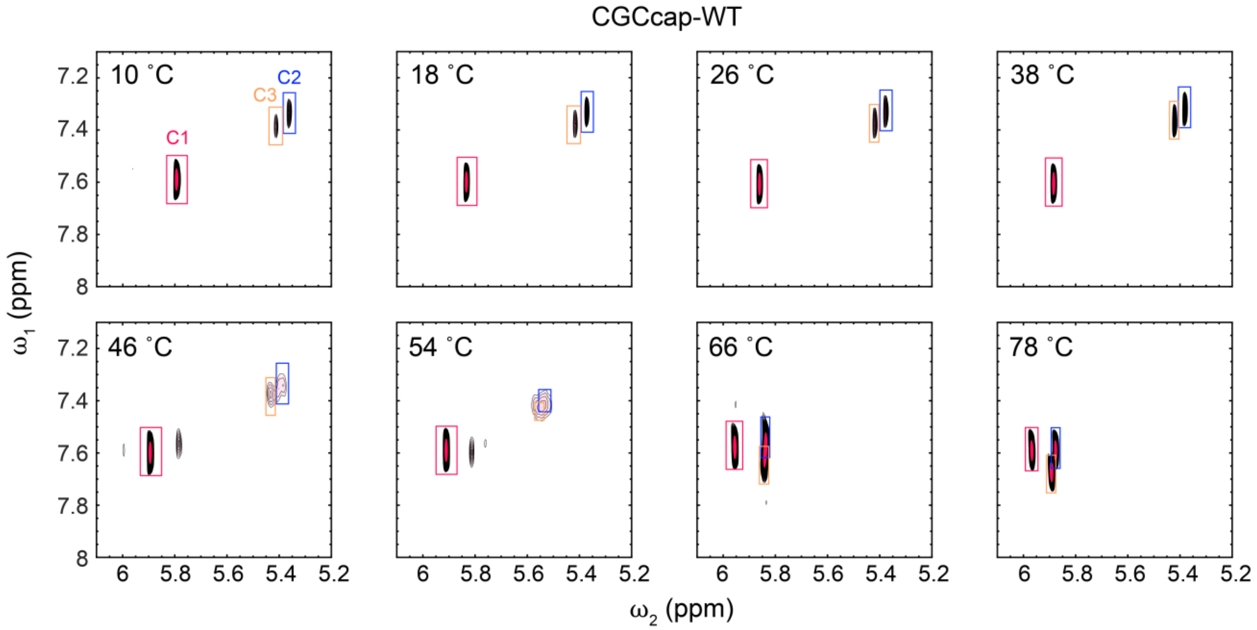


**Figure S42. TOCSY temperature series of CGCcap-WT.** TOCSY temperature series showing cross peaks between H6 and H5 nuclei for C1 (red), C2 (blue), and C3 (yellow) bases in CGCcap-WT. H5 chemical-shift profiles are shown in Fig. S44.

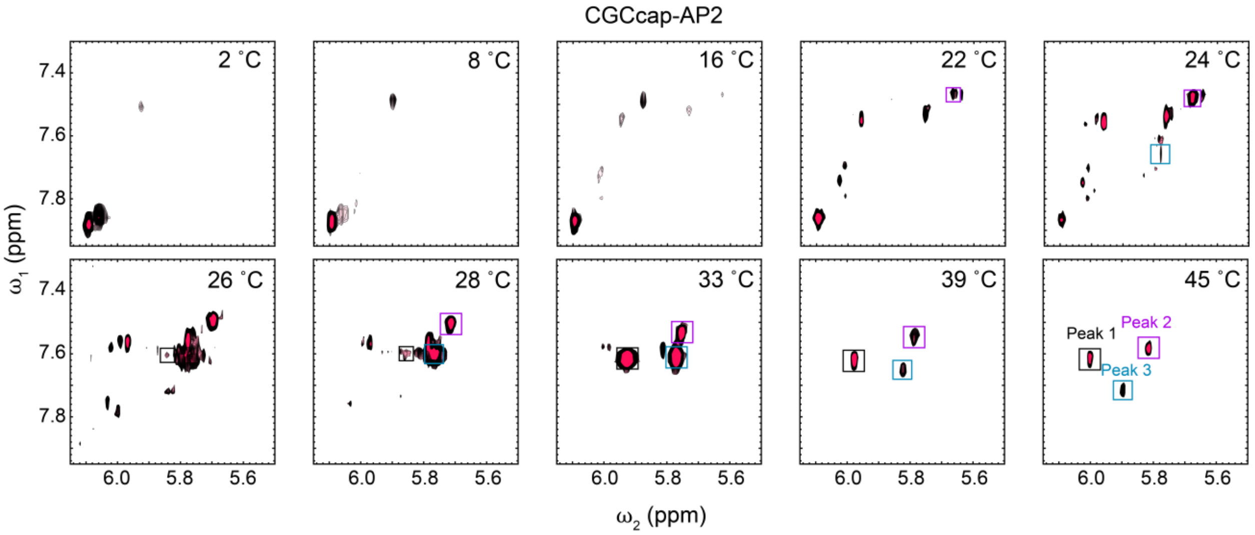


**Figure S43. TOCSY temperature series of CGCcap-AP2.** TOCSY temperature series showing cross peaks between H6 and H5 nuclei of cytosine bases in CGCcap-AP2. The most intense peaks from 2 to 16 °C occur in the 5.9 to 6.2 ppm H5 frequency region. This is followed by a gain of in intensity of peaks at lower chemical shift (5.6 – 5.8 ppm) from 16 to 28 °C with a corresponding loss of the peaks in the 5.9 – 6.2 ppm region. The peaks in the 5.6 – 5.8 ppm region move to higher chemical shift due to base-pair disruption as the temperature increases. The three cross peaks observed in the single-strand state (Peak 1, Peak 2, and Peak 3) are marked when observable.

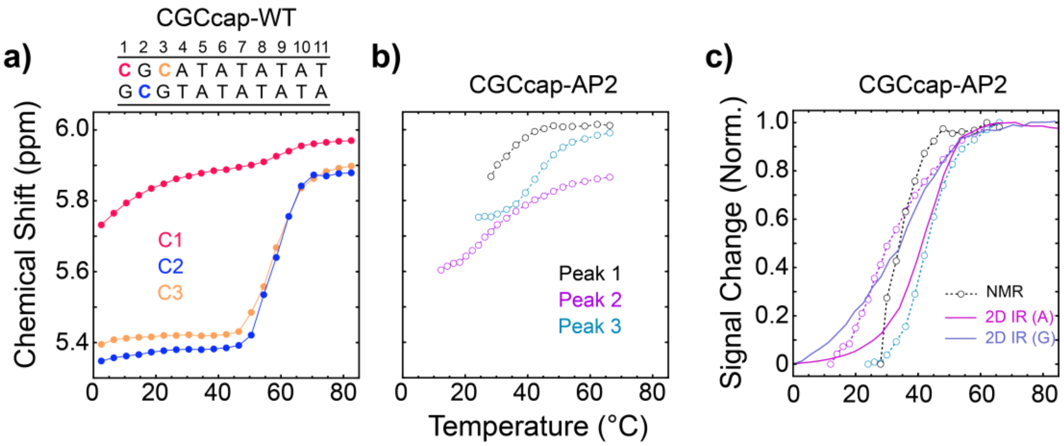


**Figure S44. Temperature trends of cytosine H5 nuclei in CGCcap sequences from TOCSY measurements. (a)** Chemical shifts of cytosine H5 nuclei for CGCcap-WT determined from TOCSY spectra shown in Fig. S41. Assignments to C1, C2, and C3 nucleobases were made using $^{1}H$-$^{1}H$ NOESY spectra (not shown). **(b)** Chemical shift temperature-trends of the three remaining cytosine H5 resonances in CGCcap-AP2 at high temperature. Peaks are marked with color-coded boxes in Fig. S43. **(c)** Normalized trends in **(b)** compared with G:C and A:T melting profiles from 2D IR (solid lines). The melting transition observed in Peak 3 from NMR is >10 °C higher than in Peak 1 and 2 and roughly matches the A:T melting profile from 2D IR. The Peak 1 and 2 transitions are not fully resolved but appear to overlap with partial dehybridization of G:C base pairs from 2D IR.

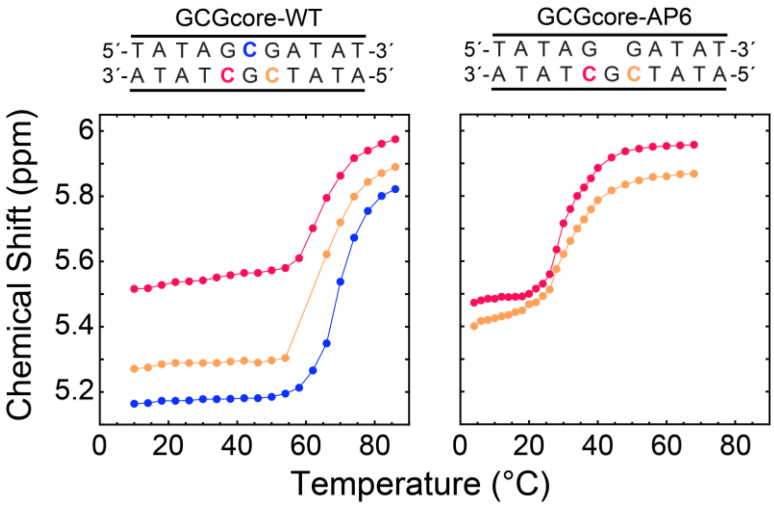


**Figure S45. Temperature-dependent cytosine $^1$H signals in GCGcore-WT and GCGcore-AP6.** H5 signals of (left) GCGcore-WT and (right) GCGcore-AP6 determined from TOCSY spectra at a mixing time of 20 ms. H5 nuclei are color-coded according to their position using assignments made from NOESY spectra (not shown). For each sequence, all H5 signals follow the same single melting transition.

#### S6.4 All-atom MD simulations of AP2 sequences

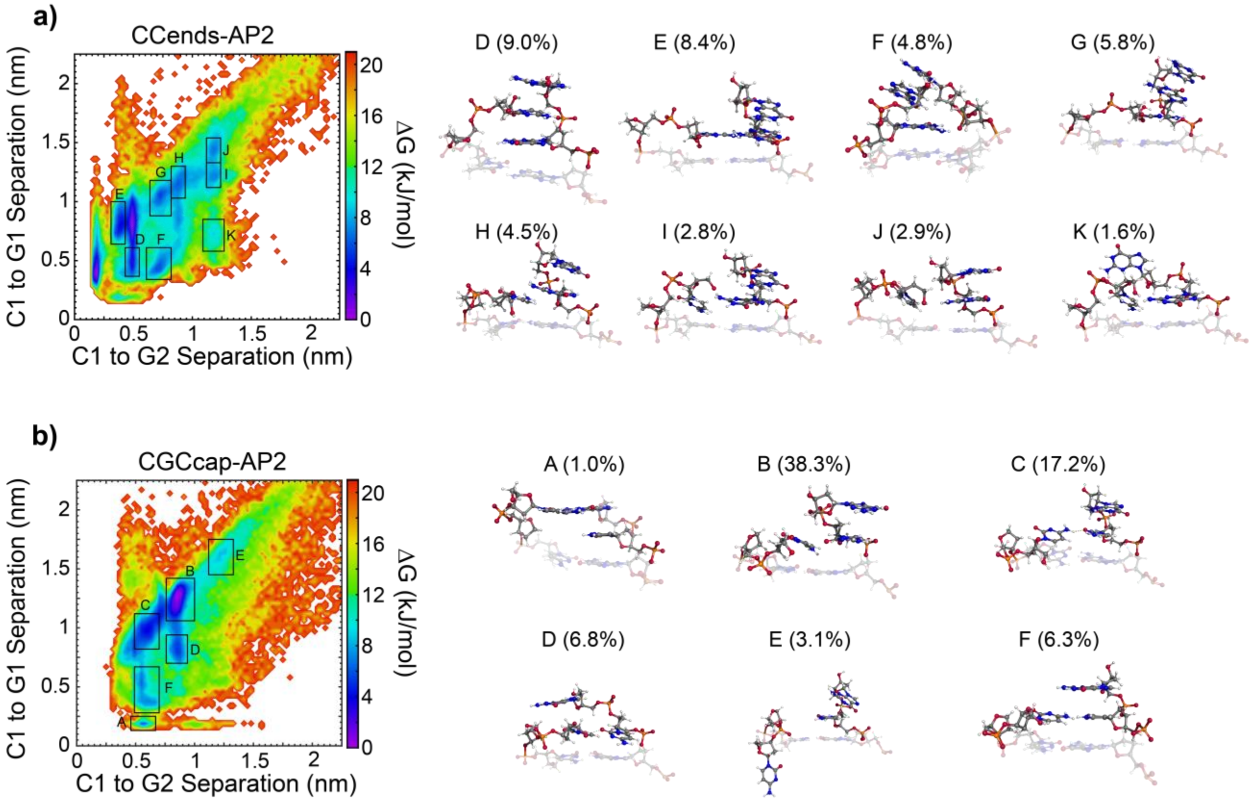

**Figure S46. Free-energy surfaces for terminal base pairing in CCends-AP2 and CGCcap-AP2.** Free-energy surfaces generated from 20 × 1 μs MD simulations with AMBER-bsc1 for truncated **(a)** CCends-AP2 and **(b)** CGCcap-AP2 duplexes. The x-axis is the average separation between hydrogen-bonding atoms between C1 and G2 (out-of-register), and the y-axis is the average separation between those atoms in C1 and G1 (in-register). Snapshots of base pairs 1-3 from time points sampling the marked local free-energy minima are shown on the right. Percentages correspond to the integrated population within each rectangle. Simulations are initialized with in-register base pairing (state A), but each sequence prefers to adopt a variety of out-of-register configurations where the C1 nucleobase stacks with T3 (CCends) or C3 (CGCcap) or intercalates into the opposing strand.

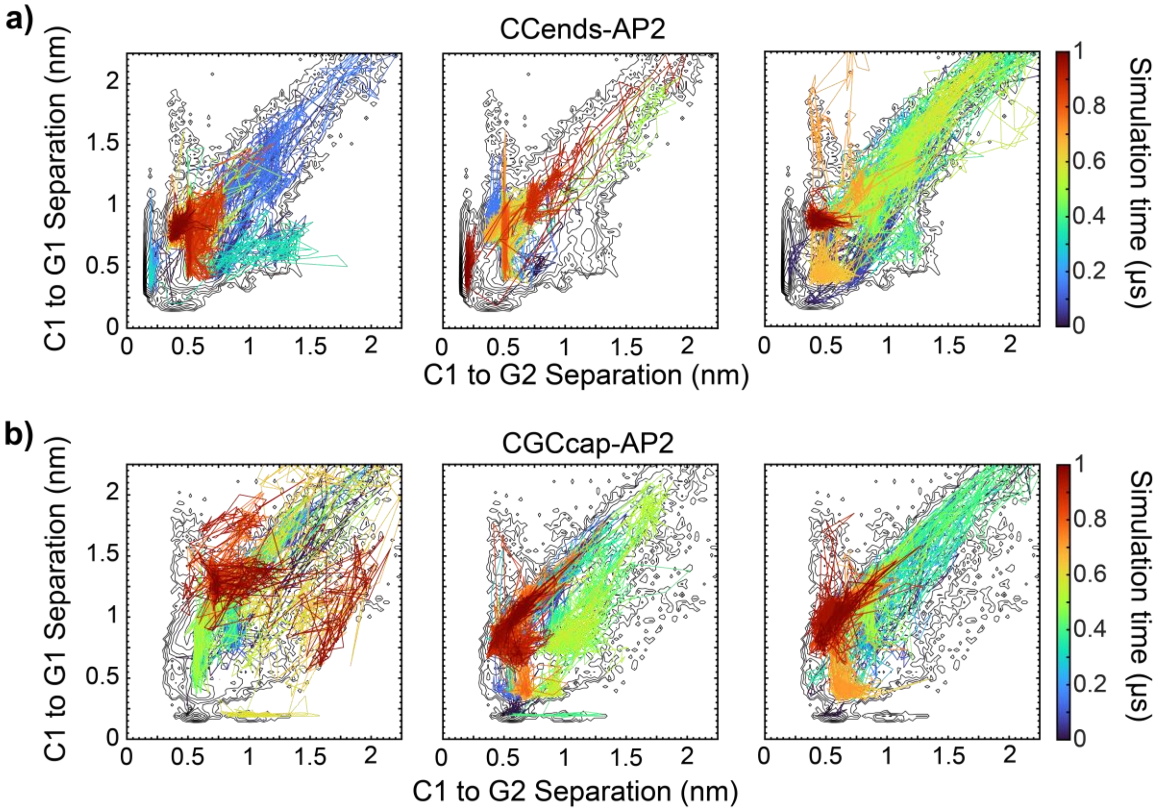


**Figure S47. Example all-atom trajectories for terminal base pairing in CCends-AP2 and CGCcap-AP2.** Select 1 μs trajectories for **(a)** CCends-AP2 and **(b)** CGCcap-AP2 are shown as lines color-coded based on simulation time. The free energy surface is shown as black contours. Each trajectory samples most of the free energy minima, illustrating that the different base-pairing configurations shown in Figs. 6d and S46 interconvert on sub-microsecond timescales.
